# High-throughput physics-based enzyme engineering

**DOI:** 10.64898/2026.09.15.751901

**Authors:** Xujian Wang, Xiang Qiu, Yuyang Wu, Taoyu Niu, Shuhao Zhang, Runtian Gao, Ilkwon Cho, Haocheng Tang, Kangdelong Hu, Xiaoguang Lei, Olexandr Isayev, Junmei Wang

## Abstract

Enzyme engineering aims to tailor natural enzymes for industrial and therapeutic applications, yet physically grounded rational design has been limited by a trade-off between accuracy and cost, leaving the field heavily dependent on expert intuition. Here we present a scalable physics-based framework that combines field-aware machine learning with molecular mechanics to capture enzyme electrostatics at quantum-mechanical accuracy while enabling efficient, atomistic exploration of reaction free-energy landscapes. Coupled with microkinetic modelling, the framework translates molecular free-energy landscapes into catalytic rates and selectivity across competing, multistep reaction pathways. Applied to a newly engineered oxidative amidase (OxiAm), the framework predicts catalytic rate constants with near-experimental accuracy, quantitatively resolves the selectivity between hydrolysis and aminolysis, and generalizes across substrates, mutations and enzyme homologues. Transition-state ensemble analysis further reveals the reaction mechanism and guides the design of enzyme variants for pharmaceutical synthesis. By bringing chemical accuracy and high-throughput sampling to enzyme catalysis, this approach shifts rational design from static, empirical practice toward dynamic, free-energy-driven design, and should accelerate the engineering of biocatalysts.

## 1 Introduction

Enzymes are nature’s catalysts, achieving remarkable rate accelerations and selectivity under physiologically relevant conditions, and they play central roles in biosynthesis, pharmaceutical manufacturing and a broad range of industries. Such enzymes have traditionally been tailored through directed evolution,^1^ a powerful but labor- and resource-intensive process. To reduce this burden, computational and artificial intelligence (AI)-based approaches have emerged, ranging from data-driven, AI-based de novo enzyme design ^2–7^ to protein language model-based function annotation^8–12^ and kinetic-parameter prediction. ^13–16^ Comparatively underdeveloped, however, is enzyme design at the atomistic scale, grounded in physically interpretable free-energy principles. This is the domain of rational design, which rests on a rigorous physical foundation to harness the catalytic capabilities that nature has refined over millions of years of evolution. The capacity to perform such rational design at will, tailoring these natural catalysts to the full range of industrial and therapeutic applications, would be transformative. ^17^

Realizing this capacity has long been constrained by the cost of accurate simulation. Even quantum mechanics/molecular mechanics (QM/MM), which combines quantum-level accuracy with the efficiency of a classical environment,^18,19^ remain applicable only a limited range of systems, leaving mechanistic understanding heavily dependent on human expert intuition drawn from sparse calculations. The advent of machine-learning/molecular-mechanics^20–24^ (ML/MM), in which machine-learning interatomic potentials (MLIPs) describe the reactive region, has begun to transform this picture: it delivers quantum-mechanical accuracy with several orders of magnitude of acceleration, placing high-throughput, atomistic free-energy computation within easy reach. Because every component of the framework is physically grounded and interpretable, it substantially lessens the reliance on practical experience; and because it is rooted in physics, it transfers without retraining to arbitrary chemical reactions, protein families and substrate scopes, resolving subtle mutational effects.

In this study, we assembled a dataset that captures the electrostatic environment of biomolecules at quantum-mechanical accuracy, and used it to develop MACEPOL-EF, a MLIP that is explicitly conditioned on the surrounding electrostatic environment, representing a significant advance toward a fully coherent treatment of hybrid ML/MM potentials for studying complex systems. We then implemented an electrostatic-embedding ML/MM framework in AMBER^25,26^ and, coupling it with enhanced sampling, explored the free-energy landscapes of enzymatic reactions. We further introduced a microkinetic modelling scheme that faithfully reproduces the activity and selectivity of competing reactions from experiments. Applied across diverse reactions, mutations, substrates and protein families, this integrated approach proved broadly generalizable. Building on the enzymes that nature has refined over millions of years, our method offers a new philosophy for rational design and extends the boundaries of natural enzymes in a more physical and principled way.

To test this framework on a real-world problem, we selected the oxidative amidase (OxiAm) as a model system. OxiAm was recently created by engineering an aldehyde dehydrogenase (ALDH) to couple aldehydes with amines into amide bonds, an activity not found in nature.^27^ The fact that this activity can be installed simply by redirecting the enzyme’s native hydrolytic machinery toward aminolysis illustrates the latent potential of naturally evolved scaffolds, reinforcing our central premise: a wealth of existing enzymes remains to be tailored, and a physically grounded, generalizable framework is well positioned to unlock them. OxiAm is a particularly demanding testbed for this purpose: its mechanism remains unresolved, and the reaction proceeds through multiple elementary steps and two competing pathways whose closely spaced barriers blur the rate-determining step. Importantly, this study aims much less to characterize a single enzyme than to demonstrate a shift in the philosophy of rational design: from static single structures toward an approach that is dynamic, free-energy-driven, high-throughput and non-empirical.

## 2 Results

### 2.1 An electric field-aware machine-learning interatomic potential

MLIPs reproduce quantum-mechanical accuracy at a fraction of the cost,^28,29^ but deploying them inside biological macromolecules demands a coherent treatment of the hybrid ML/MM potential, in which the ML-described region must be explicitly conditioned on the surrounding electrostatic environment.^30–32^ Accounting for this environmental dependence remains a persistent challenge and a major source of error in ML/MM.^21^

To address this obstacle, we performed molecular dynamics simulations on 15 well-characterized protein systems relevant to drug discovery, encompassing more than 600 protein-ligand complexes, and quantified the electric fields experienced by individual residues and substrates (Extended Data Fig. 1a,b). These fields vary systematically with residue type and local environment, spanning approximately 0.49–1.60 V ^*−*1^ (Fig. 1b), in close agreement with vibrational Stark-effect spectroscopy.^33^ Such field strengths far exceed the range covered by the current models^34^(Fig. 1c), highlighting the need for dedicated training data to extend MLIPs into this regime. Guided by these distributions, we developed an electric field-informed strategy that reconstructs external fields matching these element-dependent statistics around molecular structures (Extended Data Figs. 1c and 2a,b) and labels the resulting configurations with density functional theory (DFT), yielding the BioPol-EF dataset (Fig. 1a; see Methods).

**Figure 1.**
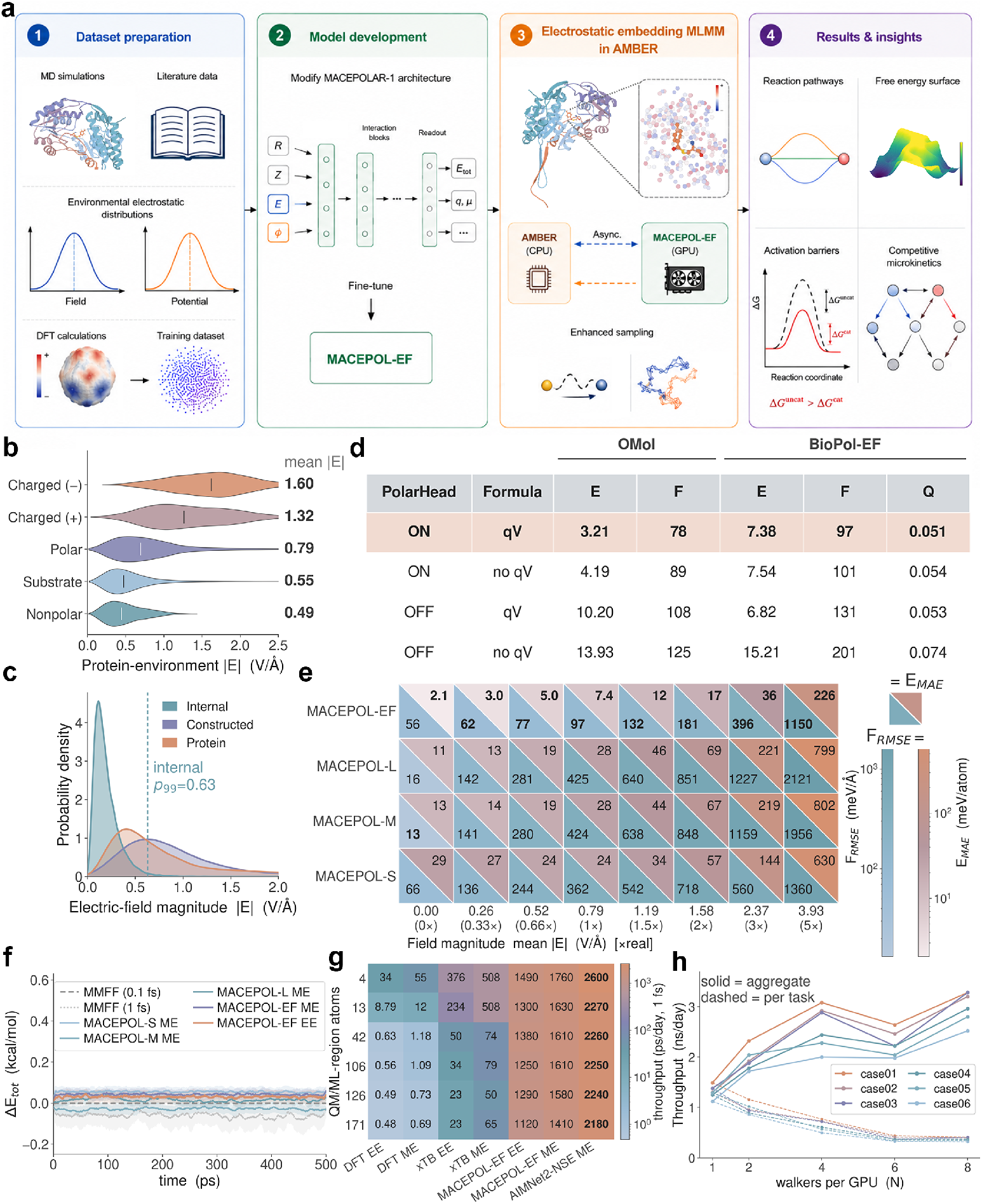
An electrostatic-embedding ML/MM framework for atomistic, free-energy-resolved enzyme catalysis. (**a**) End-to-end workflow: electric field informed dataset preparation, fine-tuning of the field-aware potential MACEPOL-EF, electrostatic-embedding ML/MM implementation in AMBER, and free-energy computation and analysis of enzymatic reactions. (**b**) Distributions of the external electric field acting on different groups, extracted from molecular-dynamics simulations. (**c**) The intramolecular (“internal”) field natively sampled by the base model (99th percentile, 0.63) compared with the true protein-derived external field and with the field reproduced by our constructed dataset. (**d**) Contribution of the two architectural additions, the adaptive polar head and the per-atom charge–potential (*qV*) coupling, across all four combinations, evaluated on held-out OMol and BioPol-EF validation sets; *E*, energy MAE; *F*, force RMSE; *Q*, charge error. (**e**) Accuracy on an out-of-distribution graded-field test set: taking the mean protein field from MD (0.79) as the unit (1×), external fields from 0× to 5× are applied and evaluated for four MACEPOL model sizes (EF, L, M, S); upper triangles, energy MAE; lower triangles, force RMSE. (**f**) Total-energy drift (Δ*E*_tot_) over 500 ps of NVE dynamics; molecular mechanics force field (MMFF) is integrated at 0.1 and 1 fs and all ML/MM variants at 0.1 fs, under mechanical (ME) and electrostatic (EE) embedding. (**g**) Throughput as a function of the number of QM/ML-region atoms for DFT, xTB, MACEPOL-EF and AIMNet2-NSE under EE and ME. (**h**) CPU–GPU load balancing for MACEPOL-EF (EE) at a fixed 16 CPU cores and 1 GPU: aggregate (solid) and per-task (dashed) throughput versus the number of walkers per GPU. Electric fields are in V Å^*−*1^; energy errors (*E*) in meV atom^*−*1^, force errors (*F*) in meV Å^*−*1^and charge errors (*Q*) in *e*; throughput is in ps day^*−*1^(**g**) and ns day^*−*1^(**h**).

Then, we extended the MACEPOLAR-1 architecture^34^ in two places (Extended Data Fig. 3). A charge–potential (*qV*) coupling lets each atom sense the anisotropic external field and potential, while an adaptive polar head captures the linear polarisation response together with higher-order contributions (Fig. 1d and Extended Data Fig. 2c). Fine-tuning on BioPol-EF yielded MACEPOL-EF, a foundation MLIP that is field-aware at atomic resolution and predicts physically meaningful charges. On an out-of-distribution graded-field test set its prediction error grows far more slowly with field strength than that of the field-agnostic base models, and it retains chemical accuracy to ~1.5 V Å^*−*1^ (Fig. 1e), covering the full range of fields found in the active sites studied here.

### 2.2 Electrostatic embedding ML/MM framework

To couple MACEPOL-EF with MM, we adopted an electrostatic-embedding scheme in which the ML and MM regions mutually respond to each other’s electrostatics: the ML atoms respond to the external electric field and potential generated by the surrounding MM atoms, while the MM atoms respond to the on-the-fly electrostatics of the ML region.^24,35,36^ This bidirectional coupling provides a physically consistent description of the embedded system, reflected in excellent energy conservation during molecular dynamics (Fig. 1f). MACEPOLEF also enables concurrent CPU-GPU computing (Fig. 1a), achieving speedups of up to several thousand-fold over DFT and semi-empirical quantum methods, with substantially better scaling with system size (Fig. 1g and Extended Data Fig. 2d,e). Optimization of the CPU-GPU workload balance further improves computational efficiency (Fig. 1h), while inference-level kernel and quantization optimizations can be applied to the trained weights without retraining, yielding additional efficiency gain at negligible cost to accuracy (Extended Data Fig. 2f,g). Together, these features provide an accurate, efficient and physically grounded framework for high-throughput simulation of enzymatic catalysis.

### 2.3 Reaction mechanism of the engineered ALDH

Protein engineering converts natural ALDHs into OxiAms that transform an aldehyde into either an acid or an amide^27^ (Fig. 2a). Previous structural studies^37,38^ identified NAD(P)^+^ binding in the left-hand tunnel of the active site (Extended Data Fig. 4a–g), while mutagenesis^27^and our enhanced sampling identified the key residues lining the catalytic pocket, where both the aldehyde and the amine enter and react (Fig. 2b and Extended Data Fig. 4a). We then explored the reaction with our electrostatic-embedding ML/MM framework (Fig. 2c) combined with OPES-MetaD enhanced sampling.^39^

**Figure 2.**
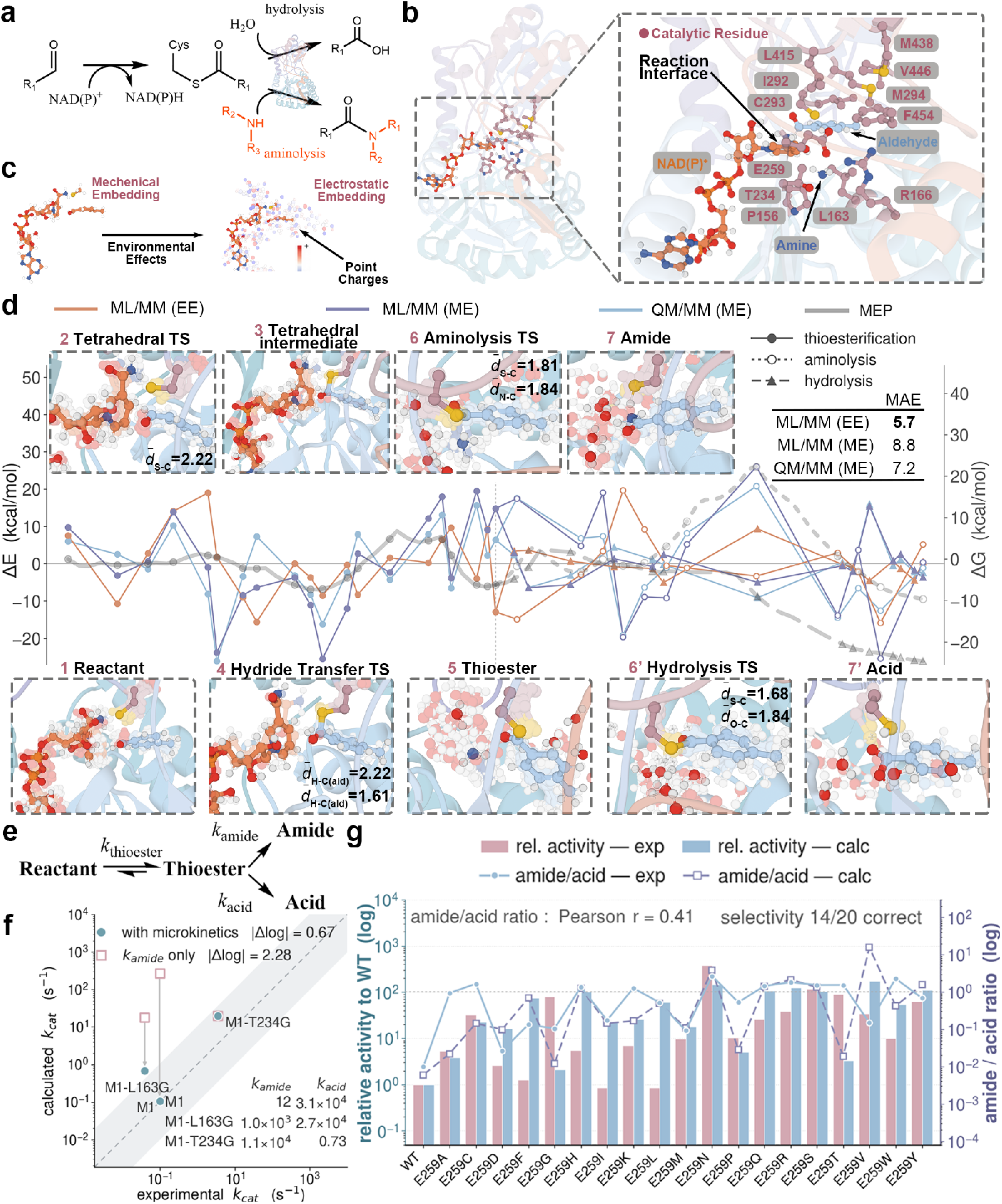
Reaction mechanism and microkinetic modelling of OxiAm catalysis. (**a**) OxiAm first oxidizes an aldehyde to a common thioester intermediate, which then partitions between two competing branches, hydrolysis and aminolysis, yielding the acid and amide products, respectively. (**b**) Overall structure and enlarged active site, showing the relative binding positions of NAD(P)^+^, the aldehyde and the amine, and the residues that form the catalytic interface. (**c**) Mechanical (ME) and electrostatic (EE) embedding representations of the reactive region; under EE, the machine-learning atoms respond to the surrounding environment, represented as point charges. (**d**) Energetics of the full mechanism across its three reaction segments (thioesterification, aminolysis and hydrolysis). The minimum-energy pathway (MEP) is extracted from the simulated free-energy surface; along it, the change in energy (Δ*E*) is evaluated by three methods, including QM/MM (ME), ML/MM (ME) and ML/MM (EE), with electrostatic-embedding QM/MM (QM/MM (EE)) as the reference (inset, mean absolute error, MAE). Filled circles, open circles and triangles denote thioesterification, aminolysis and hydrolysis, respectively. For representative reactants, intermediates and transition states (labelled 1–7 and 6^*′*^–7^*′*^), the conformational ensembles are extracted and key geometric parameters (mean interatomic distances 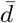, in Å) are annotated. (**e**) Reduced microkinetic network defined by *k*_thioester_, *k*_amide_ and *k*_acid_. (**f**) Calculated versus experimental *k*_cat_ using the full microkinetic model or *k*_amide_ alone; the shaded band indicates agreement within one order of magnitude. (**g**) Experimental and calculated relative activities and amide/acid ratios for the position-259 saturation-mutagenesis panel.

From the resulting free-energy surface we extracted the minimum-energy pathway (MEP) and evaluated representative structures along it under both electrostatic (EE) and mechanical (ME) embedding schemes (Fig. 2d). Electrostatic embedding markedly accelerated the free-energy convergence, and the selected ML region reliably yielded converged free-energy barriers (Extended Data Fig. 5a,b). Our method faithfully reproduced the QM/MM (EE) atomic charges (Extended Data Fig. 6a) and, using QM/MM (EE) as the reference, achieved the lowest energy and force errors among the methods tested (Fig. 2d and Extended Data Fig. 6b,c). The framework therefore retains quantum-level accuracy across the entire multistep pathway.

The validated pathway defines the catalytic mechanism (Fig. 2d). Once the aldehyde and NAD(P)^+^ are bound, Cys293, in its deprotonated thiolate form, attacks the aldehyde carbonyl carbon to form a tetrahedral intermediate; subsequent hydride transfer to NAD(P)^+^ yields NAD(P)H and a metastable thioester. This thioester undergoes one of two competing pathways, hydrolysis and aminolysis, in which water or an amine, respectively, attacks the thioester carbonyl. In both pathways, proton transfer is facilitated by a preorganized, water-mediated hydrogen-bond network, ultimately yielding the acid and amide products (Fig. 2d). Because configurations around each point along the MEP were sampled as an ensemble, we found substantial heterogeneity among the elementary steps: tetrahedral formation and hydride transfer adopt rigid and preorganized configurations, whereas aminolysis and hydrolysis populate broad, heterogeneous transition-state ensembles in which a longer nucleophile–electrophile distance is compensated by a better attack angle (Fig. 2d). Ensemble-level analysis is therefore required, as a single static structure would obscure the conformational diversity needed to accurately describe reactivity.

### 2.4 Microkinetic modelling of enzyme catalysis

Microkinetic modelling neither presupposes a rate-determining step (RDS) nor restricts the reaction to a single route, ^40,41^ making it well suited to OxiAms, in which thioester formation, hydrolysis and aminolysis have comparable barriers that vary in a coordinated manner upon mutation: no single step dominates the overall rate, and the two competing branches cannot be reduced to one without distorting both activity and selectivity. To keep the model tractable, we retained only the three dominant barriers, thioester formation, hydrolysis and aminolysis, with rate constants *k*_thioester_, *k*_acid_ and *k*_amide_ (Fig. 2e). The overall *k*_cat_ then follows from these simulated barriers (Methods).

The microkinetic treatment markedly improved agreement with the experimental *k*_cat_ and captured the mutational effects (Fig. 2f). Its advantage is greatest for competing pathways: only when one branch becomes negligible does the model collapse to the conventional single-RDS Eyring limit,^42^ as for M1-T234G, whose hydrolysis is effectively switched off (*k*_acid_ ≪ *k*_amide_; Fig. 2f). Quantitatively, accounting for the full network brought the calculated *k*_cat_ to within |Δ log| = 0.67 of experiment compared to |Δ log| = 2.28 when only *k*_amide_ was considered (Fig. 2f). Extending the approach to a saturation-mutagenesis panel at position 259, the predicted *k*_cat_ reproduced the relative-activity trend across variants and, more importantly, largely recovered the product selectivity: the amide/acid ratio correlated positively with experiment (Pearson *r* = 0.41), and the qualitative selectivity was correctly predicted for 14 of 20 variants (Fig. 2g). Enabled by high-throughput computation that explicitly models each elementary step, this approach provides a dynamic, non-empirical route to quantitative mechanistic analysis of multistep enzyme catalysis and a practical foundation for large-scale rational design.

### 2.5 Substrate scope and atomistic interpretability

To probe the generalizability of the framework and the origins of substrate effects, we computed log_10_ *k*_cat_ for 25 aldehyde–amine combinations spanning aliphatic and aromatic amines across several variants (Fig. 3a). Across all 25 products, the calculated log_10_ *k*_cat_ correlated with the experimental yield (*r* = 0.48, Fig. 3b). Thus, despite the limited dynamic range of the experimental yields (51–85%) and additional experimental and computational uncertainties, the calculated log_10_ *k*_cat_ captured the dominant experimental trend.

**Figure 3.**
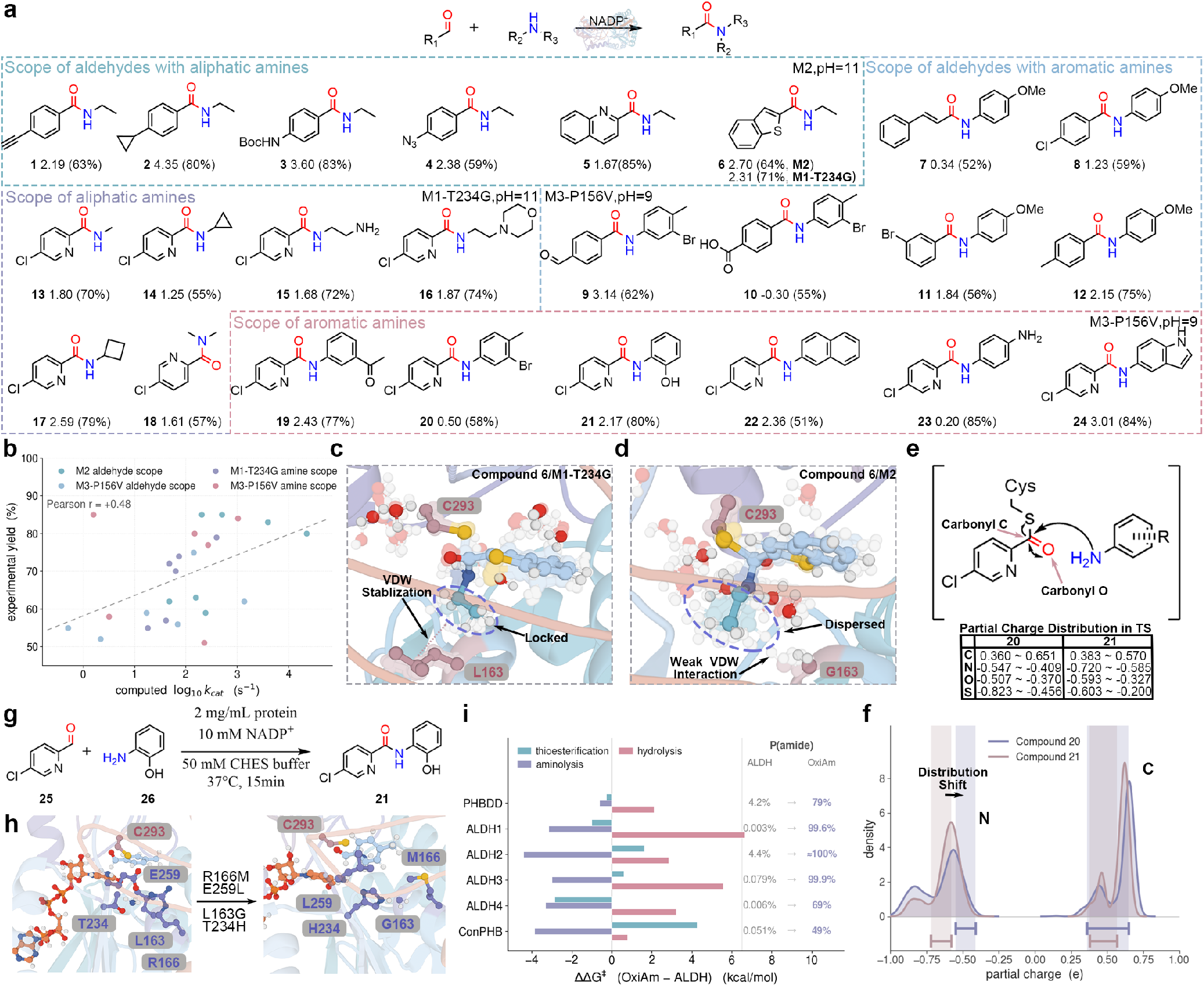
Substrate scope and transfer of OxiAm activity across the ALDH family. (**a**) Substrate scope over diverse aldehyde and amine pairs; for each product, the calculated log10 *k*_cat_ is followed by the experimental yield in parentheses. (**b**) Calculated log10 *k*_cat_ versus experimental yield across the 25 substrate–enzyme combinations (Pearson *r* = 0.48). (**c**,**d**) Representative aminolysis transition-state ensembles of compound **6** in two variants: the L163 environment locks the substrate through van der Waals stabilization (**c**), whereas G163 permits a more dispersed ensemble with weaker interactions (**d**). (**e**) Aminolysis reaction scheme, with the ranges of the reactive-atom partial charges in the transition state for compounds **20** and **21.** (**f**) Partial-charge distributions of the carbonyl carbon and the nucleophilic nitrogen sampled over the entire simulation; shaded bands mark the corresponding transition-state charge ranges. (**g**) Model amidation reaction and conditions used to compare engineered members of the ALDH family. (**h**) Active-site comparison highlighting the four substitutions associated with the OxiAm phenotype. (**i**) Change in activation free energy, 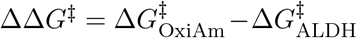, for thioester formation, aminolysis and hydrolysis across the ALDH family, and the resulting shift in the predicted probability of amide formation, *P* (amide).

The physical grounding of the model allowed these trends to be traced to their atomistic origins. Compound **6** provides a clean example, showing different yields between two variants that differ at position 163 (Fig. 3c,d). Analysis of the aminolysis transition-state ensemble showed that the N-ethyl group of compound **6** is conformationally locked in M1-T234G, where the L163 side chain stabilizes it through van der Waals contacts, whereas in M2 the N-ethyl group sampled a broader range of conformations because of the absence of this contact with G163 (Fig. 3c,d). This stabilization explains the higher yield in M1-T234G (71%) than in M2 (64%).

Because the machine learning potential predicts physically meaningful atomic charges (Extended Data Fig. 6a), enables direct dissection of substituent-induced electronic effects. For compounds **20** and **21**, bearing an electron-withdrawing bromide and an electron-donating hydroxyl, respectively, we extracted the partial charges of the four reactive atoms across the transition-state ensemble (Extended Data Fig. 7a–c). The attacking nitrogen becomes markedly more negative in the presence of the electron-donating substituent, resulting in a clear shift in its charge distribution (Fig. 3e,f), whereas the carbonyl carbon, remote from the substitution site, showed nearly identical charge distributions for the two compounds (Fig. 3e,f). Electronic effects alone, however, do not fully account for the observed yields. Analysis of the transition-state ensembles further revealed an important role for neighbouring-group participation: the chloropyridine nitrogen can engage in the hydrogenbond network and and stabilize the oxyanion of the tetrahedral intermediate (Extended Data Fig. 7f,g). Together, these atomistic analyses of mutational, electronic and neighbouring-group effects demonstrate how the framework resolves distinct molecular determinants of catalytic activity and reveals their underlying chemical mechanisms.

### 2.6 A shared strategy converts diverse ALDHs through divergent mechanisms

Previous work identified four mutations (L163G, R166M, T234H and E259L) that convert a broad range of ALDHs into OxiAms^27^ (Fig. 3g,h). Our calculations show that this shared outcome arises through distinct atomistic mechanisms (Fig. 3i). Across all six scaffolds, the mutations lower the aminolysis barrier, favouring amide over acid product formation at the branch point downstream of the thioester. In ALDH2, ALDH3 and ConPHB, they additionally raise the barrier of the thioesterification, the common step preceding both branches, such that amide becomes the major product but overall turnover is reduced. In PHBDD, ALDH1 and ALDH4, by contrast, the thioesterification barrier is lowered and the competing hydrolysis barrier raised, improving selectivity and efficiency simultaneously; among them, ALDH4 is the most effective engineered variant, consistent with previous experiments^27^ and highlighting it as a promising target for further engineering. Thus, a single macroscopic result can arise from fundamentally different atomistic mechanisms across closely related scaffolds. Because thioesterification and competing hydrolysis remain incompletely controlled, deliberately tuning these barriers through rational design represents an opportunity for further improvement. Resolving such hidden mechanistic differences highlights the value of our field-aware, ML-based framework for physically grounded, atomistic enzyme engineering.

### 2.7 Applyting OxiAm to pharmaceutical amide synthesis

Biocatalysis is increasingly being adopted in industrial pharmaceutical manufacturing, and OxiAm has already shown promise for amide bond construction in drug synthesis.^27^ ABBV-318, a Na_V_1.7/Na_V_1.8 blocker for the treatment of pain, is assembled through a palladium-catalysed carbonylation followed by stoichiometric HATU coupling of an enantiopure fluoropyrrolidine^43^ (Fig. 4a). We devised a chemoenzymatic route in which a single OxiAmcatalysed oxidative amidation of aldehyde **38** replaces this three-step activation–coupling sequence, eliminating both the precious metal catalyst and the coupling reagent (Fig. 4b).

**Figure 4.**
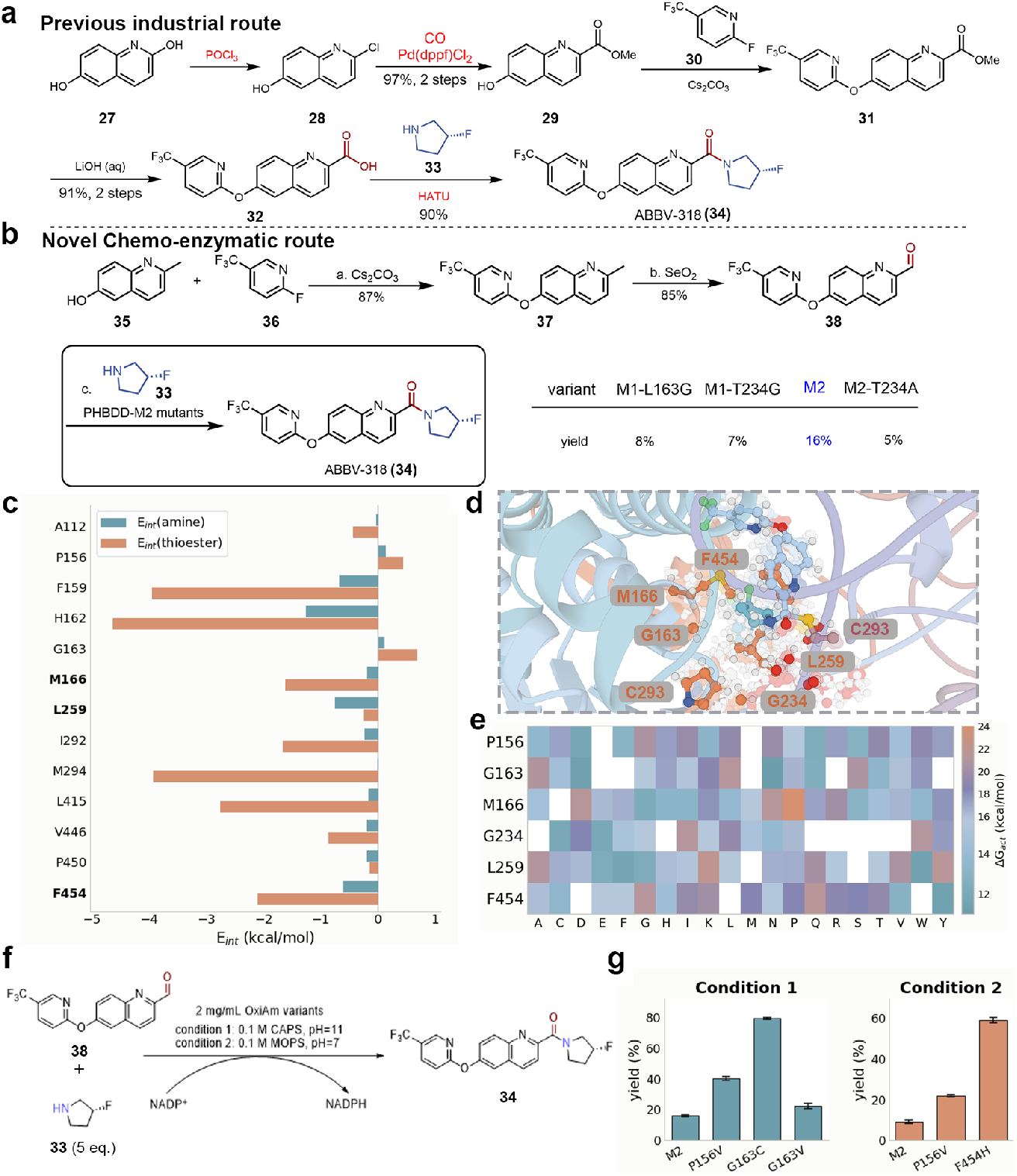
Mechanism-guided engineering of OxiAm for the chemoenzymatic synthesis of ABBV-318. (**a**) Previous industrial route to ABBV-318 (**34**); steps requiring stoichiometric or precious-metal reagents are highlighted in red. (**b**) Chemoenzymatic route developed here. Right, yields of **34** for the first round of engineered variants. Reaction conditions: (a) 1.3 eq. **36**, 1.5 eq. Cs_2_CO_3_; (b) 2.0 eq. SeO_2_; (c) PHBDD variants were screened under the following condition: 2 mg mL*−*1 enzyme, 5 mM **38**, 25 mM **33**, 6 mM NADP+, 0.1 M CAPS, pH = 11, 10% v/v DMSO, 37 °C, 16 h. The reaction was quenched with 400 *µ*L MeCN and analyzed by UPLC-MS. The yields were calculated from UPLC peak areas and quantified using UPLC calibration with authentic standards. (**c**) Per-residue decomposition of the interaction energy *E*_*int*_ between non-reacting active-site residues and the two reacting partners, the amine nucleophile (teal) and the thioester intermediate (orange), averaged over the aminolysis transition-state ensemble of M2. The ten largest contributors are shown; the three positions selected for redesign are in bold. (**d**) Representative aminolysis transition-state configuration of the ABBV-318-forming complex in M2, showing Cys293 and the six positions taken forward for redesign. (**e**) Virtual saturation mutagenesis at these six positions: computed activation free energy Δ*G*_*act*_ of the aminolysis step for all 19 substitutions. White cells denote barriers above 24 kcal mol*−*1 or no productive reaction. (**f**) Model reaction and conditions used to test the top-40 ranked variants under condition 1 (0.1 M CAPS, pH 11) and condition 2 (0.1 M MOPS, pH 7). (**g**) Variants improving yield more than 1.5-fold over M2 under condition 1 (left) and condition 2 (right). General conditions: 2 mg mL*−*1 enzyme, 5 mM **38**, 25 mM **33**, 6 mM NADP+, 0.1 M CAPS, pH 11 or 0.1 M MOPS, pH 7, 10% v/v DMSO, 37 °C, 16 h. The reaction was quenched with 400 *µ*L MeCN and analyzed by UPLC-MS. The yields were calculated from UPLC peak areas and quantified using UPLC calibration with authentic standards. Data are shown as mean ± SEM (n = 3, biologi1ca4l replicates).

Extensive experimental mutagenesis nonetheless yielded only one competent variant, M2, and at low yield (Fig. 4b). Taking M2 as the starting point, we decomposed the interaction energy across the aminolysis transition-state ensemble to identify the ten residues that couple most strongly to the reacting partners (Fig. 4c). Among these, F454 has its side chain that points into the reaction centre, whereas M166 and L259 clamp the amine from opposite faces (Fig. 4d). Together with positions 159, 163 and 234, previously shown to control the OxiAm phenotype, these six sites were selected for virtual saturation mutagenesis, with the barrier for each variant obtained from ML/MM OPES-MetaD sampling (Fig. 4e).

The 40 top-ranked variants were assayed under two pH conditions (Fig. 4f and Extended Data Fig. 8). At pH 11, G163C increased the yield to nearly 80%. We then screened the same panel at pH 7, a milder and more process-compatible condition under which the parent is barely active. At pH 7, F454H increased yield to 60%. Notably, position 454 lies outside the sites explored in previous engineering campaigns and was nominated solely by the transition-state interaction analysis (Fig. 4c,d). Using a threshold of more than 1.5-fold improvement in yield, the calculated hit rates are 3*/*40 (7.5%) and 2*/*40 (5%) for the two pH conditions, respectively (Fig. 4g). These gains were made on a scaffold already refined by an exhaustive experimental campaign, highlighting the difficulty of further optimization within the vast remaining sequence space. Despite this challenge, productive variants emerged from computational ranking alone, demonstrating that the ML/MM framework can prioritize beneficial substitutions beyond previously explored engineering positions. High-throughput free-energy calculations of this kind therefore offer a promising route to expanding the searchable sequence space while reducing reliance on empirical screening, providing a general strategy for rationally guiding the design of improved biocatalysts.

## 3 Conclusion

We have presented an accurate, efficient and physically grounded computational framework for enzyme engineering. Across diverse reaction types, mutations, substrates and homologues, the framework maintained chemical accuracy and, when coupled with microkinetic modelling, enabled the high-throughput, non-empirical treatment of complex catalytic systems. Applied to OxiAm, it guided the discovery of variants for pharmaceutical amide synthesis, demonstrating its potential for practical biocatalyst design.

Considerable opportunities for further development remain. Further improvements in computational efficiency would enable high-throughput campaigns at substantially larger scales. The present microkinetic treatment connects simulated free-energy barriers to *k*_*cat*_ but omits substrate binding, product release and cofactor exchange, which may become rate-influencing in other systems and will require explicit treatment. The simulation workflow is also amenable to automation, with AI models guiding the exploration of reaction pathways and analysis of the resulting ensembles.

We nonetheless anticipate that continued advances in efficiency, automation and mechanistic modelling will establish this framework as a powerful platform for free-energy-driven enzyme engineering, enabling rational exploration of sequence space and accelerating the design of improved biocatalysts for chemical and pharmaceutical applications.

## 4 Data and Code Availability

The software developed in this work is available at https://github.com/ClickFF/MLMM4AMBER. The source code would be integrated into future AMBER release. The AutoNCAA package is available at https://github.com/Hsuchein/AutoNACC. The source code of the MACEPOLEF is available at https://github.com/ClickFF/MACEPOL-EF.

## 5 Methods

### 5.1 Dataset preparation

Initial structures were taken from the OMol25 dataset,^44^ restricted to closed-shell systems containing the 14 elements H, B, C, N, O, F, Si, P, S, Cl, As, Se, Br and I. Sampling roughly 500 structures per atom count from 2 to over 200 atoms yielded ~100,000 structures. All calculations used the *ω*B97M-V functional with the def2-TZVPD basis set, ^45–47^ the def2/J auxiliary basis with the RIJCOSX approximation,^48–50^ the DEFGRID3 integration grid, and self-consistent treatment of the VV10 non-local correlation for gradient evaluations, consistent with the OMol25 reference protocol, ^44^ conducted by ORCA 6.1.0.^51^ For every structure we computed the total energy, the analytical nuclear gradient (forces), and minimal-basis iterative stockholder (MBIS) populations,^52^ with an otherwise identical field-free calculation providing the zero-field baseline. External fields were of two types, inhomogeneous and homogeneous.

For anisotropic inhomogeneous fields applied to a molecule of *N* atoms at positions 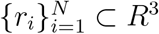 with elements {*e*_*i*_}, we required the external field at each atomic site to follow a prescribed, element-dependent Gaussian along each Cartesian axis *a* ∈ {*x, y, z*},

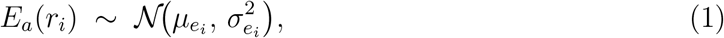

where the per-element parameters (*µ*_*e*_, *σ*_*e*_) set the range of electrostatic environments. Since the perturbation has no preferred sign, we took *µ*_*e*_ ≈ 0 and calibrated the second moment *σ*_*e*_, which controls the field strength.

External charges were placed on two atom-centred spherical shells of radii *R* ∈ {2.0 Å, 3.5 Å}. On each shell, *P* points (default *P* = 4) were distributed quasi-uniformly by the Fibonacci-sphere construction,^53^ *z*_*k*_ = 1 − (2*k* + 1)*/P, ϕ*_*k*_ = 2*πk/φ* with 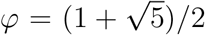. Candidate charges closer than *d*_min_ = 1.5 Å to any atom were discarded, and the total was capped at *M* = min(5*N*, 300) by uniform subsampling. Within the point-charge approximation the field at atom *i* is linear in the charge vector *q* = (*q*_1_, …, *q*_*M*_)^⊤^; collecting the 3*N* Cartesian components into *E* ∈ *R*^3*N*^,

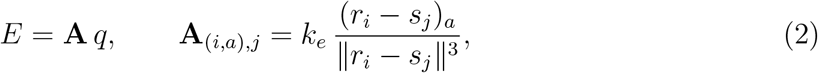

with *k*_*e*_ = 14.3996 VÅ*/*e the Coulomb constant. Because **A** is severely ill-conditioned (*κ* ~ 10^6^), a direct inverse *q* = **A**^+^*E*^*⋆*^ yields unphysical charges (~ 10^5^ *e*) that collapse once clipped to [−1, +1] *e*. We therefore sampled forward: charges were drawn from a zero-mean Gaussian, 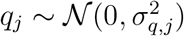, whose variance was calibrated so that the field ensemble satisfied Eq. (1). As each field component is a linear combination of independent charges, its variance propagates exactly,

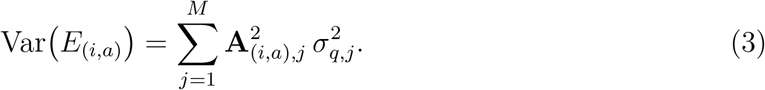

To avoid the underdetermined per-charge fit, each charge *j* was assigned to the element group *g*(*j*) of its nearest atom and all charges in a group shared one variance 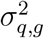, reducing Eq. (3) to Var 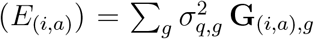 with 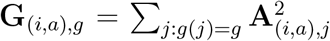. Writing 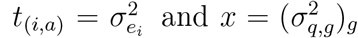, the group variances follow from a non-negativity-constrained least-squares problem,

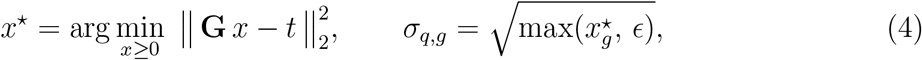

solved by NNLS^54^(*ϵ* = 10^*−*12^). The constraint *x* ≥ 0 enforces physical (non-negative) variances and renders the problem well posed. For each requested configuration the charges were then realised and clipped to the physical range,

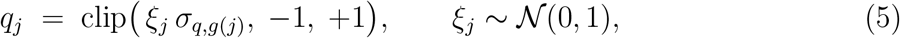

drawn from a single seeded stream for reproducibility. The realised ensemble *E* = **A***q* reproduced the target per-element widths to within a factor of 0.75–1.12 across H, C, N and O.

The second type applied a spatially uniform field directly. For each structure a single magnitude was drawn uniformly, *r* ~ U(0, *r*_max_), with *r*_max_ (default 2.5 V Å^*−*1^) a user-controlled bound, so that the zero-field limit was sampled continuously within the same distribution. To probe orientational response, *N*_f_ unit directions {*u*_*k*_} were drawn uniformly on *S*^2^ (*u* = *g/*∥*g*∥, *g* ~ N(0, **I**_3_)) and accepted by rejection under a minimum-pairwise-angle constraint,

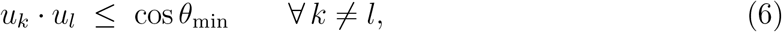

so that all pairwise angles satisfy *θ*_*kl*_ ≥ *θ*_min_ (default 30^°^). The field for direction *k* is *E*_*k*_ = *r u*_*k*_.

For each structure, a separate calculation was performed under every generated field configuration together with the field-free baseline, and the resulting energies, forces and charges constituted the BioPol-EF dataset (~280,000 entries).

### 5.2 Model architecture

MACEPOL-EF predicts the response of a molecule to a static external electrostatic environment. The total energy of a configuration {_*i*_, *Z*_*i*_} under an external potential *V* and field = −∇*V* is written as the pretrained baseline plus two explicit field-coupling terms,

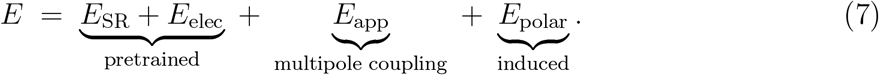

The baseline is the pretrained MACEPOLAR-1 potential,^34^ a higher-order *E*(3)-equivariant message-passing network; ^55^ its short-range and self-consistent electrostatic terms, and its per-atom readout of the on-site monopoles *q*_*i*_, are retained in their original functional form (all parameters are subsequently fine-tuned; see Computational details). Of the two coupling terms, *E*_polar_ is new, whereas *E*_app_ reformulates the per-system field coupling of the baseline as a per-atom, potential-based one. Forces follow from automatic differentiation of Eq. (7). The backbone used here carries only *l*=0 node features, so the on-site charge distribution reduces to monopoles and the coupling to the external field is evaluated at monopole order,

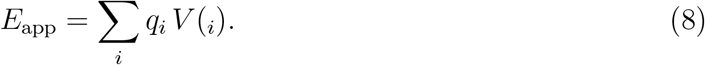

For a homogeneous field the scalar potential is defined relative to a reference origin **R**_0_,

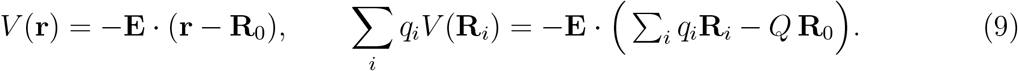

The monopole term therefore already contains the coupling of the field to the charge-position dipole Σ_*i*_ *q*_*ii*_, so no separate·*µ*_tot_ term is needed. For a net-charged system (*Q* ≠ 0) the energy depends on the reference origin through *Q* **R**_0_; in practice the fields are generated by finite external point charges, for which the natural gauge *V* → 0 at infinity fixes the reference unambiguously. Dipolar and higher-order responses are carried by the induced branch.

The induced branch expands the on-site polarisation energy in the external field at atom *i*,

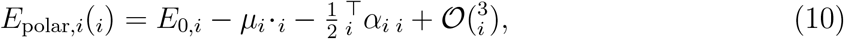

where the response coefficients depend only on the last-layer equivariant message **m**_*i*_ and are realised as rotationally invariant contractions of **m**_*i*_ against tensor powers of the field, making the field dependence at each order an exact linear or quadratic form. Decomposing the quadratic response into an isotropic (*l*=0) trace and a traceless anisotropic (*l*=2) part yields four field invariants: the scalar 0_*i*_, the first-order coupling ⟨1_*i*_,_*i*_ ⟩, and the two second-order responses 0_*i*_|_*i*_|^2^ and ⟨2_*i*_, *Y*_2_(_*i*_)⟩, with *Y*_2_ the unnormalised degree-2 real spherical harmonics so that the last invariant is exactly quadratic in. Each is a Clebsch–Gordan reduction to *l*=0 and hence an exact *O*(3) invariant, and the induced energy is a per-atom multilayer perceptron of these invariants,

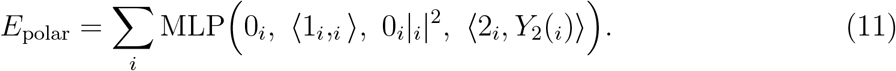

The head is not a truncated expansion but a nonlinear function of the invariants appearing at each order of Eq. (10): the explicit low-order invariants fix the correct field dependence, while the network is free to resum higher-order response. Because the head consumes only **m**_*i*_, whose irrep signature is identical across backbone sizes, a single head design transfers between model sizes unchanged. Its output projection is zero-initialised, so the induced branch vanishes identically at initialisation and training starts from the pretrained baseline. After training, the retained zeroth-order channel 0_*i*_ lets the branch also supply a small field-independent residual correction, while the field dependence is carried entirely by the remaining three invariants.

### 5.3 Electrostatic embedding for ML/MM

The system is partitioned into a ML region, described by a MLIP, and a MM region, described by a classical force field, following the general multiscale partitioning of QM/MM methods.^18^ The total potential energy is

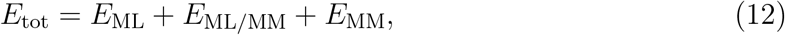

where *E*_MM_ collects all bonded and nonbonded interactions internal to the MM region and is evaluated entirely with the classical force field; *E*_ML_ is the energy of the ML region; and *E*_ML*/*MM_ is the coupling between the two regions,

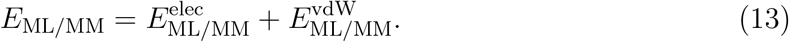

The van der Waals coupling 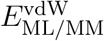 is retained at the classical Lennard–Jones level of the force field for every ML–MM pair. The electrostatic interactions 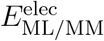 is evaluated in the electrostatic environment created by the MM charges using MLIPs. At every step the MM region is reduced, at each ML site *a*, to a scalar electrostatic potential *ϕ*_*a*_ and a vector electric field **E**_*a*_; these are supplied to the model together with the ML coordinates and elements.

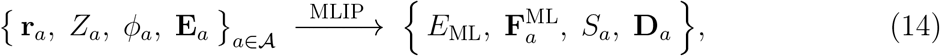

where A indexes the ML sites (real ML atoms and link atoms), 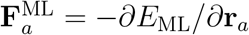 are the intrinsic ML forces at fixed embedding, and

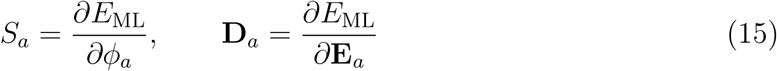

are the sensitivities of the model energy to the embedding potential and field. Physically *S*_*a*_ is the effective (field-polarized) partial charge carried by site *a* and **D**_*a*_ is its induced dipolar response; both are returned by a automatic-differentiation pass, alongside 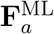. The energy *E*_ML_ produced by this pass already contains the ML–MM electrostatic coupling 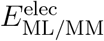: the model is trained to retain its applied-field work term, so the interaction of the ML charge density with the embedding (*ϕ*, **E**) is evaluated internally rather than as a separate classical sum. We keep 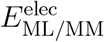 as a label in Eq. (13) to make the coupling explicit, but it is not double-counted: it enters *E*_tot_ once, through *E*_ML_.

Let *x* ∈ X the MM charge sources. Writing **r**_*ax*_ = **r**_*a*_ − **R**_*x*_ and *r*_*ax*_ = ∥**r**_*ax*_∥, the potential at ML site *a* is a screened Coulomb sum over the MM charges *Q*_*x*_,

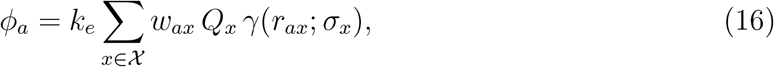

with *k*_*e*_ the Coulomb constant and the screened kernel *γ* defined in Eq. (19). The pair weight

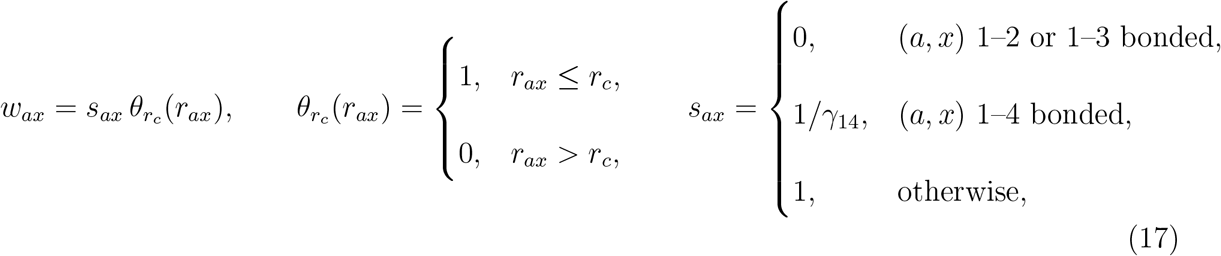

imposes the topological bookkeeping and a real-space cutoff. Following the standard AMBER convention, 1–2 and 1–3 bonded pairs are excluded, 1–4 pairs are scaled by the inverse of the force-field factor *γ*_14_, and all remaining pairs interact fully. The cutoff function 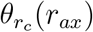 equals one for *r*_*ax*_ ≤ *r*_*c*_ and zero otherwise.

The electric field at ML site *a* is obtained consistently as the negative gradient of the same potential,

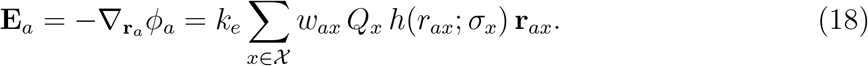

To avoid the short-range singularity of point charges, we represent each MM charge by a normalized spherical Gaussian of element-specific width *σ*_*x*_.^56^ The resulting potential kernel and radial field coefficient are

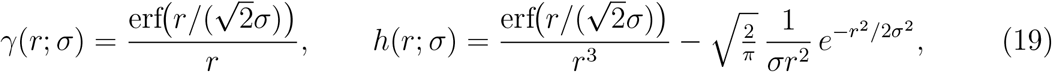

which remain finite as *r* → 0 and satisfy 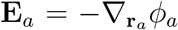. In the point-charge limit *σ*_*x*_ → 0, they recover the bare Coulomb forms *γ* → 1*/r* and *h* → 1*/r*^3^.

Every force is −*∂E*_tot_*/∂***R**. The Lennard–Jones and MM-internal terms are classical; the electrostatic contribution comes entirely from *E*_ML_, which depends on the atomic positions both directly, through the ML coordinates **r**_*a*_, and indirectly, through the embedding *ϕ*_*a*_(**R**) and **E**_*a*_(**R**). Combining the autograd sensitivities of Eq. (15) with the analytic derivatives of the screened kernel yields a clean split between the two regions.

The forces on the ML sites at fixed embedding, 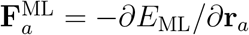, are returned directly by the model’s backward pass and already include the polarization of the ML density by (*ϕ*, **E**). The additional dependence of an ML site’s own *ϕ*_*a*_, **E**_*a*_ on its coordinate is captured by the analytic terms below (which act on both partners of each pair).

An MM atom *x* enters *E*_ML_ only through the potentials and fields it generates at the ML sites. Applying the chain rule through Eqs. (16) and (18) and contracting with the model sensitivities gives two analytic response channels. The potential-response (from *S*_*a*_ = *∂E*_ML_/ *∂ ϕ*_a_), using ∇_ra_ *ϕ*_ax_ = −**E**_ax_

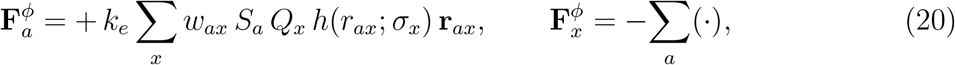

whose leading term (*S*_*a*_ ≈ *q*_*a*_) is exactly the screened Coulomb force between the ML charge and *Q*_*x*_; the correction accounts for the response of the charge to the potential. The field-response (from **D**_*a*_ = *∂E*_ML_*/∂***E**_*a*_) is

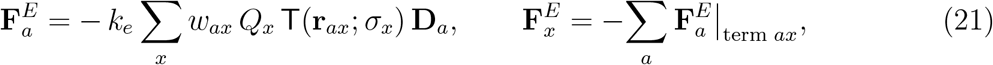

with the screened field-gradient tensor

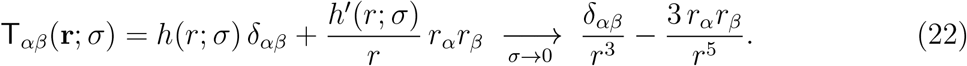

Equations (20) and (21) are the exact chain-rule image of the model’s sensitivity to its inputs: because *S*_*a*_ and **D**_*a*_ are taken with the explicit *ϕ*_*a*_, **E**_*a*_ tensors held fixed, and the geometric dependence is supplied analytically, the two channels together reproduce −*∂E*_ML_*/∂***R**_*x*_ with no double counting. Because all channels share the screened kernel of Eq. (19), the propagated forces are the exact gradient of the reported energy and the dynamics conserve energy.

### 5.4 General Molecular Modelling Settings

Classical MD simulations were performed with `PMEMD.CUDA` and the subsequent reactive simulations with our ML/MM module in `SANDER`, both based on the AMBER source code. ^25,26^

The protein was described with the AMBER14SB force field^57^ and water with TIP3P.^58^ The covalent acyl–enzyme (thioester) intermediate formed between the catalytic cysteine and the substrate aldehyde was treated as a single non-natural residue comprising the cysteine backbone and side chain and the covalently bound acyl group: the broken valences to the neighbouring residues were capped with acetyl and methylamine groups, the capped structure was optimized in Gaussian^59^ at the *ω*B97X-D/6-31G* level,^60,61^ and RESP charges^62^ were derived at HF/6-31G* and constrained to an integer total charge. The capping groups were removed with AutoNCAA^63^ to give the residue template, the remaining parameters were assigned with GAFF2,^64^ and an additional parameter file supplies the cross terms at the AMBER14SB/GAFF2 junction between the cysteine backbone and the acyl group. The same protocol was applied to the deprotonated tyrosine and arginine residues required at high pH. Ligands (amine nucleophiles, free aldehydes) and the NAD^+^/NADP^+^ cofactors were parameterized with GAFF2, with RESP charges obtained in *Antechamber* following *ω*B97X-D/6-31G* optimization and an HF/6-31G* single point.

### 5.5 Microkinetic modelling

The catalytic cycle passes through a single thioester intermediate, formed by acylation with rate constant *k*_thioester_ and consumed by two parallel deacylation channels, aminolysis (*k*_amide_) and hydrolysis (*k*_acid_), both of which regenerate free enzyme; we write *k*_*d*_ = *k*_amide_ + *k*_acid_. Hydrolysis is thus a fully competent turnover that returns the enzyme to the same state as aminolysis while delivering the alternative product, so the productive channel is defined only relative to its competitor and cannot be captured by any single barrier. We treated substrate and both nucleophiles as saturating, binding, product release and cofactor exchange as fast relative to the three chemical steps, and each chemical step as irreversible over one turnover; rate constants followed transition-state theory with unit transmission coefficient^65^ at *T* = 310.15 K (*RT* = 0.616 kcal mol^*−*1^),

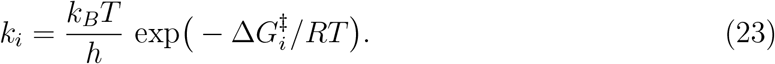

The steady state on the thioester under saturating substrate gives the total, amide and acid turnover numbers, all sharing one denominator,

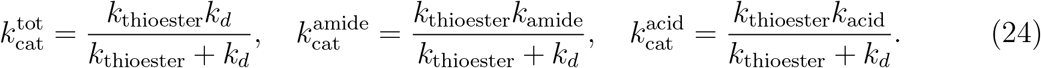

With the branching probability *P*_amide_ = *k*_amide_*/k*_*d*_, the amide turnover factorises exactly as

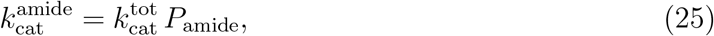

so that how fast the enzyme cycles 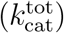 and which product it makes at the fork (*P*_amide_) are mathematically independent factors.

The product ratio is set by deacylation alone: the thioester concentration cancels between the two competing fluxes, so

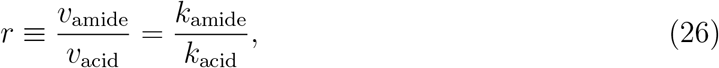

which contains no *k*_thioester_ and is blind to the acylation step. When the nucleophile-bound sub-states of the thioester interconvert rapidly relative to the chemical steps (the Curtin– Hammett condition^66^), their populations drop out and the branch is set entirely by the two deacylation transition states,

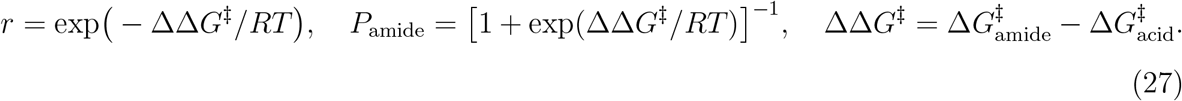

Both deacylation barriers must therefore be referenced to a common thioester basin with a single free-energy zero *G*_thioester_; barriers taken from separate local minima do not define a valid ΔΔ*G*^*‡*^. Because everything entering through *G*_thioester_ cancels, together with the prefactor and much of the systematic error shared by the two channels, ΔΔ*G*^*‡*^ is generally expected to be more robust than either absolute barrier considered independently.

Casting the amide turnover number in Eyring form defines an apparent activation free energy,

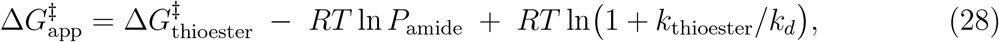

in which the branch enters as a non-negative selectivity cost −*RT* ln *P*_amide_ added to the acylation barrier. When acylation is rate-limiting (*k*_thioester_ ≪ *k*_*d*_) the third term vanishes and 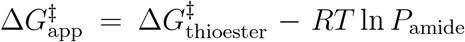 if hydrolysis additionally dominates the branch (ΔΔ*G*^*‡*^ ≫ *RT*) this becomes 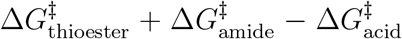: the apparent barrier of the branched mechanism is not the height of any single transition state but a composite of three, and it depends on the competing channel with the sign opposite to the productive one, so that raising the hydrolysis barrier lowers the effective barrier to amide formation. All turnover numbers and the product ratio were computed from the three simulated barriers 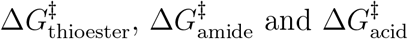 through Eqs. (23)–(28).

### 5.6 Materials

Primer and gene synthesis were conducted at Xianghong Biotech Co. Ltd. DNA sequencing was performed at RuiBiotech Company. ClonExpress II One Step Cloning Kit and TransStart FastPfu DNA polymerase were purchased from Vazyme Biotech and Transgene Biotech respectively. Genes cloned into plasmid pQlinkHx containing an 8× His-tag and a TEV cleavage site in N-terminal. All chemicals and reagents were purchased from commercial suppliers (Sigma-Aldrich, J&K, Alfa Aesar Chemicals, TCI, etc.) and used without further purification unless otherwise stated.

Analytical thin-layer chromatography was performed using 0.25 mm silica gel 60-F plates, using 254/365 nm UV light as the visualizing agent. Flash column chromatography was performed using 200–300 mesh silica gel. Preparative high-performance liquid chromatography (prep-HPLC) was performed using an XBridge Prep C18 OBD column (length, 150 mm; inner diameter, 19 mm; particle size, 5 µm; Waters). Yields refer to chromatographically and spectroscopically pure materials unless otherwise stated.

NMR spectra were recorded on a Bruker 400 MHz NMR spectrometer (AVANCE III) or a Bruker 500 MHz NMR spectrometer (AVANCE III) at ambient temperature with Acetone or DMSO as the solvent unless otherwise stated. Chemical shifts are reported in parts per million (ppm), relative to Acetone (1H, *δ* 2.05; 13C, *δ* 29.84), DMSO (1H, *δ* 2.50; 13C, *d* 39.52) unless otherwise stated. Data for 1H NMR are reported as follows: chemical shift, multiplicity, coupling constant, integration.

High-resolution mass spectrometry (HRMS) was recorded on a Q-TOF (AB SCIEX X500R with ESI source, and Agilent 7250 with EI source), which combines quadrupole precursor ion selection and a high-resolution accurate-mass (HR/AM) Time of Flight (TOF) mass analyzer to deliver mass accuracy.

Ultraperformance liquid chromatography (UPLC)/mass spectrometry (MS) analysis was performed on an ACQUITY UPLC H-class system (Waters) coupled with an SQ Detector 2 equipped with an electrospray ionization source. All the UPLC/MS analysis was conducted on an ACQUITY UPLC BEN C18 column (length, 50 mm; inner diameter, 2.1 mm; particle size, 1.7 µm; Waters) at a flow rate of 0.3 mL/min at 40. The gradient elution consists of 0.1% formic acid–H_2_O (A) and MeCN (B). The standard gradient programs were 5–20% B, 0–2 min; 20–45% B, 2–3 min; 45–100% B, 3–7 min.

### 5.7 Plasmid construction

For protein expression of aldehyde dehydrogenases, linear vectors pQlinkHx were generated by PCR using the primers pQlinkHx-F/R (Supplementary Table 3) and plasmid pQlinkHx-CtGST as template. DNA fragments of PHBDD, ALDH3 and conPHB were amplified with TransStart FastPfu DNA polymerase using corresponding primers listed in Supplementary Table 3 and synthesized genes as templates. The gene fragment of HPA were obtained using the similar PCR conditions as PHBDD utilizing *E. coli* DH5*α* genomic DNA as the template and pQlinkHx-HPA-F/R (Supplementary Table 3) as primers. The resulting DNA fragments were cloned into the linear vector pQlinkHx using ClonExpress II One Step Cloning Kit (Vazyme Biotech) following standard protocols.

The plasmid pQlinkMx-TpNOX, used for NADPH regeneration, was constructed by cloning the TpNOX gene into the pQlinkMx vector containing an MBP-tag followed by a His_8_-tag at the N-terminus. The TpNOX DNA fragment was amplified by PCR using primers pQlinkMx-TpNOX-F/R and a synthesized gene as the template. The linear vector pQlinkMx was generated by PCR amplification with primers pQlinkHx-F/R and plasmid pQlinkMx-Fhb7 as template. All the gene sequences and primers were listed in Supplementary Tables 2 and 3, respectively.

### 5.8 Variant construction

All variants were constructed using the MEGAWHOP PCR technique. DNA fragments containing the target mutations were generated by PCR using the respective forward primers and the standard reverse primer pQlinkHx-Down-R (Supplementary Table 3), with the corresponding plasmids as the template. These fragments served as megaprimers in a subsequent MEGAWHOP PCR amplification, using the corresponding plasmid as templates. The resulting PCR products were digested with *Dpn*I at 37°C for 5 h, followed by enzyme inactivation at 80°C for 20 min. The digested products were then introduced into *E. coli* DH5*α* cells via standard transformation. The sequences of all variants were confirmed by Sanger sequencing (RuiboBiotech). All the primers were listed in Supplementary Table 3.

### 5.9 Activity screening of OxiAm variants

To screen PHBDD variants, reactions were performed in 100 µL of 0.1 M CAPS buffer (pH 11.0, condition 1) containing 2 mg/mL PHBDD variant or *E. coli* harboring PHBDD variant, 5 mM aldehyde **38**, 25 mM (5 equiv.) **33**, and 10% v/v DMSO. The same panel was screened under condition 2, in which the CAPS buffer (pH 11.0) was replaced by 0.1 M MOPS buffer (pH 7.0) and all other components were unchanged. Reactions were incubated at 37°C for 16 h and subsequently quenched with 400 µL of acetonitrile. Precipitated protein was removed by centrifugation at 20,000 × *g* for 15 min, and the supernatant was analyzed by UPLC–MS. The conversion was determined by the corresponding UPLC peak areas and corrected using calibration curves generated with authentic standards.

### 5.10 Purification of OxiAms, its homologs, and TpNOX for activity test

PHBDD and its variants were expressed in *E. coli* DH5*α*. A single colony was inoculated into 15 mL of LB medium containing 100 µg/mL ampicillin and grown overnight at 37°C with shaking at 220 rpm. The entire culture was used to inoculate 1 L of fresh LB medium supplemented with 100 µg/mL ampicillin and 0.5 mM IPTG for protein induction. Cells were grown for approximately 18 h at 37°C with shaking at 220 rpm, harvested, and resuspended in lysis buffer (30 mM Tris-HCl, 0.2 M NaCl, 40 mM imidazole, pH 8.0). Cell lysis was performed by ultrasonication, and the lysate was centrifuged at 15,000 × *g* for 20 min. The supernatant was loaded onto a Ni-NTA gravity column pre-equilibrated with lysis buffer. After washing with lysis buffer, the His_8_-tagged target protein was eluted using elution buffer (30 mM Tris-HCl, 0.2 M NaCl, 0.3 M imidazole, pH 8.0). The eluted protein was supplemented with 10% (v/v) glycerol and concentrated to 15–20 mg/mL using Amicon Ultra centrifugal filters (Millipore, molecular weight cutoff of 30 kDa). Finally, samples were flash-frozen using liquid nitrogen and stored at −80°C for further use.

TpNOX, XylB, and LkADH were expressed in *E. coli* DH5*α* harboring the respective plasmids. The strains were grown at 37°C with shaking at 220 rpm until the OD_600_ value reached 0.8–1.0, then induced with 0.5 mM IPTG and further incubated at 18 °C for 18 h. Proteins were subsequently purified following the same procedure as described for PHBDD variants.

To remove imidazole and exchange the buffer, the eluted protein was dialyzed extensively for 8 h against dialysate buffer (30 mM Tris-HCl, 0.2 M NaCl, pH 8.0), yielding a final imidazole concentration below 1 mM. After dialysis, glycerol was added to a final concentration of 10% (v/v) and the sample was concentrated as described above.

## Acknowledgement

J.W. discloses support for the research of this work from the National Institutes of Health (R01GM147673 and R01GM149705), the National Science Foundation (1955260), and the Pittsburgh Supercomputer Center (BIO210185). X.W. discloses support for the research of this work from the Dean’s Fellowship, School of Medicine, University of Pittsburgh.

O.I. discloses support for the research of this work from the National Science Foundation (NSF) Designing Materials to Revolutionize and Engineer our Future (DMREF) program, grant number DMR-2323749. This work used Expanse at the San Diego Supercomputer Center and Bridges-2 at the Pittsburgh Supercomputing Center through allocation CHE200122 from the Advanced Cyberinfrastructure Coordination Ecosystem: Services Support (ACCESS) program, supported by NSF, grant numbers 2138259, 2138286, 2138307, 2137603 and 2138296.

## 6 Author Contributions

Xujian Wang (X.W.) conceived the study, designed and performed the simulations, analyzed the data, and wrote the original draft. Xiang Qiu (X.Q.) and Kangdelong Hu (K.H.) performed the experimental validation. Yuyang Wu (Y.W.), Runtian Gao (R.G.), and Ilkwon Cho (I.C.) prepared the dataset. Shuhao Zhang (S.Z.) optimized the model performance. Taoyu Niu (T.N.) and Haocheng Tang (H.T.) performed simulations. Xiaoguang Lei (X.L.) reviewed and revised the manuscript. Olexandr Isayev (O.I.) and Junmei Wang (J.W.) acquired funding and reviewed and revised the manuscript.

---

**Extended Data Fig. 1.**
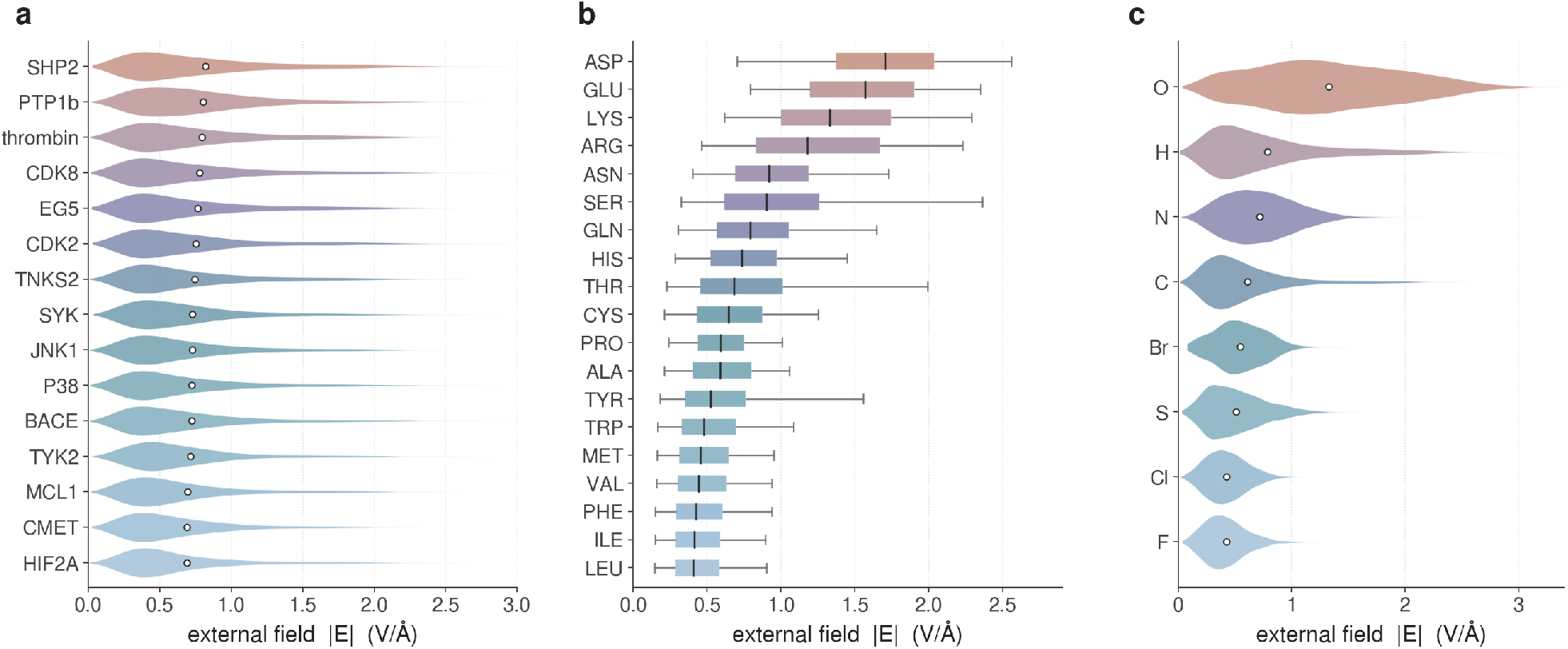
Electrostatic environments sampled across protein–ligand systems. Distributions of the external electric-field magnitude |**E**| acting on individual atoms, extracted from molecular-dynamics simulations of 15 well-characterized drug-discovery targets encompassing more than 600 protein– ligand complexes. The field at each atomic site is computed from the surrounding point charges of the classical force field. (**a**) Violin plots of |**E**| for each protein system, ordered by mean field strength; white circles denote the mean. (**b**) Box plots of |**E**| resolved by residue type, ordered by median. Charged residues (Asp, Glu, Lys, Arg) experience systematically stronger fields than polar and nonpolar residues; boxes span the interquartile range, the central line marks the median and whiskers extend to 1.5× the interquartile range. (**c**) Violin plots of |**E**| resolved by element; white circles denote the mean. Electronegative atoms (O, N) and hydrogen-bond donors (H) sample the broadest and strongest field distributions. These element-resolved statistics define the target distributions used to construct the BioPol-EF dataset (Methods). Electric fields are in V Å^*−*1^.

**Extended Data Fig. 2.**
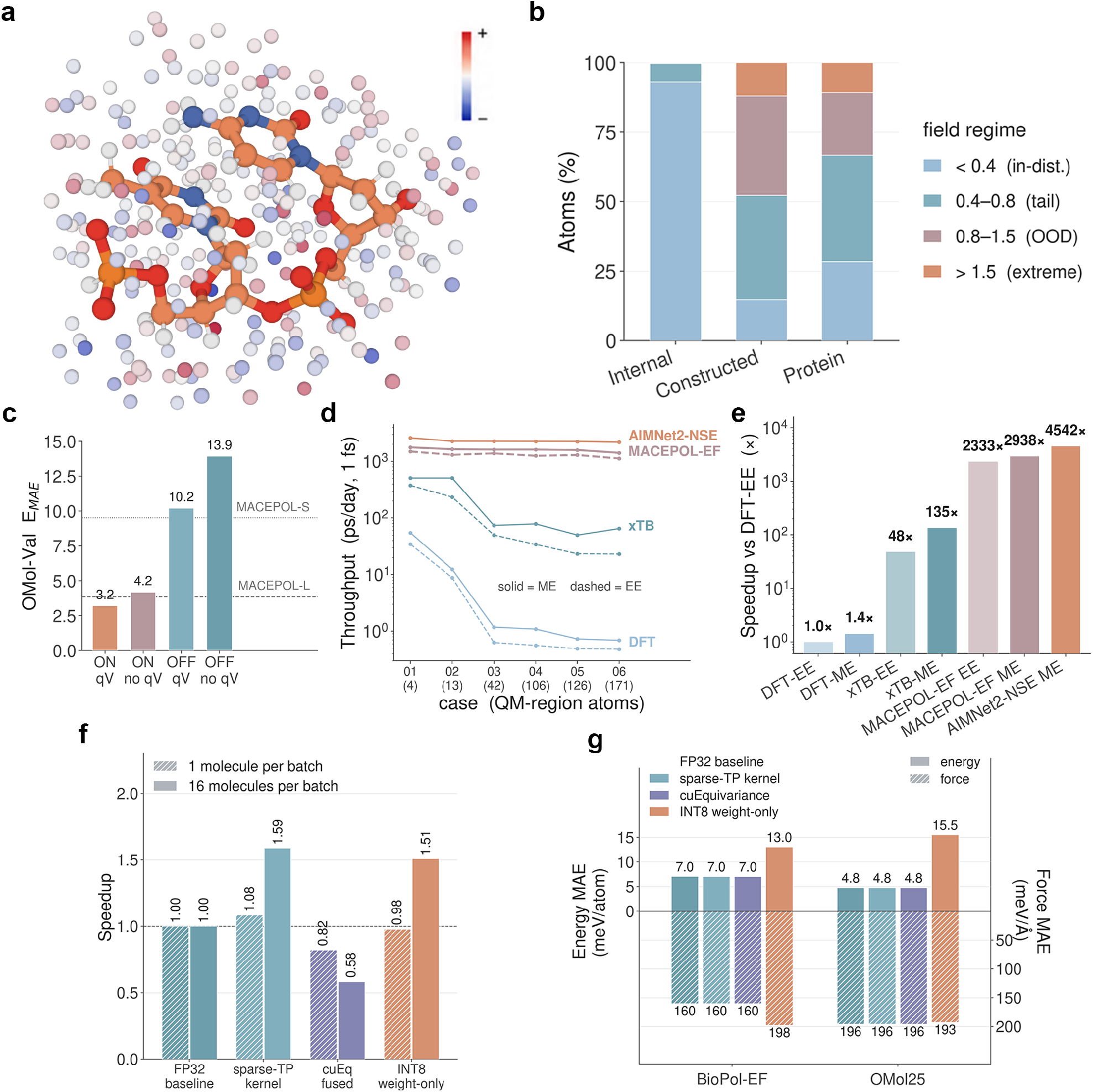
Field-informed dataset construction and the accuracy–efficiency profile of MACEPOL-EF. (**a**) Representative BioPol-EF training configuration: a molecular structure (orange) embedded in external point charges distributed. (**b**) Proportion of atoms in four field regimes for the intramolecular field natively sampled by the base model (Internal), the field generated by the constructed dataset (Constructed) and the true protein-derived external field (Protein). (**c**) Ablation of the adaptive polar head (ON/OFF) and of the charge–potential (*qV*) coupling on the held-out OMol validation set; dashed lines mark the corresponding errors of the MACEPOL-S and MACEPOL-L base models. (**d**) Throughput (ps day^*−*1^at a 1 fs timestep) for DFT, xTB, MACEPOL-EF and AIMNet2-NSE across six systems of increasing QM/ML-region size (atom counts in parentheses), under mechanical (ME, solid) and electrostatic (EE, dashed) embedding. (**e**) Speedup relative to DFT under electrostatic embedding (DFT-EE), measured on the largest test system (case 06). (**f**) Model performance optimization of MACEPOL-EF. The original MACEPOL-EF (FP32) and serves as the reference (speedup = 1); three strategies are then applied to the same trained weights without any retraining, including a sparse tensor-product kernel, fused cuEquivariance kernels, and INT8 weight-only quantization. Bars give the inference speedup over the baseline with 1 molecule (hatched) and 16 molecules (solid) per batch. (**g**) Accuracy cost of each acceleration strategy, reported as energy MAE (meV atom^*−*1^, solid, left axis) and force MAE (meV Å^*−*1^, hatched, right axis) on the BioPol-EF and OMol25 validation sets. 41

**Extended Data Fig. 3.**
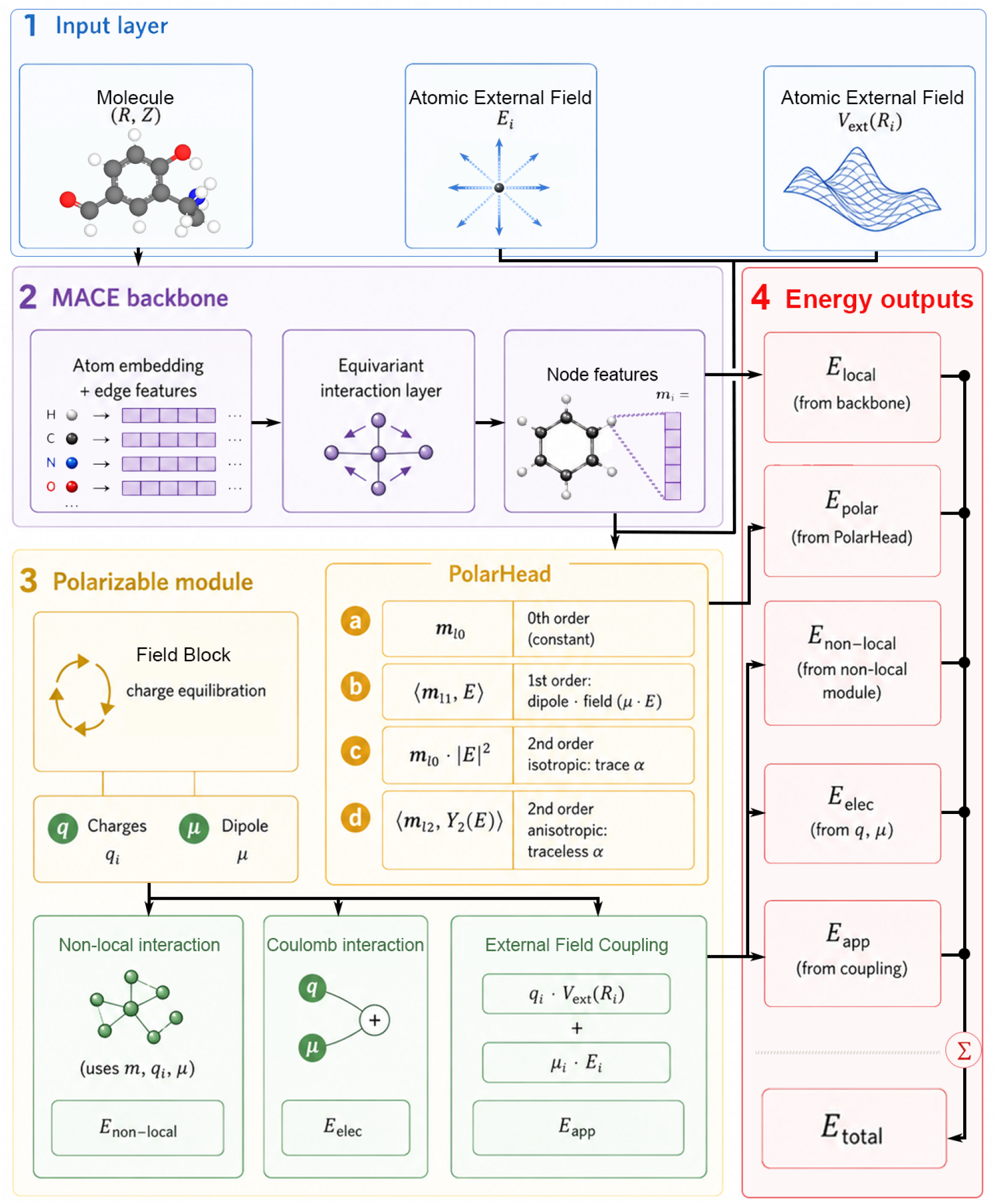
Architecture of the field-aware potential MACEPOL-EF. The model inherits the equivariant backbone of MACEPOLAR-1 and extends it with atom-resolved external-field inputs, a dedicated PolarHead and field-coupled energy terms. (**1**) Input layer: besides the molecular geometry and elements (**R**, *Z*), each atom additionally receives the local external field **E**_*i*_ and external potential *V*_ext_(**R**_*i*_). (**2**) MACE backbone: atom embeddings and edge features pass through equivariant interaction layers to produce node features *m*_*i*_. (**3**) Polarizable module: a Field Block performs charge equilibration to yield atomic charges *q*_*i*_ and dipoles *µ*, while the PolarHead expands the polarization energy in orders of the field—(**a**) zeroth order (constant, *m*_*l*0_), (**b**) first order (dipole–field, ⟨*m*_11_, **E**⟩ = *µ* · **E**), (**c**) second-order isotropic (*m*_*l*0_ |**E**|^2^, trace *α*) and (**d**) second-order anisotropic (⟨*m*_*l*2_, *Y*_2_(**E**)⟩, traceless *α*). The resulting *m, q*_*i*_ and *µ* feed a non-local interaction (*E*_non-local_), a Coulomb interaction (*E*_elec_) and an external-field coupling (*E*_app_ = *q*_*i*_*V*_ext_(**R**_*i*_) + *µ*_*i*_ · **E**_*i*_). (**4**) Energy outputs: the backbone energy *E*_local_, the PolarHead energy *E*_polar_, *E*_non-local_, *E*_elec_ and *E*_app_ sum to the total energy *E*_total_.

**Extended Data Fig. 4.**
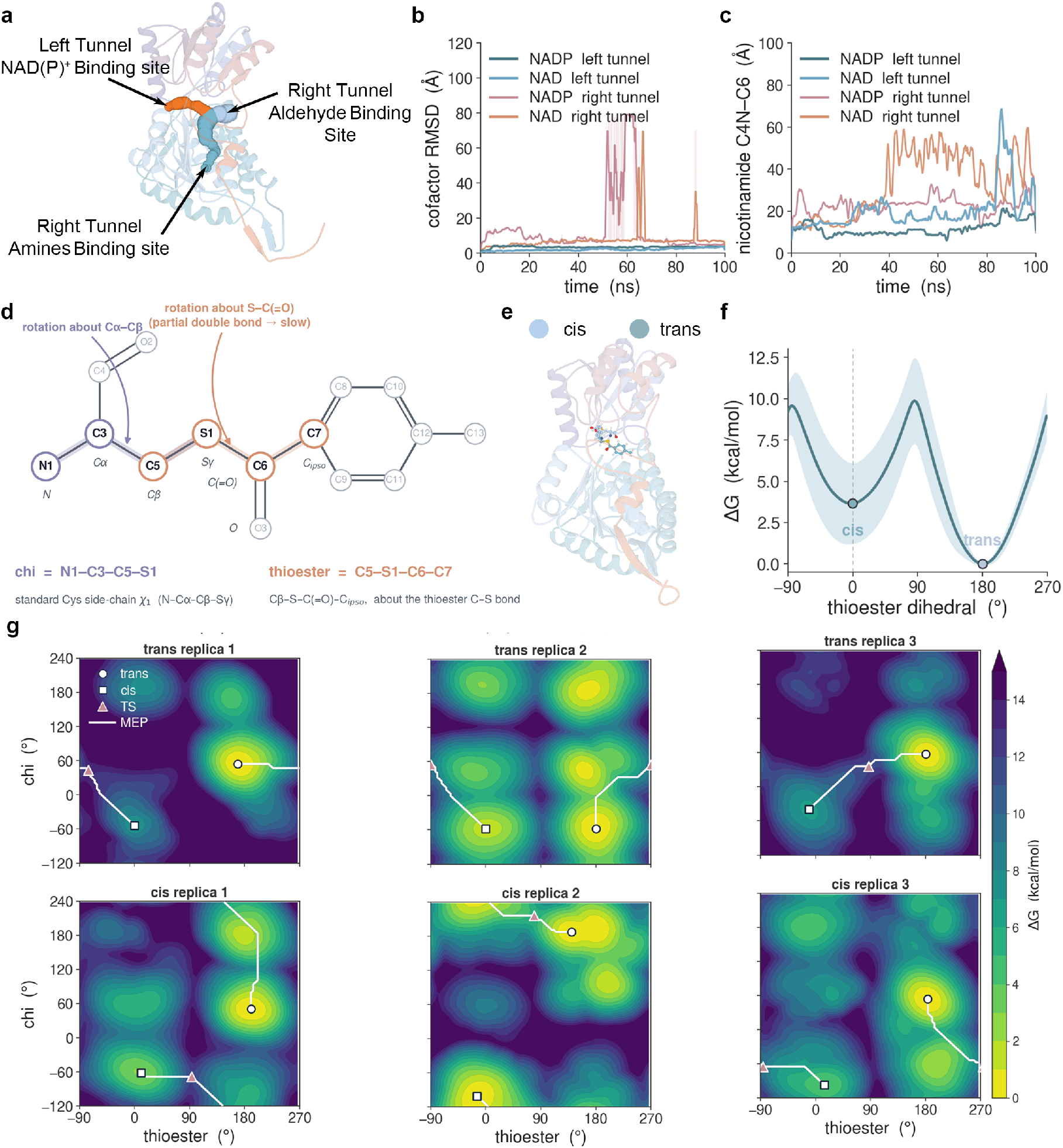
Cofactor binding site and conformational landscape of the thioester intermediate. (**a**) Overall fold showing the left tunnel containing the NAD(P)^+^binding site (orange) and the right tunnel containing the aldehyde and amine binding sites (teal). (**b**) Cofactor RMSD over 100 ns of unbiased molecular dynamics for NAD^+^and NADP^+^placed in either tunnel. (**c**) Distance between the nicotinamide C4N of NAD(P)^+^, the hydride-accepting carbon, and C6, the aldehyde carbonyl carbon of the substrate, over the same trajectories. (**d**) Definition of the two torsions used as collective variables: the cysteine side-chain *χ*_1_ (N1–C3–C5–S1, blue) and the thioester dihedral about the S–C(=O) bond (C5–S1– C6–C7, orange). (**e**) Representative structures of the *cis* (light blue) and *trans* (dark teal) thioester rotamers. (**f**) Free-energy profile along the thioester dihedral; shaded band, standard deviation over three independent replicas. (**g**) Two-dimensional free-energy surfaces in the (*χ*, thioester) plane for three replicas initiated from the *trans* (top) and *cis* (bottom) rotamers. Circles, *trans* minima; squares, *cis* minima; triangles, transition states; white lines, minimum-energy pathway.

**Extended Data Fig. 5.**
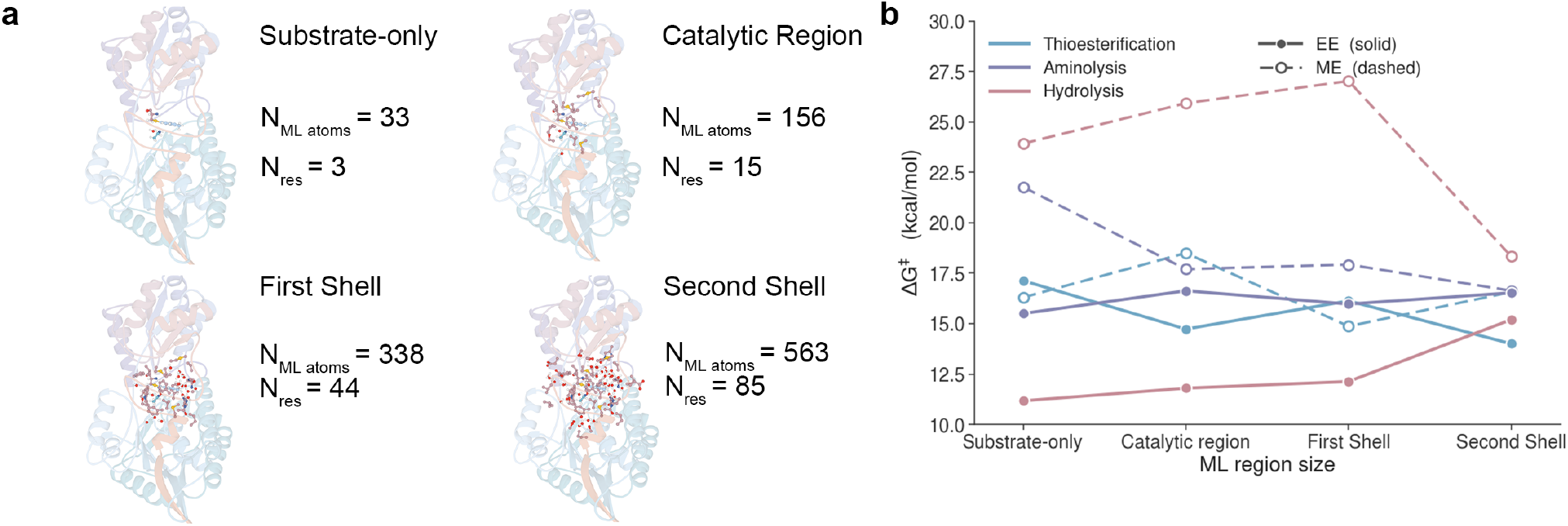
Convergence of activation free energies with ML region size. (**a**) The four ML region definitions used, with the number of machine-learning atoms (*N*_ML atoms_) and residues (*N*_res_) in each: substrate-only (33 atoms, 3 residues), catalytic region (156, 15), first shell (338, 44) and second shell (563, 85). ML atoms are shown as sticks and the remaining MM protein as cartoon. (**b**) Activation free energy Δ*G*^*‡*^ of thioesterification (blue), aminolysis (purple) and hydrolysis (red) as a function of ML region size, under electrostatic (EE, solid, filled circles) and mechanical (ME, dashed, open circles) embedding.

**Extended Data Fig. 6.**
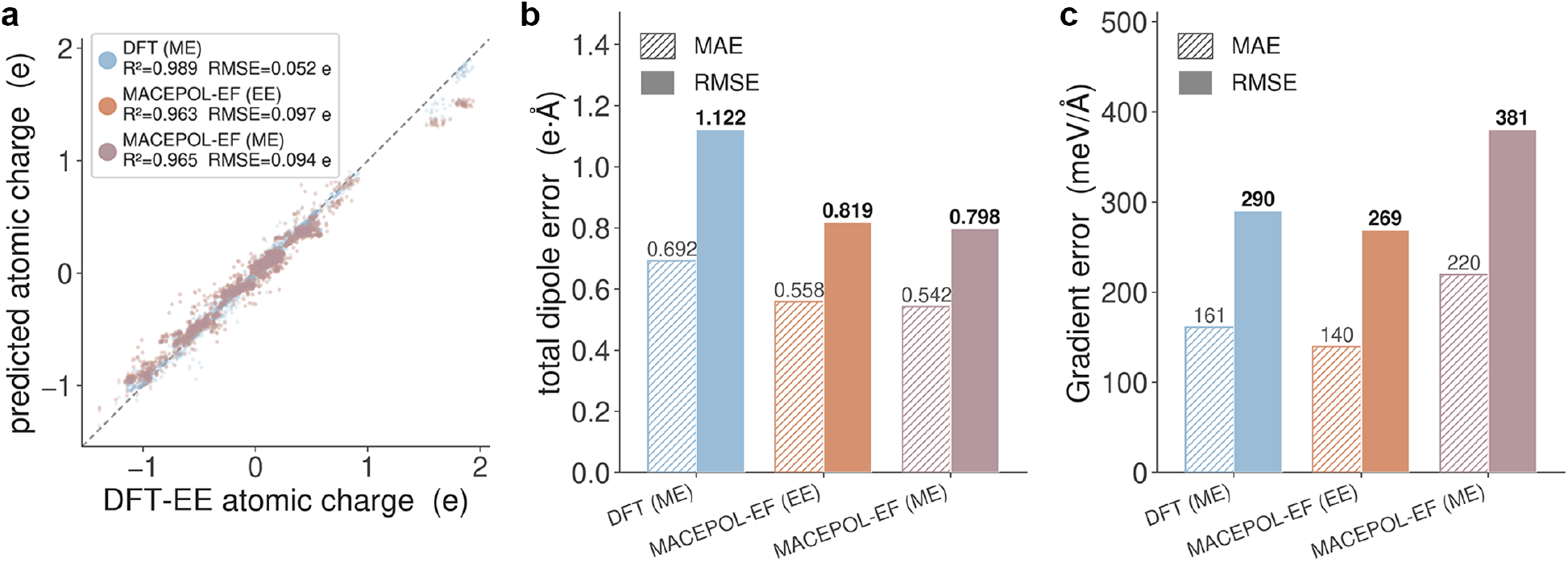
Accuracy of MACEPOL-EF against the electrostatic-embedding DFT reference. (**a**) Predicted versus DFT-EE atomic charges for DFT (ME), MACEPOL-EF (EE) and MACEPOL-EF (ME), evaluated over structures along the reaction pathway; *R*^2^ and RMSE are given for each method and the dashed line marks the identity. (**b**) Total dipole error (e Å) of the same three methods relative to DFT-EE; hatched bars, MAE; solid bars, RMSE. (**c**) Gradient error (meV Å^*−*1^) of the same three methods relative to DFT-EE; hatched bars, MAE; solid bars, RMSE.

**Extended Data Fig. 7.**
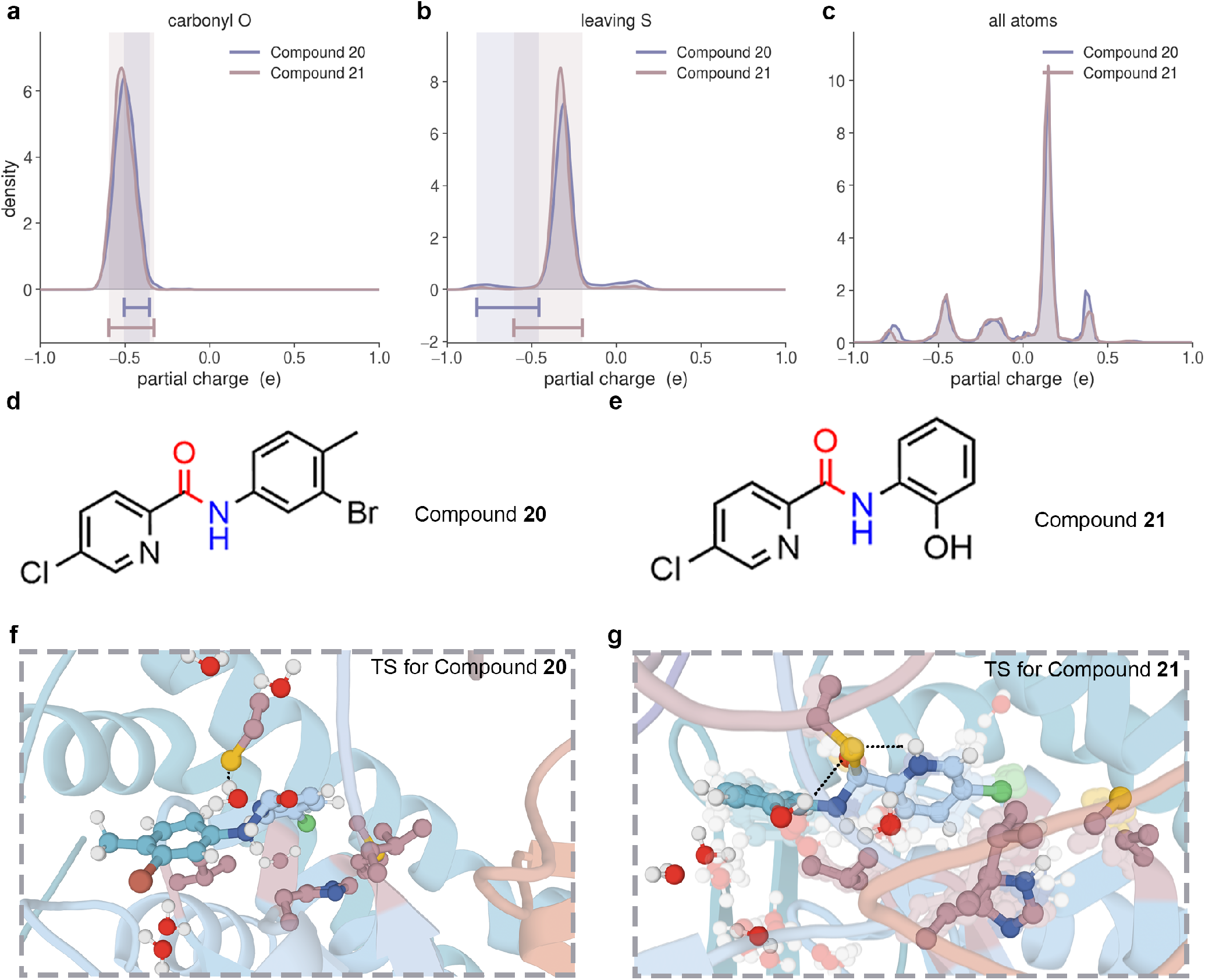
Transition-state partial charges and structures for compounds 20 and 21. (**a–c**) Partial-charge distributions sampled over the aminolysis transition-state ensembles of compound **20** (purple) and compound **21** (red): the carbonyl oxygen (**a**), the leaving sulfur (**b**) and all atoms of the substrate (**c**). Shaded bands and the horizontal bars below each distribution mark the corresponding charge ranges for the two compounds in the transition state. (**d**,**e**) Structures of compound **20**, bearing an electron-withdrawing bromide (**d**), and compound **21**, bearing an electron-donating hydroxyl (**e**). (**f**,**g**) Representative transition-state configurations for compound **20** (**f**) and compound **21** (**g**); dashed lines indicate the hydrogen bonds formed by the chloropyridine nitrogen within the active-site hydrogen-bond network.

**Extended Data Fig. 8.**
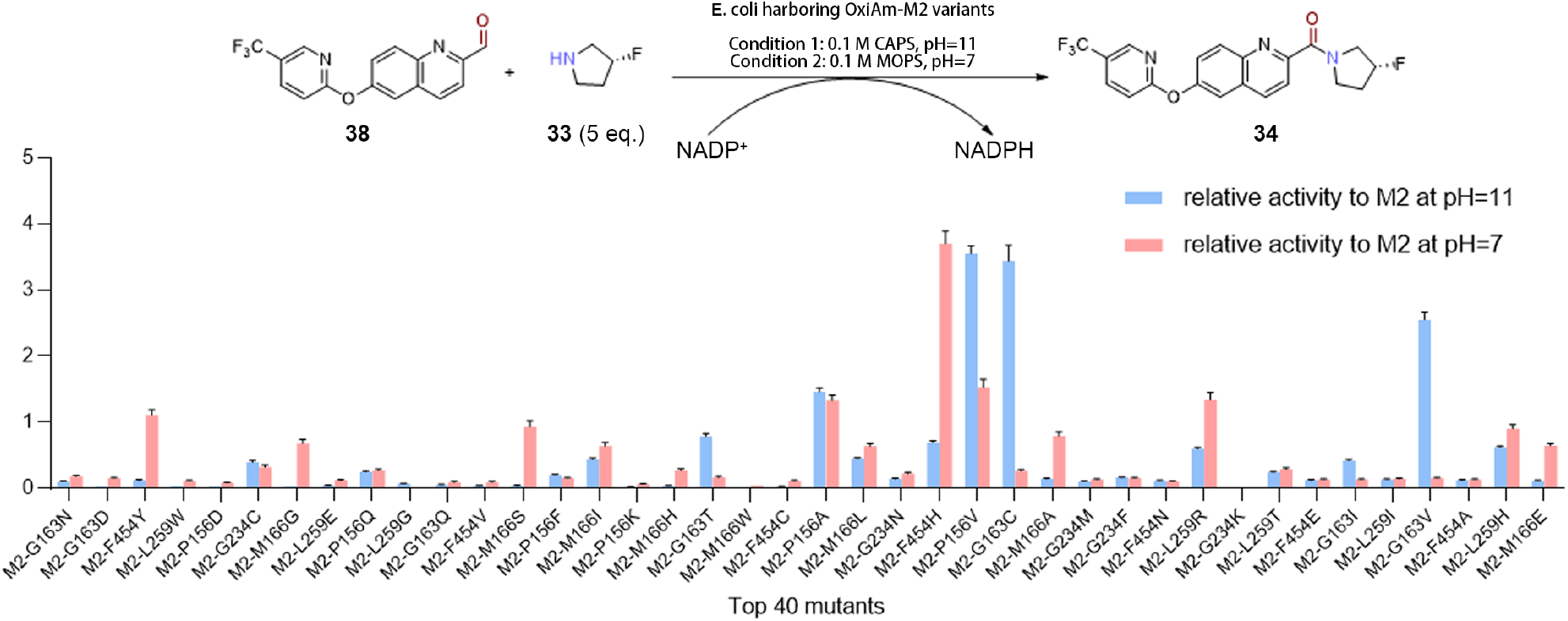
Activity screening of the OxiAm-M2 variant panel. Top, model amidation of the ABBV-318 aldehyde with 5 eq. of 3-fluoropyrrolidine, catalysed by *E. coli* whole cells harbouring OxiAm-M2 variants with NADP^+^/NADPH cofactor recycling, under condition 1 (0.1 M CAPS, pH 11) and condition 2 (0.1 M MOPS, pH 7). Bottom, relative activity of the top 40 variants predicted by calculations normalized to M2 at pH 11 (blue) and at pH 7 (red). General condition: *E. coli* harboring M2 variants resuspended in 200 µL reaction system (5 mM **38**, 25 mM **33**, 6 mM NADP^+^, 0.1 M CAPS, pH 11 or 0.1 M MOPS, pH 7, 10% v/v DMSO), then incubated at 37 ^°^C for 16 h. The reaction was quenched with 500 µL MeCN and analyzed by UPLC–MS. The relative activities were calculated from UPLC peak areas and quantified by comparison with M2. Data are shown as mean ± SEM (*n* = 3, biological replicates).

