## Supplementary material for "High-throughput physics-based enzyme engineering": SI

### Table of Contents

**Theoretical Details** **S4**

Boundary treatment . . . . . S4

**Computational Methods** **S5**

Hardware . . . . . S5

Molecular Dynamics Simulations of Protein–Ligand Complexes . . . . . S5

Calculation of electric fields from the MD trajectories . . . . . S7

Training Dataset . . . . . S9

Training Protocol . . . . . S9

Model Benchmarks . . . . . S10

Ablation Experiment . . . . . S11

Model Performance Optimization . . . . . S11

Protonation states . . . . . S14

Construction of the acyl–enzyme systems . . . . . S15

Construction of the ternary enzyme–aldehyde–NAD(P)$^+$ systems . . . . . S15

Molecular Modelling . . . . . S16

Selection of reactive starting structures . . . . . S17

ML/MM metadynamics . . . . . S17

QM/MM and ML/MM Performance Benchmarks . . . . . S20

ML Region and Embedding Convergence Test . . . . . S21

Single-blind ML/MM screen of ABBV variants . . . . . S22

**Experimental Protocols and Results** **S25**

**Chemo-enzymatic synthesis of ABBV-318 (34)** **S28**

**Compound 37:**: diaryl ether . . . . . S28

**Compound 38:**: aldehyde . . . . . S29

**Compound 34:**: ABBV-318 . . . . . S30

**NMR spectra** **S31**

**References** **S41**

### Theoretical Details

#### Boundary treatment

Covalent bonds severed by the ML/MM partition are treated using our previously reported link-atom scheme, including link-atom placement and force redistribution onto the real boundary atoms.[1] Here, we extend this treatment to electrostatic embedding by combining boundary charge redistribution with virtual dipoles, which restore the local electrostatic environment perturbed by removal of the ML-region force-field charges while preserving the total system charge.[2]

For each boundary $\mathrm{MM_1}$–$\mathrm{MM_2}$ bond, a pair of virtual charges $\pm\delta q$ is placed on opposite sides of $\mathrm{MM_2}$,

$$\mathbf{R}_v = (1 - C_v)\,\mathbf{R}_{\mathrm{MM}_1} + C_v\,\mathbf{R}_{\mathrm{MM}_2}, \qquad (q_v, C_v) \in \big\{(+\delta q,\, 0.94),\, (-\delta q,\, 1.06)\big\}, \tag{1}$$

thereby restoring the local bond dipole without changing the net charge. The virtual charges are included as MM sources and therefore contribute through the same screened kernel as the physical MM charges. Each link atom is treated as a full ML site and receives the resulting embedding potential $\phi_L$ and field $\mathbf{E}_L$.

Because the virtual sites are not independent degrees of freedom, their forces are redistributed to the two real boundary atoms by the chain rule,

$$\mathbf{F}_{\mathrm{MM}_1} \mathrel{+}= (1 - C_v)\,\mathbf{F}_v, \qquad \mathbf{F}_{\mathrm{MM}_2} \mathrel{+}= C_v\,\mathbf{F}_v, \tag{2}$$

providing the link atom with a consistent electrostatic environment while closing all forces onto physical atoms.

### Computational Methods

#### Hardware

All calculations reported in this work were carried out on compute nodes equipped with Intel Xeon Platinum 8358 processors and NVIDIA RTX 6000 Ada Generation graphics processing units, running PyTorch 2.11 against CUDA 13.0. Unless stated otherwise, every timing, throughput and latency measurement reported below was obtained on this hardware and software stack.

#### Molecular Dynamics Simulations of Protein–Ligand Complexes

A total of 640 protein-ligand complexes were prepared and subjected to molecular dynamics (MD) simulations. These complexes encompassed 15 protein targets that have previously been used to benchmark free-energy perturbation methods.[3,4] The initial protein structures were obtained from the Protein Data Bank (PDB; `https://www.rcsb.org/`), and the corresponding PDB IDs and number of ligands are summarized in Table S1.

Table S1: Protein targets, corresponding PDB structures, and the number of ligands.

| **Target name** | **BACE** | **CDK2** | **CDK8** | **c-MET** |
|---|---|---|---|---|
| PDB ID | 4DJW | 1H1Q | 5HNB | 4R1Y |
| No. of ligands | 44 | 31 | 33 | 24 |
| **Target name** | **SYK** | **Thrombin** | **TNKS2** | **TYK2** |
| PDB ID | 4PV0 | 2ZFF | 4UI5 | 4GIH |
| No. of ligands | 44 | 26 | 27 | 91 |
| **Target name** | **Eg5** | **HIF-2$\alpha$** | **JNK1** | **MCL1** |
| PDB ID | 3L9H | 5TBM | 2GMX | 4HW3 |
| No. of ligands | 28 | 42 | 59 | 72 |
| **Target name** | **p38** | **PTP1B** | **SHP2** | |
| PDB ID | 3FLY | 2QBS | 5EHR | |
| No. of ligands | 35 | 48 | 26 | |

Prior to simulation, co-crystallized water molecules, ions, and solvent molecules were removed from the protein structures. Histidine protonation and tautomeric states, and alternate residue conformations were manually assigned to favor chemically reasonable local interactions. Redundant protein chains arising from crystal multimers were removed unless the oligomeric state was considered relevant to the biological function of the target. The resulting protein structures were subsequently used for molecular mechanics calculations.

For the ligands adopted from Goel et al.,[5] the protonation states reported in their study were retained. Protonation states of ligands obtained from Schindler et al.[4] were assigned using the LigPrep module in Maestro.[6] Because most ligands associated with a given protein target shared a common chemical scaffold, their binding poses were generated by manually aligning the conserved core structures to that of the corresponding co-crystallized ligand, thereby maintaining consistent binding orientations across each congeneric series.

Proteins were parameterized using the AMBER ff14SB force field, whereas ligands were parameterized using the General AMBER Force Field 2 (GAFF2). Ligand atomic partial charges were derived using the restrained electrostatic potential (RESP) fitting based on electrostatic potentials calculated at the HF/6-31G* level of theory. RESP charge fitting were carried out using the Antechamber[7] module implemented in AMBER 16. The topology and coordinate files of the resulting protein-ligand complexes were subsequently generated using the TLEaP module.

Unless otherwise specified, all MD simulations were performed under constant pressure. Pressure was regulated using the Berendsen barostat, while temperature was controlled using a Langevin thermostat with a collision frequency of 5.0 $ps^{-1}$. Bonds involving hydrogen atoms were constrained using the SHAKE algorithm. Short-range nonbonded interactions were evaluated using a cutoff distance of 10.0 Å, whereas long-range electrostatic interactions were treated using the particle mesh Ewald (PME) method.

The simulation protocol for each protein-ligand complex comprised four stages. First, energy minimization was performed through five consecutive steps with progressively weakened harmonic restraints applied to the protein and ligand. The restraint force constants applied to the protein were 10, 5, 2, 1, and 0 kcal $mol^{-1}$ $Å^{-2}$, respectively, whereas those applied to the ligand were 100, 50, 20, 10, and 0 kcal $mol^{-1}$ $Å^{-2}$. Each minimization step consisted of 1,000 optimization cycles, including 200 steps of steepest-descent minimization followed by 800 steps of conjugate-gradient minimization.

In the second stage, restrained MD simulations were carried out at 298 K while gradually reducing the restraints on both the protein and ligand. Harmonic restraint force constants of 20.0, 10.0, 5.0, and 1.0 kcal mol$^{-1}$ Å$^{-2}$ were sequentially applied. The system was simulated for 0.2 ns under the 20.0 kcal mol$^{-1}$ Å$^{-2}$ restraint, followed by 1.0 ns simulations at each of the remaining three restraint strengths, resulting in a total duration of 3.2 ns. A time step of 1 fs was used throughout this stage.

In the third stage, the system was gradually heated from 50 to 298 K using a 2 fs integration time step. The heating procedure was divided into six sequential temperature windows with 50 K increments, and 2 ns of MD simulation was performed at each temperature, corresponding to a total heating period of 12 ns.

Finally, production MD simulations were conducted for 90 ns at 298 K using a 2 fs integration time step. The resulting trajectories were subsequently used for downstream structural and energetic analyses.

#### Calculation of electric fields from the MD trajectories

Electric fields were evaluated on the unaltered production trajectories described above; no additional equilibration or restraint was applied. For each complex, 100 configurations spaced 10 ns apart were taken from the final portion of the 90 ns production run, so that every configuration was drawn from a well-equilibrated segment of the trajectory. Each configuration was retained in its original periodic box with all explicit water molecules and counterions present, so that the fields reported below include the explicit hydration and ionic atmosphere rather than a continuum representation of it.

Fields were computed at every atom of a designated probe region, which was excised from the system electrostatically: the region was defined purely as a partition of the atom list, and all atoms remained at their trajectory coordinates. Two classes of probe region were used.

1. **Ligand probes.** All atoms of the bound ligand, taken from every one of the 640 protein–ligand complexes, giving the field exerted on the ligand by the complete protein–solvent environment. Because the ligands of a congeneric series differ from one another, each complex was treated separately.

2. **Side-chain probes.** The side chain of each non-glycine amino acid residue, defined as the residue minus its backbone atoms (N, H, $C_\alpha$, $H_\alpha$, C, O, and, where present, the N-terminal H1/H2/H3 and C-terminal OXT). Glycine, which has no side chain beyond $H_\alpha$, was excluded. Side-chain probes were taken from the first complex of each of the 15 protein targets only, since the members of a congeneric series share a common binding mode and scaffold and therefore present an essentially identical electrostatic environment to the protein side chains; this gave 139–587 side-chain probes per target and 4763 in total.

For a probe atom $i$ at position $\mathbf{r}_i$, the electric field generated by its environment was obtained as a direct Coulomb sum over the fixed force-field partial charges $q_j$ of the surrounding atoms,

$$\mathbf{E}_i = \frac{1}{4\pi\varepsilon_0} \sum_{j \in \mathrm{env}} q_j \frac{\mathbf{r}_{ij}}{r_{ij}^3}, \qquad \mathbf{r}_{ij} = \mathbf{r}_i - \mathbf{r}_j, \tag{3}$$

with $(4\pi\varepsilon_0)^{-1} = 14.3997$ V Å $e^{-1}$, so that $\mathbf{E}$ is obtained in V Å$^{-1}$; the sign convention is such that $\mathbf{E}_i$ points away from a positive environment charge. The charges $q_j$ are those used in the MD simulations themselves, i.e. ff14SB for the protein and RESP-fitted GAFF2 charges for the ligand, so that the fields are strictly consistent with the trajectories from which the configurations were drawn. The environment sum runs over all atoms lying within 8 Å of the probe atom, evaluated under the minimum-image convention using the instantaneous box vectors of each configuration, and excludes all atoms belonging to the probe region's own residue or, for the ligand probes, to the ligand itself; excluding the parent residue in its entirety removes both the self-interaction of the probe and the intramolecular contribution of the covalently attached backbone, neither of which forms part of the environment field that the probe experiences.

Equation 3 was evaluated for every probe atom in every configuration, yielding $8.4 \times 10^5$ per-atom field vectors in total. The analysis of the main text is based on the field magnitude $|\mathbf{E}_i|$, summarised by its mean, standard deviation, median, 5th and 95th percentiles, and maximum, and stratified in three ways: by chemical element (H, C, N, O, S and the ligand halogens F, Cl, Br, I); by protein target; and by residue type. For the last of these, residues were grouped as Charged($-$) (Asp, Glu), Charged($+$) (Arg, Lys), Polar (Asn, Cys, Gln, His, Ser, Thr, Trp, Tyr) and Nonpolar (Ala, Ile, Leu, Met, Phe, Pro, Val), with the ligands forming a separate Substrate group; alternative protonation and disulfide states were mapped onto their parent residue type (HIE/HID/HIP $\rightarrow$ His, CYX/CYM $\rightarrow$ Cys, ASH $\rightarrow$ Asp, GLH $\rightarrow$ Glu, LYN $\rightarrow$ Lys) before grouping.

#### Training Dataset

The BioPol-EF dataset developed in this work was used as the fine-tuning dataset. Configurations with a maximum atomic force above 50 eV $\text{Å}^{-1}$ were discarded, and the remainder was split 85/15 into training and validation subsets, grouping configurations by molecular identity so that no molecule appears on both sides of the split. This yields 178,393 training and 31,482 validation configurations carrying a nonzero embedding field, and a further 33,868/5,976 field-free configurations of the same systems that were held out of the production run (zero-field behaviour is instead anchored by the replay data described below). The training set spans 2–198 atoms (median 43) and net charges from $-2$ to $+2$, with 18% of configurations charged; the mean field magnitude at the nuclei is 0.93 V $\text{Å}^{-1}$.

Fine-tuning a foundation potential on a narrow chemical domain degrades it elsewhere.[8] To counteract this, every training batch was mixed with a 1.5% fraction drawn from 42,337 OMol25 configurations[9] (median 86 atoms), retained with their original reference labels, that sample the chemistry the backbone was pretrained on. The mixing is performed inside the sampler, so each optimiser step sees both distributions.

#### Training Protocol

Starting from the released MACEPOLAR-1-S model,[10] training proceeded in three stages. In the first, all pretrained weights were frozen and only the PolarHead was trained for 10 epochs at a constant learning rate of $2\times10^{-3}$ after one warm-up epoch. The remaining two stages released all parameters at a constant learning rate of $2 \times 10^{-4}$: 11 epochs weighted towards forces, then 9 epochs with the energy weight raised to parity with the force weight. The objective was

$$\mathcal{L} = w_E \left\langle \frac{|\Delta E|}{N_{\text{at}}} \right\rangle + w_F \left\langle |\Delta \mathbf{F}_i|^2 \right\rangle + w_q \left\langle |\Delta q_i| \right\rangle, \tag{4}$$

where $\langle\cdot\rangle$ denotes a mean over configurations, atoms or Cartesian components as appropriate, and $\Delta E$, $\Delta \mathbf{F}_i$ and $\Delta q_i$ are the total-energy, atomic-force and monopole errors, the last taken against the MBIS reference;[11] the charge term was masked for replay configurations, which carry no monopole labels. The stage weights $(w_E, w_F, w_q)$ were $(10, 100, 0)$, $(10, 100, 10)$ and $(100, 100, 10)$.

All stages used Adam[12] with AMSGrad,[13] weight decay $5 \times 10^{-7}$, gradient-norm clipping at 10, single precision, a 6 Å interaction cutoff, and a batch of 12 configurations per device under distributed data parallelism on NVIDIA RTX 6000 Ada GPUs. Checkpoints were written each epoch, and the checkpoint carried into the next stage was selected on held-out data by force error for the force-weighted stages and by energy MAE for the final stage. The learning rate and charge weight of the last two stages were selected on held-out data from sweeps over $\{1, 2, 5\} \times 10^{-4}$ and $w_q \in \{10, 20, 30\}$.

### Model Benchmarks

The model was evaluated on three sets.

*(i) Field-strength sweep.* Eight buckets were generated from a common set of small neutral molecules (10–26 atoms) by scaling the embedding point charges by factors of $0$, $0.33$, $0.67$, $1$, $1.5$, $2$, $3$ and $5$ and recomputing the reference data at the same level of theory, the unit factor corresponding to the condensed-phase field. Each bucket contained 6,000 configurations (three field realisations per structure), except the field-free bucket (2,000) and the $3\times$ and $5\times$ buckets (5,921 and 5,556, the remainder failing to converge). The mean field magnitude ranged from 0.23 to 3.47 V Å$^{-1}$, reaching 10.3 V Å$^{-1}$ locally at the strongest scaling; the molecules were held fixed across the series.

*(ii) In-domain validation.* This set comprised the 31,482 held-out field-embedded configurations described above, whose molecules do not occur in training.

*(iii) Retention on the pretraining distribution.* This set was the OMol25 validation split restricted to closed-shell species of the fourteen elements of the training chemistry (2,085,832 configurations, 75.5% of the split); only energies and forces were evaluated, the set carrying no charge or field labels, and a 27,014-configuration subset was monitored each epoch during

training.

Errors were reported as energy MAE per atom (meV atom$^{-1}$), force RMSE (meV Å$^{-1}$) and monopole MAE against MBIS ($e$). All evaluations used single precision with forces from automatic differentiation and supplied the model with the physical field $\mathbf{E} = -\nabla V$, the internal sign convention being applied once at configuration construction. For the field-embedded sets, positions were referred to the per-molecule barycentre before evaluation; no centring was applied to the field-free OMol25 set.

#### Ablation Experiment

MACEPOL-EF differs from the pretrained baseline in two independent respects: the charge–environment coupling is reformulated as the per-atom potential, and the PolarHead is added. We trained the full $2 \times 2$ factorial of these two changes. The coupling factor took either the reformulated form or the baseline form.[10] The PolarHead was either present or absent; cells without it omitted the head-only pretraining stage and began fine-tuning from the pretrained weights, while all other cells followed the three-stage schedule above. The four cells were otherwise identical: same training and replay data, epochs, learning rates and loss weights in each stage, optimiser settings, and checkpoint-selection rule. The cell with both changes is the production model. Each cell was evaluated on the three benchmarks above.

#### Model Performance Optimization

We explored model compression with MLIP-accelerator and parameter-preserving code optimization with OpenEvolve.[14] The development benchmarks reported in this subsection used an NVIDIA RTX A4000 GPU (16 GB), PyTorch 2.12.0 and CUDA 13.0; dynamic int8 quantization was evaluated on the CPU. These measurements are distinct from the batched MACEPOL-EF validation and inference benchmarks in Extended Data Fig. 3f,g of the main text.

**MLIP-accelerator design.** MLIP-accelerator is a human-in-the-loop, large-language-model (LLM)-assisted workflow that accepts a model repository and checkpoint and produces modified model artifacts with comparative evaluations. Four specialized agents are orchestrated through a LangGraph state machine. The research agent inspects the repository, proposes a model-loading script and inventories the loaded module hierarchy, parameter dtypes and eligible modification targets. The planning agent combines this inventory with source-code checks for dtype constraints to propose a strategy and target modules. Plans are checked deterministically for valid module names and strategy settings before user approval. The modification agent applies the approved transformations, verifies achieved sparsity, dtypes or quantized-layer counts, and exports the artifact with its loading code. The evaluation agent constructs a user-approved harness and compares the original and modified models for energy and force deviations, latency and artifact size. Model loading and evaluation execute in subprocesses within a user-selected software environment.

Each stage requires explicit user approval, and unsuccessful transformations are recorded rather than treated as valid optimized models. Numerical weight transformations are executed by deterministic routines, not generated directly by the LLM. A strategy registry separates plan validation, transformation code and strategy-specific evaluation requirements, supporting magnitude pruning, reduced precision, dynamic int8 and weight-only int8 quantization. After evaluation, users may accept a variant, terminate the run or return the results to the planning agent for another round. Persisted state, approved scripts, plans, verification reports and benchmark outputs allow runs to be resumed and modifications to be audited. The workflow is accessible through command-line and web-chat interfaces.

**Compression trials.** For the 8.33-million-parameter MACEPOL-EF checkpoint, we tested 30% global magnitude pruning, bfloat16 readout and polarization-related heads, dynamic int8 quantization, and weight-only int8 quantization without retraining. An eager model reconstructed from the supplied TorchScript checkpoint enabled changes to module structure. Small-molecule single-point tests showed that pruning substantially perturbed predictions; the zeroed weights remained in dense storage. Dynamic int8 did not provide valid energy-gradient forces and was unsuitable for force-driven simulations. Weight-only quantization instead rounded weights once to a symmetric int8 grid, with per-channel scales for multidimensional tensors and groups of 128 for flattened weights. Activations remained floating point, preserving differentiation with respect to coordinates. Packed int8 storage with on-the-fly dequantization reduced the checkpoint from 34.5 to 9.8 MB. Force mean absolute deviations from the unmodified model were 0.016–0.026 eV $\text{Å}^{-1}$ on water, methanol and ammonia; these are deviations from the original predictions, not errors against quantum-chemical labels. The initial compression trials did not establish a reliable GPU speedup or validate accuracy for enzyme simulations.

**Parameter-preserving code evolution.** OpenEvolve (v0.3.2) performed 50 iterations of code evolution targeting `SparseUvuTensorProduct` in the MACEPOL-EF codebase, benchmarked with the pretrained `polar-1-m` model rather than the fine-tuned MACEPOL-EF checkpoint. This operator is called six times within the self-consistent charge-equilibration calculation: four `dot_products` calls and two `tp_out` calls. Its optimization is distinct from weight pruning. Candidate edits retained the module interface and parameter/buffer layout and were evaluated in separate subprocesses. Checkpoint compatibility was first checked by loading the reference state dictionary. Outputs and gradients with respect to both inputs and weights were then compared with the reference in float64, at an absolute tolerance of $10^{-9}$, for both operator configurations, both feature layouts and input sizes of 7 and 64. Only passing candidates were eligible for a nonzero performance score.

The integrated implementation reassociated tensor contractions to avoid the large pairwise intermediate of shape $B \times 512 \times 512$ and replaced contractions with explicit matrix multiplications and layout-aware views. It uses eager PyTorch and adds no instance state. A faster candidate achieved a $58.5\times$ block-level score but was not integrated because it depended on initialization-time attributes absent from the existing serialized models. Its CUDA float32 compiled path was also not directly covered by the float64 parity gate. Accordingly, that candidate is not used for the model-level results reported below.

**Definition of the block-level speedup.** The integrated candidate's $33.7\times$ result is a call-count-weighted arithmetic mean of speed ratios for an isolated operator, not a speedup of the complete model. For each configuration, float32 forward and backward execution was timed at $B = 216$ and $1024$, using 10 warm-ups followed by 50 synchronized measurements and cycling through four pregenerated input sets. Here $B$ counts input feature rows (atomic sites), not molecules per batch. With $t^{\mathrm{ref}}_{c,B}$ and $t^{\mathrm{opt}}_{c,B}$ denoting reference and optimized forward-plus-backward times per call, the selection score was

$$S_{\text{block}} = \frac{1}{2(4+2)} \sum_{B\in\{216,1024\}} \left[ 4\frac{t^{\text{ref}}_{\text{dot},B}}{t^{\text{opt}}_{\text{dot},B}} + 2\frac{t^{\text{ref}}_{\text{out},B}}{t^{\text{opt}}_{\text{out},B}} \right] \simeq 33.7. \tag{5}$$

The weights 4 and 2 reflect operator call counts, with equal weighting of the two input sizes. This score is neither a ratio of summed execution times nor an end-to-end inference speedup. In particular, the operator benchmark includes gradients with respect to inputs and weights, whereas the model benchmark measures energy and force evaluation.

**Model-level performance and scope.** Separately from OpenEvolve, corrections to the cuEquivariance conversion path resolved feature-layout and PolarHead configuration/compatibility problems without changing trained parameters. These corrections preceded code evolution and are not an OpenEvolve-generated model variant. On a 216-atom diamond supercell, complete energy-and-force evaluation took 367.4 ms with the original implementation, 140.5 ms with corrected cuEquivariance support, and 84.0 ms with both cuEquivariance and the integrated OpenEvolve implementation. Timings are means of 20 synchronized evaluations after five warm-ups. Thus, OpenEvolve provided a $1.67\times$ additional model-level speedup over the corrected cuEquivariance baseline; the $4.37\times$ improvement over the original code includes both contributions. Without cuEquivariance, the OpenEvolve change reduced the same evaluation from 367.4 to 308.6 ms ($1.19\times$). These results are specific to the tested model, system and execution setup; neither the block score nor the model-level ratios represent full ML/MM trajectory throughput or establish the speedup for the different workloads in Extended Data Fig. 3f,g.

#### Protonation states

Missing side chains and hydrogens of the crystal structure (PDB 9JW2) were rebuilt with PDBFixer;[15] crystallographic waters were discarded and the solvent regenerated. Protonation states were assigned from PROPKA3 $pK_a$ predictions[16] at the pH of the biotransformation: a titratable group was modelled deprotonated when its predicted $pK_a$ lay below the simulated pH, and histidines as the neutral $\varepsilon$-tautomer. The catalytic cysteine was always modelled as the thiolate, overriding the PROPKA3 assignment. All hydrogens were stripped before assembly and rebuilt by *tleap* according to the assigned residue names. Mutants were generated by truncating the target side chain to C$\beta$ and renaming the residue, with *tleap* rebuilding the side chain from the AMBER14SB template; multiple mutations were applied sequentially.

#### Construction of the acyl–enzyme systems

For each substrate the cysteine–thioester adduct was assembled, grafted and docked with AutoNACC,[17] an in-house tool built on Open Babel,[18] RDKit[19] and LeDock.[20] The adduct was generated from the aldehyde by replacing the aldehydic C–H bond with the C–S bond to the catalytic cysteine, pre-capped and parameterized as a non-natural residue, and the quantum-optimized adduct was grafted onto the catalytic cysteine by Kabsch superposition of the seven conserved core atoms (backbone N, C$\alpha$, C, O and side-chain C$\beta$, S$\gamma$ and the thioester carbonyl carbon), matched by atom name; residual strain was relieved during minimization and equilibration. The amine nucleophile was placed with LeDock using a single box enclosing the active site and substrate channel, generating 20 poses per ligand; poses were ranked by the distance between the nucleophilic nitrogen and the thioester carbonyl carbon rather than by docking score, and the most attack-ready pose was retained, with the nucleophilic atom specified explicitly for substrates containing more than one nitrogen. Docked coordinates were transferred into the RESP-charged template without altering atom types or charges. Hydrolysis systems contain no additional ligand; the attacking water was selected from the equilibrated ensemble.

#### Construction of the ternary enzyme–aldehyde–NAD(P)$^+$ systems

For the oxidative half-reaction the catalytic cysteine is the free thiolate and the substrate the free aldehyde. The NADP$^+$ pose was transferred from a homologous crystal structure with resolved cofactor (the ternary complex of AlDHPyr1147 with NADP$^+$ and isobutyraldehyde, PDB ID 5EK6): our sequence was aligned to the template chain by global pairwise alignment, the matched C$\alpha$ atoms were superimposed by Kabsch fitting, and the crystal cofactor coordinates were transformed into our frame. The coordinates were then mapped onto the RESP-charged cofactor template by atom name, hydrogens were rebuilt from local frames of their parent heavy atoms, and the charge values were copied verbatim. The free aldehyde was positioned by pinning its carbonyl carbon at the apex of the near-attack triad defined by the cysteine sulfur and the nicotinamide C4 (both distances 3.2 Å) and rotating the molecular body about that apex to minimize steric overlap.

#### Molecular Modelling

**System setup.** Systems were built with *tleap* (AMBERTOOLS) and solvated in a TIP3P box with at least 10 Å padding between solute and box edges. $Na^+$ and $Cl^-$ were added to neutralize the system, and for a subset the salt concentration was adjusted to 0.15 M. The systems contained approximately 60,000–75,000 atoms, and total charges were verified atom-by-atom after assembly.

**Energy minimization.** Each system was minimized by 2,500 cycles of steepest descent followed by 2,500 cycles of conjugate gradient (5,000 total), with all non-solvent heavy atoms restrained to their initial positions at 10 kcal $mol^{-1}$ $Å^{-2}$. Grafted and docked systems were then relaxed by a further 10,000 unrestrained cycles (5,000 steepest descent, 5,000 conjugate gradient); where hard overlaps persisted, the LBFGS (`XMIN`) minimizer was applied before returning to the standard protocol. Bonds to hydrogen were constrained with SHAKE,[21] an 8.0 Å nonbonded cutoff was used throughout, and long-range electrostatics were treated by particle-mesh Ewald.

**Equilibration.** Equilibration used classical MD without MLIPs at a 2 fs timestep. Each system was heated from 100 K to 310.15 K over 500,000 steps (1 ns) in the NPT ensemble, with temperature held by a Langevin thermostat (friction 2.0 $ps^{-1}$) and pressure at 1.0 bar by a Berendsen barostat (relaxation time 2.0 ps). Periodic boundary conditions and an 8.0 Å cutoff for van der Waals and short-range electrostatics were applied.

**Production MD.** Production runs were 50,000,000 steps (100 ns) per system in the NPT ensemble at 310.15 K and 1.0 bar with the same thermostat, barostat, constraint and cutoff settings, saving coordinates every 100 ps.

#### Selection of reactive starting structures

For the acyl–enzyme systems, 100 frames were extracted from a representative window (the last 10 ns of production). The proton transferred from the nucleophile (N–H of the amine or O–H of the water) to the leaving thiolate is mediated by a single bridging water, identified in each frame as the water minimizing $\max[d(\mathrm{O_w}-\mathrm{S}),\ d(\mathrm{O_w}-\mathrm{Nu})]$. A frame was accepted only when this water lay within $d(\mathrm{O_w}-\mathrm{Nu}) \leq 5.0$ Å and $d(\mathrm{O_w}-\mathrm{S}) \leq 5.5$ Å; systems for which no frame satisfied this criterion were reported as failures. The best-scoring frame with the smallest total attack and relay distance was taken as the ML/MM starting structure.

#### ML/MM metadynamics

Reactive free-energy profiles were computed with ML/MM metadynamics. Three chemical steps were treated: (i) oxidation of the aldehyde substrate by $NADP^+$, in which the catalytic cysteine attacks the aldehyde carbon and the resulting thiohemiacetal transfers a hydride to the nicotinamide ring; (ii) aminolysis of the acyl–enzyme thioester by an amine nucleophile, giving the amide product; and (iii) hydrolysis of the same thioester by an active-site water. All three used the identical ML/MM, thermostatting and enhanced-sampling machinery described below and differed only in the collective variables and in the sampling parameters listed under each reaction.

**ML/MM molecular dynamics.** The ML region was described by the MACE-POLAR-1(EF) machine-learned interatomic potential (`mlp_model='macepol_ef'`) under electrostatic embedding (`mlp_embedding=2`). The smeared electrostatic field and scalar potential generated by the MM environment were passed as inputs to the network, and forces on the MM atoms were recovered by automatic differentiation of the model energy with respect to these external quantities, so that ML and MM subsystems polarise one another self-consistently within a single energy evaluation. Charge smearing used a per-element Gaussian width table with a scaling factor of 1.0. The ML–MM interaction range followed the 8 Å MM real-space cutoff.

Trajectories were propagated in the NPT ensemble at 310.15 K and 1.01325 bar with a Langevin thermostat (collision frequency 5 $\mathrm{ps}^{-1}$) and a Berendsen barostat (relaxation time

2 ps), using a 0.5 fs time step. Three independent replicates were run per system from the same equilibrated coordinates, differing only in the random seed used to initialise velocities and the Langevin thermostat (`ig` = $-1$, `irest` = 0). Each replicate was run for 200,000 steps (100 ps). Coordinates were saved every 200 steps (0.1 ps) and energies every 1,000 steps (0.5 ps).

Free-energy sampling used the on-the-fly probability enhanced sampling variant of metadynamics (OPES-MetaD) as implemented in PLUMED, biasing two collective variables simultaneously on a single two-dimensional surface. Kernels were deposited every 100 MD steps (50 fs) at 310.15 K with a neighbour list for the kernel sum, and the compressed kernel state was written every 50,000 steps (25 ps). Initial kernel widths (`SIGMA`) were rescaled adaptively during the run subject to a per-CV lower bound (`SIGMA_MIN`); the values and the `BARRIER` parameter are given per reaction below. One-sided harmonic upper walls were applied to the individual interatomic distances to keep the reacting fragments associated, since the substrate and the nucleophile otherwise diffuse out of the active site once the reactive contact is broken; the wall bias was recorded separately and was accounted for when the surfaces were reweighted. Collective variables, the OPES bias, the reweighting factor $c(t)$, the effective sample size, the number of kernels and each wall contribution were printed every 200 steps (0.1 ps).

**Reaction 1: aldehyde oxidation.** Two collective variables were used. CV1 was the forming sulfur–carbon distance $d(\mathrm{S}_\gamma \cdots \mathrm{C})$ between the cysteine thiolate and the aldehyde carbon, which reports on thiohemiacetal formation. CV2 was the antisymmetric hydride-transfer coordinate $d(\mathrm{H} \cdots \mathrm{C4_N}) - d(\mathrm{C{-}H})$, formed from the distance between the transferring hydrogen and C4 of the nicotinamide ring and the distance between that hydrogen and the substrate carbon. The `BARRIER` parameter was set to 18–30 kcal mol$^{-1}$ (most commonly 22 kcal mol$^{-1}$), with initial kernel widths of 0.06–0.08 Å for CV1 and 0.04–0.14 Å for CV2 and lower bounds of 0.01–0.06 Å and 0.01–0.08 Å respectively. Upper walls were placed at 3.0 Å on $d(\mathrm{S}_\gamma \cdots \mathrm{C})$ with a force constant of 20–50 kcal mol$^{-1}$ Å$^{-2}$ and at 2.0–2.3 Å on $d(\mathrm{H} \cdots \mathrm{C4_N})$ with a force constant of 20–25 kcal mol$^{-1}$ Å$^{-2}$. ML regions for this step contained 127–211 atoms (median 157) with formal charges of 0 or $-1$.

**Reaction 2: thioester aminolysis.** CV1 was the antisymmetric substitution coordinate

$d(\mathrm{N}\cdots\mathrm{C}) - d(\mathrm{C}\text{–}\mathrm{S}_\gamma)$, built from the distance between the amine nitrogen and the thioester carbonyl carbon and the distance between that carbon and the cysteine sulfur, so that it changes sign as the tetrahedral intermediate collapses. CV2 was the mean of the two proton-relay distances through the bridging water, $\frac{1}{2}\left[d(\mathrm{O_w}\cdots\mathrm{H_N}) + d(\mathrm{H_w}\cdots\mathrm{S}_\gamma)\right]$, where $\mathrm{H_N}$ is the proton donated by the nucleophile to the bridging water and $\mathrm{H_w}$ the proton donated by that water to the leaving thiolate; this variable describes the concerted shuttle that deprotonates the nucleophile and protonates the leaving group. The `BARRIER` parameter was set to 15–28 kcal mol$^{-1}$, with initial kernel widths of 0.06–0.10 Å for CV1 and 0.04–0.08 Å for CV2 and lower bounds of 0.01–0.05 Å and 0.01–0.08 Å respectively. Upper walls were placed at 2.5–3.0 Å on the nitrogen–carbon distance (20–50 kcal mol$^{-1}$ Å$^{-2}$), at 4.0 Å on the carbon–sulfur distance (50 kcal mol$^{-1}$ Å$^{-2}$) and at 2.0–2.4 Å on each of the two proton-relay distances (5–50 kcal mol$^{-1}$ Å$^{-2}$). ML regions for this step contained 135–269 atoms (median 170) with formal charges of 0 or $+1$.

**Reaction 3: thioester hydrolysis.** The hydrolysis step used the same pair of collective variables as aminolysis, with the amine nitrogen replaced by the oxygen of the attacking water, so that CV1 is $d(\mathrm{O}\cdots\mathrm{C}) - d(\mathrm{C}\text{–}\mathrm{S}_\gamma)$ and CV2 is the mean of $d(\mathrm{O_w}\cdots\mathrm{H_O})$ and $d(\mathrm{H_w}\cdots\mathrm{S}_\gamma)$ through the same bridging water. Sampling parameters were matched to the aminolysis runs so that the two nucleophiles could be compared directly on the same footing: `BARRIER` was set to 15–28 kcal mol$^{-1}$, with initial kernel widths of 0.06–0.10 Å and 0.04–0.08 Å and lower bounds of 0.01–0.06 Å and 0.01–0.08 Å, and kernels were deposited every 100 steps. Walls were identical to those used for aminolysis. ML regions for this step contained 136–185 atoms (median 154) with formal charges of 0 or $+1$.

**Free-energy surfaces and barriers.** For each replicate the first 10% of the trajectory was discarded as transient. The remaining frames were reweighted with $w_i = \exp[+\beta V_{\mathrm{OPES}}(\mathbf{s}_i)]$, using only the OPES bias, and the two-dimensional free-energy surface was obtained from a weighted Gaussian kernel density estimate on a $120 \times 120$ grid with bandwidths taken from the PLUMED `SIGMA` values. The effective sample size was monitored for every replicate. An independent estimate was obtained directly from the compressed OPES kernel state by reconstructing $P(\mathrm{s})$ from the deposited kernels on a $100 \times 100$ grid with the same normalisation

($Z$, $\epsilon$ and kernel cutoff) used internally by PLUMED, and setting $F(\mathbf{s}) = -k_\mathrm{B}T \ln P(\mathbf{s})$; the two estimates were compared as a convergence cross-check.

The minimum free-energy path between the reactant and product basins was located on the resulting surface by first computing a Boltzmann-weighted shortest path between the two minima (Dijkstra) and then relaxing it with a gradient-descent string method using 200 images. The activation free energy $\Delta G^\ddagger$ was taken as the difference between the highest point of the relaxed path and the reactant minimum, and the reaction free energy $\Delta G_\mathrm{rxn}$ as the difference between the product and reactant minima. Barriers were also extracted from the one-dimensional projection onto CV1 by locating the reactant minimum, the product minimum and the highest intervening maximum. Reported values are means over the independent replicates.

#### QM/MM and ML/MM Performance Benchmarks

To assess the computational efficiency of our ML/MM interface, we performed a systematic benchmarking study by comparing ML/MM and QM/MM simulation speeds across a series of test systems with progressively expanding QM/ML regions.

**QM/MM setup.** QM/MM simulations were carried out through the GROMACS–CP2K interface obtained by joint compilation (GROMACS 2024.2 and CP2K 2024.2),[22,23] which automatically generates a CP2K input template from the system topology. Two QM treatments were compared: density functional theory at the PBE level and the semiempirical GFN1-xTB method, each combined with both mechanical (ME) and electrostatic (EE) embedding, giving four schemes in total. The PBE calculations used the auto-generated template directly; for GFN1-xTB the QM method in the template was switched accordingly, and all other settings were retained. Each simulation used a 0.5 fs time step and ran for 100 steps on 16 MPI processes, and throughput (ps day$^{-1}$) was estimated from the total wall-clock time of the run.

**ML/MM setup.** ML/MM simulations employed the interface developed in this work, using either MACEPOL-EF or AIMNet2-NSE[24] as the ML potential. The same 0.5 fs time step was applied. Each simulation was run for 10,000 steps and the average throughput was computed from the final 1,000 steps.

#### ML Region and Embedding Convergence Test

We scanned the computed activation free energies in two dimensions, over ML region size and ML/MM embedding scheme, for all three chemical steps of the PHBDD M1 (R166M/E259L) catalytic cycle: (i) oxidation of the bound *p*-tolualdehyde, in which the catalytic cysteine thiolate adds to the aldehyde carbon and the resulting thiohemiacetal transfers a hydride to $NADP^+$; (ii) aminolysis of the *p*-toluoyl–cysteine thioester by ethylamine, giving the amide; and (iii) the competing hydrolysis of the same thioester by an active-site water.

Four nested ML region definitions were constructed for each reaction step:

- **Substrate only.** The reaction core alone, i.e. the fragments that make or break bonds along the reaction coordinate: for oxidation, the cysteine thiolate sidechain, the *p*-tolualdehyde and the complete $NADP^+$ cofactor; for aminolysis and hydrolysis, the acyl–cysteine sidechain, the incoming nucleophile (ethylamine or the catalytic water) and the bridging water that mediates proton transfer. No pocket residues and no additional waters.
- **Catalytic (main-text region).** The production ML region of all ML/MM OPES simulations in the main text: the reaction core plus the first-sphere catalytic sidechains.
- **First shell (6 Å).** The catalytic region augmented with every protein sidechain and water molecule having at least one atom within 6 Å of the reaction centre.
- **Second shell (8 Å).** The same construction at 8 Å.

The reaction core was held fixed across all four tiers. Shell radii were measured from the reaction centre rather than from the whole ML region; for the oxidation step the reaction centre comprised the cysteine sidechain, the aldehyde and the nicotinamide ring. Protein residues entered as sidechains only, from $C_\beta$ onwards, with Gly and Pro skipped; water molecules entered as whole molecules; the $NADP^+$ cofactor was always included in full, never truncated at the glycosidic bond; and counter-ions were left at the MM level. The ML-region total charge was set to the formal integer charge of the included fragments and recomputed for every tier.

that carbon and the cysteine sulfur, so that it changes sign as the tetrahedral intermediate collapses. CV2 was the mean of the two proton-relay distances through the bridging water, $\frac{1}{2}\left[d(\mathrm{O_w}\cdots\mathrm{H_N}) + d(\mathrm{H_w}\cdots\mathrm{S_\gamma})\right]$, describing the concerted shuttle that deprotonates the nucleophile and protonates the leaving thiolate. Kernels were deposited every 100 MD steps (50 fs) at 310.15 K; initial kernel widths (`SIGMA`) of 0.06–0.08 Å were rescaled adaptively subject to a per-CV lower bound (`SIGMA_MIN`) of 0.02–0.04 Å, and the `BARRIER` parameter was 20–26 kcal $\mathrm{mol^{-1}}$. A fixed set of one-sided harmonic upper walls kept the reacting fragments associated: 3.0 Å on the nitrogen–carbon distance (20 kcal $\mathrm{mol^{-1}}$ $\mathrm{Å^{-2}}$), 4.0 Å on the carbon–sulfur distance (50 kcal $\mathrm{mol^{-1}}$ $\mathrm{Å^{-2}}$) and 2.2 Å on each of the two proton-relay distances (5 kcal $\mathrm{mol^{-1}}$ $\mathrm{Å^{-2}}$); the wall bias was recorded separately and accounted for on reweighting. ML regions contained 135–205 atoms (median 170) with formal charges of $0$, $+1$ or $-1$.

#### Single-blind ML/MM screen of ABBV variants

**System preparation and classical molecular dynamics.** A prospective, single-blind screen was carried out in which ML/MM aminolysis barriers were computed for a panel of single-point variants before the corresponding experimental activities were disclosed. Variants were generated by near-saturation single-point mutagenesis at six active-site positions — P156, G163, M166, G234, L259 and F454 — of the engineered parent enzyme, each modelled as the covalent acyl–enzyme (a thioester between the catalytic cysteine and the ABBV-derived acyl group) in complex.

Each variant was built with the same automated protocol. The protein was described with the ff14SB force field, water with TIP3P, and the acyl–enzyme intermediate and the amine with GAFF2 and RESP partial charges; non-standard protonation states were assigned from PROPKA at the target pH. The amine was placed in the active site, the complex was assembled in `tleap`, solvated in a rectangular box with a 10 Å buffer and neutralised with $\mathrm{Na^+/Cl^-}$ counterions.

The assembled system was energy-minimised in two stages (restrained heavy atoms, then unrestrained), heated gently from 50 to 298 K over 100 ps with a 0.5 fs time step and strong Langevin coupling to relax the newly introduced side chain, and equilibrated for 1 ns in the

NPT ensemble at 310.15 K. Production dynamics were then run for 10 ns in the NPT ensemble at 310.15 K and 1 bar with a 2 fs time step, SHAKE on bonds to hydrogen, an 8 Å real-space cutoff, a Langevin thermostat (collision frequency 2 $ps^{-1}$) and a Berendsen barostat (relaxation time 2 ps), using `pmemd.cuda`. From the production trajectory the frame in which a single water molecule bridged the amine nucleophile and the catalytic sulfur selected to provide the reactive starting coordinates for the ML/MM stage.

**ML/MM metadynamics.** Reactive free-energy profiles for thioester aminolysis were computed with ML/MM metadynamics. The ML region, comprising the reacting fragments, the catalytic residues and the bridging water, was described by the MACEPOL-EF under electrostatic embedding, the ML–MM interaction range followed the 8 Å MM cutoff. Trajectories were propagated in the NPT ensemble at 310.15 K and 1.01325 bar with a Langevin thermostat (collision frequency 5 $ps^{-1}$) and a Berendsen barostat (relaxation time 2 ps) using a 0.5 fs time step. Each variant was run in one independent 100 ps replicate.

Free-energy sampling used the OPES-MetaD as implemented in PLUMED, biasing two collective variables on a single two-dimensional surface. CV1 was the antisymmetric substitution coordinate $d(\mathrm{N}\cdots\mathrm{C}) - d(\mathrm{C–S}_\gamma)$, built from the distance between the amine nitrogen and the thioester carbonyl carbon and the distance between that carbon and the cysteine sulfur, so that it changes sign as the tetrahedral intermediate collapses. CV2 was the mean of the two proton-relay distances through the bridging water, $\frac{1}{2}\left[d(\mathrm{O_w}\cdots\mathrm{H_N}) + d(\mathrm{H_w}\cdots\mathrm{S}_\gamma)\right]$, describing the concerted shuttle that deprotonates the nucleophile and protonates the leaving thiolate. Kernels were deposited every 100 MD steps (50 fs) at 310.15 K; initial kernel widths (`SIGMA`) of 0.06 Å were rescaled adaptively subject to a per-CV lower bound (`SIGMA_MIN`) of 0.024 Å, and the `BARRIER` parameter was 26 kcal $mol^{-1}$. A fixed set of one-sided harmonic upper walls kept the reacting fragments associated: 3.0 Å on the nitrogen–carbon distance (20 kcal $mol^{-1}$ $Å^{-2}$), 4.0 Å on the carbon–sulfur distance (50 kcal $mol^{-1}$ $Å^{-2}$) and 2.2 Å on each of the two proton-relay distances (5 kcal $mol^{-1}$ $Å^{-2}$); the wall bias was recorded separately and accounted for on reweighting. ML regions contained 135–205 atoms (median 170) with formal charges of $0$, $+1$ or $-1$. $\Delta G^{\ddagger}$ was extracted for every run from the one-dimensional projection of the STATE-reconstructed surface onto CV1, and the per-variant prediction was taken as the mean over its replicates. All predictions were frozen before comparison with experiment.

### Experimental Protocols and Results

Table S2: Protein and DNA sequence of aldehyde dehydrogenases, XylB and TpNOX used in this study.

| No. | Protein and DNA sequence |
|---|---|
| 1 | **Protein sequence:**<br>>PHBDD (NCBI accession ID: AAA75634.2) [*Pseudomonas putida*]<br>MSQRLAAYENMSLQLIAGEWRVGKAGRDLDVLDPFTQEKLLQIPLANREDLDEAYRSARQAQVAWAACGPSERAQVMLNAVRI<br>FDERRDEIIDWIIRESGSTRIKAQIEWGAARAITQESASLPSRVHGRILASDVPGKESRVYREPLGVIGIISPWNFPLHLTAR<br>SLAPALALGNACVIKPASDTPVTGGLLLAHIFEEAGLPKGVLSVVVGSGSEIGDAFVEHEVPGFISFTGSTQVGRNIGRIAAG<br>GEHLKHVALELGGNSPFVVLADADLDQAVNAAVVGKFLHQGQICMAINRIIVEDSVYDEFVNRYAERVKSLPYGDPSKPETVV<br>GPVINAKQLAGLQDKIATAKSEGARVMVEGEAQGNVLPPHVFADVTADMEIAREEIFGPLVGIQRARDEAHALELANSSEYGL<br>SSAVFTSSLERGVKFARGIRAGMTHINDIPVNDEPNAPFGGEKNSGLGRFNGDWAIEEFTTDHWITVQHAPRRYPF<br>**DNA sequence:**<br>>PHBDD<br>ATGTCACAGCGTCTGGCCGCCTATGAAAATATGTCTTTACAGCTGATTGCAGGCGAATGGCGCGTGGGTAAAGCCGGTCGTGA<br>TCTGGATGTGCTTGATCCGTTTACCCAGGAAAAACTGCTTCAGATTCCGCTGGCCAATCGCGAGGACCTGGATGAAGCCTATC<br>GTAGTGCACGTCAGGCACAGGTTGCATGGGCAGCCTGTGGTCCGAGCGAACGTGCACAGGTTATGCTGAATGCAGTTCGTATT<br>TTTGATGAACGTCGCGATGAAATTATTGATTGGATTATTCGTGAAAGTGGCTCTACCCGTATTAAAGCCCAGATTGAATGGGG<br>CGCCGCACGCGCCATTACCCAGGAATCAGCCTCTCTGCCGAGTCGTGTTCATGGTCGCATTCTGGCCTCAGATGTTCCGGGTA<br>AAGAATCTCGTGTGTATCGCGAACCGTTAGGTGTGATTGGCATTATTTCACCGTGGAATTTTCCGTTACATCTGACCGCCCGC<br>TCACTGGCCCCGGCCTTAGCCTTAGGCAATGCCTGTGTGATTAAACCGGCAAGTGATACCCCGGTGACCGGCGGCTTACTGTT<br>AGCACATATTTTTGAAGAAGCAGGCTTACCGAAAGGCGTGCTGTCAGTTGTTGTGGGTAGTGGTAGCGAAATTGGCGATGCCT<br>TTGTGGAACATGAAGTTCCGGGCTTTATTTCTTTTACCGGCTCAACCCAGGTGGGTCGCAATATTGGTCGCATTGCAGCAGGC<br>GGTGAACATCTGAAACATGTTGCCTTAGAACTGGGCGGTAATAGTCCGTTTGTGGTTTTAGCAGATGCCGATCTGGATCAGGC<br>AGTTAATGCAGCAGTTGTTGGCAAATTTCTGCATCAGGGCCAGATTTGTATGGCAATTAATCGTATTATTGTGGAAGATAGCG<br>TGTATGATGAATTTGTTAATCGCTATGCAGAACGTGTTAAATCACTGCCGTATGGTGATCCGAGTAAACCGGAAACCGTTGTG<br>GGTCCGGTTATTAATGCCAAACAGTTAGCCGGCTTACAGGATAAAATTGCCACCGCCAAATCAGAAGGTGCACGAGTTATGGT<br>TGAAGGCGAAGCACAGGGTAATGTGCTGCCGCCGCATGTGTTTGCCGATGTGACCGCCGATATGGAAATTGCACGCGAAGAAA<br>TTTTTGGTCCGTTAGTGGGCATTCAGCGCGCACGCGATGAAGCCCATGCCCTGGAACTGGCCAATAGCTCAGAATATGGTCTG<br>TCTAGCGCAGTGTTTACCTCCTCGTTAGAACGCGGTGTTAAATTTGCACGCGGCATTCGCGCAGGTATGACCCATATTAATGA<br>TATTCCGGTTAATGATGAACCGAATGCCCCGTTTGGCGGCGAAAAAAATAGCGGCTTAGGTCGCTTTAATGGTGATTGGGCCA<br>TTGAAGAATTTACCACCGATCATTGGATTACCGTTCAGCACGCCCCGCGTCGCTATCCGTTTTAA |
| 2 | **Protein sequence:**<br>>ALDH3 (NCBI accession ID: WP_192907353.1) [*Pseudomonas*]<br>MTQLAGKYAAFNQQFIAGVWRDGRDSTVLQVNNPFDGSLLTEIVQADREDLDEAYRQAAEAQKQWAATGPAERAAVLHRVVQI<br>FDQRREEIIDWIIRESGSTRLKAQIEWGAARGITLESASFPARVHGRILTSNIPGKESRVYREPLGVVGVISPWNFPLHLSQR<br>SVAPALALGNAVVIKPASDTPVTGGLLLARIYEEAGLPAGLLSVLVGSGAKIGDAFVDHEVPKFISFTGSTPVGLNIGRLASG<br>GKHLKHVALELGGNSPFVVLADADLEQAVNAAVMGKFLHQGQICMAINRIIVEDGLYEAFVTRYAQRVNTLKVGDPDDADTVI<br>GPIINARQLQGLVAKIDRAKEEGAKVVVQGDVVGQLLPPHVFADVTADMEIAQEEIFGPLVGIQRAVDEAHALELANHSEFGL<br>SSAVFTRDLERGVRFARQVKAGMTHVNDIPVNDEAHAPFGGEKNSGLGRFNGEWAIEEFTTDHWISVQHTPRHYPF<br>**DNA sequence:**<br>>ALDH3<br>ATGACCCAGCTGGCGGGCAAATACGCCGCGTTCAACCAGCAGTTCATTGCGGGTGTGTGGCGCGATGGCCGCGATAGCACCGT<br>TCTGCAGGTGAATAACCCGTTTGATGGCTCCCTGCTGACCGAAATTGTGCAGGCGGATCGTGAAGATCTGGATGAAGCGTATC<br>GTCAGGCGGCCGAAGCCCAGAAACAGTGGGCCGCGACCGGCCCGGCGGAACGCGCCGCGGTACTGCACCGTGTGGTGCAGATT<br>TTTGATCAGCGCCGTGAAGAAATTATCGATTGGATTATTCGTGAAAGTGGCAGCACCCGCCTGAAAGCGCAGATTGAATGGGG<br>CGCGGCACGTGGCATTACCCTGGAAAGCGCAAGCTTTCCGGCCCGCGTTCACGGCCGTATTCTGACCAGTAATATTCCGGGCA<br>AAGAAAGCCGCGTGTATCGCGAACCGCTGGGCGTGGTGGGCGTGATTAGCCCGTGGAACTTTCCGCTGCATCTGAGCCAGCGC<br>AGCGTTGCGCCGGCCCTGGCGCTGGGCAACGCCGTGGTGATCAAACCGGCGAGCGATACCCCGGTGACCGGCGGCCTGCTGCT<br>GGCCCGCATTTACGAAGAAGCGGGCCTGCCGGCGGGCTTACTGAGCGTGCTGGTTGGCAGCGGCGCGAAAATTGGCGATGCCT<br>TTGTGGATCATGAAGTGCCGAAATTTATTAGCTTTACCGGCAGCACCCCGGTGGGCCTGAACATTGGTCGCCTGGCCAGCGGC<br>GGCAAACATCTGAAACATGTGGCCCTGGAACTGGGCGGCAACAGCCCGTTTGTGGTGCTGGCCGATGCGGATCTGGAACAGGC<br>AGTGAATGCGGCCGTGATGGGCAAATTCCTGCATCAGGGCCAGATTTGTATGGCGATTAACCGCATTATTGTGGAAGATGGCC<br>TGTACGAAGCGTTCGTGACCCGTTACGCGCAGCGCGTGAACACCCTGAAAGTTGGCGATCCGGATGATGCCGACACCGTGATT<br>GGTCCGATCATCAACGCCCGCCAACTGCAGGGCCTGGTGGCCAAAATTGATCGCGCGAAAGAAGAAGGGGCGAAAGTCGTGGT<br>TCAGGGCGATGTTGTTGGCCAGCTGCTGCCGCCGCATGTGTTCGCGGATGTGACCGCGGACATGGAAATCGCGCAGGAAGAAA<br>TTTTTGGCCCGCTGGTGGGCATTCAGCGCGCGGTGGATGAAGCGCATGCCCTGGAACTGGCCAACCACAGCGAGTTTGGCCTG<br>AGCTCGGCGGTGTTCACCCGCGATCTGGAACGTGGCGTGCGCTTTGCCCGCCAGGTGAAAGCGGGCATGACCCACGTGAACGA<br>CATTCCGGTGAACGATGAAGCACATGCACCGTTTGGTGGCGAAAAAAATAGCGGCCTGGGCCGTTTTAATGGCGAATGGGCGA<br>TCGAAGAATTTACCACCGATCACTGGATCTCCGTTCAGCATACCCCGCGTCATTACCCGTTCTAG |

*continued on next page*

**Table S2** (continued)

| No. | Protein and DNA sequence |
|---|---|
| 3 | **Protein sequence:**<br>>ConPHBDD (Consensus sequences obtained using Consensus)<br>MSQTMAAYQDFDLQFIAGEWREGSSGRTLEDTNPYTGETLLEIPLASREDLDEAYRAAAEAQKEWAATLPAERAAVLLRAAEI<br>MDERREEIIDWLIRESGSTRIKAEIEWGAARAITLEAASFPYRVHGRILPSDIPGKENRVYRKPLGVVGVISPWNFPLHLSMR<br>SVAPALALGNAVVLKPASDTPVTGGLLLAKIFEEAGLPAGVLNVVVGAGSEIGDAFVEHPVPRLISFTGSTPVGRHIGELAGG<br>GPHLKRVALELGGNNPFVVLDDADLDQAVEAAVFGKFLHQGQICMAINRIIVDASVYDEFVERFVERVKALKVGDPSDPDTVI<br>GPIINQKQLEGLQEKIARAREEGARLLLGGEPEGNVLPPHVFTDVTNDMEIAREEIFGPVAPIIRARDEEHALELANDTEYGL<br>SSAVFTRDLERGVRFARRIEAGMTHVNDQPVNDEPNVPFGGEKNSGLGRFNGEWAIEEFTTDHWISVQHEPRQYPF<br>**DNA sequence:**<br>>ConPHBDD<br>ATGAGCCAGACCATGGCAGCGTACCAGGATTTTGATCTGCAGTTTATTGCGGGCGAATGGCGCGAAGGCAGCAGCGGCCGTAC<br>CCTGGAAGATACGAACCCGTATACCGGCGAAACCCTGCTGGAAATCCCGCTGGCGAGCCGCGAAGATCTGGATGAAGCCTATC<br>GCGCGGCGGCCGAAGCACAGAAAGAATGGGCCGCCACCCTGCCGGCGGAACGCGCCGCGGTGCTGCTGCGCGCGGCGGAAATT<br>ATGGATGAACGCCGCGAAGAAATTATTGATTGGCTGATTCGTGAATCAGGTAGCACCCGTATTAAAGCGGAAATCGAATGGGG<br>CGCCGCACGCGCGATTACGCTGGAAGCGGCCAGTTTTCCGTATCGCGTTCATGGCCGCATTCTGCCGAGTGATATTCCGGGCA<br>AAGAAAACCGCGTATACCGTAAACCGCTGGGTGTGGTGGGTGTTATTAGCCCGTGGAATTTCCCGCTGCACCTGAGCATGCGC<br>AGCGTGGCACCGGCCCTGGCGCTGGGTAACGCGGTGGTGCTGAAACCGGCGAGCGATACCCCGGTGACCGGCGGCCTGCTGCT<br>GGCAAAAATTTTTGAAGAAGCGGGCCTGCCGGCGGGCGTGCTGAATGTTGTGGTGGGCGCGGGCAGCGAGATTGGTGATGCGT<br>TTGTTGAACATCCGGTGCCGCGCCTGATTAGTTTCACCGGCAGCACCCCGGTGGGCCGCCATATTGGCGAACTGGCGGGCGGC<br>GGCCCGCATCTGAAACGTGTTGCCCTGGAGCTGGGCGGCAACAACCCGTTTGTGGTGCTGGACGATGCGGATCTGGACCAGGC<br>GGTTGAAGCGGCGGTTTTTGGTAAATTCCTGCACCAGGGTCAGATTTGCATGGCCATTAACCGCATTATTGTGGATGCCAGCG<br>TGTACGATGAATTTGTGGAACGTTTTGTTGAACGCGTGAAAGCGCTGAAAGTGGGCGATCCGTCGGATCCGGATACCGTGATT<br>GGCCCGATTATTAACCAGAAACAGCTGGAAGGTCTGCAGGAAAAAATTGCGCGCGCACGCGAAGAAGGCGCGCGCCTGCTGCT<br>GGGCGGCGAACCGGAAGGCAACGTGCTGCCGCCGCATGTCTTCACCGATGTGACCAACGACATGGAAATTGCGCGCGAAGAAA<br>TTTTTGGTCCGGTGGCCCCGATCATTCGCGCCCGTGATGAAGAACATGCGCTGGAACTGGCGAACGACACCGAATACGGCCTG<br>TCAAGTGCGGTGTTCACCCGCGATCTGGAACGCGGCGTGCGCTTTGCGCGCCGCATTGAAGCGGGCATGACCCATGTGAACGA<br>TCAGCCGGTGAACGATGAGCCGAATGTACCGTTTGGCGGCGAAAAAAATAGCGGCCTGGGCCGCTTTAACGGCGAATGGGCCA<br>TTGAAGAATTTACCACCGATCATTGGATTAGCGTGCAGCATGAACCGCGCCAGTACCCGTTTTAG |
| 4 | **Protein sequence:**<br>>TpNOX (*Lb*NOX, NCBI accession ID: BAN07126.1) [*Lactobacillus brevis* KB290]<br>MKVTVVGCTHAGTFAIKQILAEHPDAEVTVYERNDVISFLSCGIALYLGGKVADPQGLFYSSPEELQKLGANVQMNHNVLAID<br>PDQKTVTVEDLTNHAQTTESYDKLVMTSGSWPIVPKIPGIDSDRVKLCKNWAHAQALIEDAKEAKRITVIGAGYIAAELAEAY<br>STTGHDVTLIARSARVMRKYFDADFTDVIEQDYRDHGVQLALGETVESFTDSATGLTIKTDKNSYETDLAILCIGFRPNTDLL<br>KGKVDMAPNGAIITDDYMRSSNPDIFAAGDSAAVHYNPTHQNAYIPLATNAVRQGILVGKNLVKPTVKYMGTQSSSGLALYDR<br>TIVSTGLTLAAAKQQGLNAEQVIVEDNYRPEFMPSTEPVLMSLVFDPDTHRILGGALMSKYDVSQSANTLSVCIQNENTIDDL<br>AMVDMLFQPNFDRPFNYLNILAQAAQAKVAQSVNA<br>**DNA sequence:**<br>>TpNOX<br>ATGAAGGTGACCGTTGTTGGCTGCACCCATGCAGGTACCTTTGCCATTAAACAGATTCTGGCCGAACATCCGGATGCCGAAGT<br>TACCGTGTATGAACGTAATGATGTTATCAGCTTCCTGAGTTGTGGCATTGCACTGTATCTGGGCGGCAAAGTGGCAGATCCGC<br>AGGGTTTATTTTATAGTAGCCCGGAAGAACTGCAGAAACTGGGCGCAAATGTTCAGATGAATCATAATGTTCTGGCGATTGAT<br>CCGGATCAGAAAACCGTTACCGTGGAAGATCTGACCAATCATGCCCAGACCACCGAAAGCTATGATAAACTGGTTATGACCAG<br>TGGCAGTTGGCCGATTGTGCCGAAAATTCCGGGCATTGATAGTGATCGCGTGAAACTGTGCAAAAATTGGGCCCATGCCCAGG<br>CCCTGATTGAAGATGCAAAAGAAGCCAAACGCATTACCGTGATTGGTGCCGGTTATATTGCAGCAGAACTGGCAGAAGCCTAT<br>AGCACCACCGGTCATGATGTTACCCTGATTGCACGTAGCGCCCGCGTGATGCGTAAATATTTTGATGCAGATTTCACCGACGT<br>GATTGAACAGGATTATCGTGATCATGGTGTTCAGCTGGCACTGGGCGAAACCGTGGAAAGTTTTACCGATAGCGCCACCGGTC<br>TGACCATTAAAACCGATAAAAATAGCTACGAGACCGATCTGGCCATTCTGTGCATTGGCTTTCGCCCGAATACCGATCTGCTG<br>AAAGGTAAAGTGGATATGGCACCGAATGGTGCAATTATTACCGATGATTATATGCGTAGCAGCAATCCGGATATTTTTGCAGC<br>CGGTGATAGCGCCGCAGTTCATTATAATCCGACCCATCAGAATGCCTATATTCCGCTGGCAACCAATGCAGTGCGTCAGGGTA<br>TTCTGGTTGGTAAAAATCTGGTGAAACCGACCGTGAAATATATGGGTACCCAGAGCAGCAGCGGCCTGGCATTATATGATCGC<br>ACCATTGTGAGCACCGGTCTGACACTGGCAGCAGCTAAACAGCAGGGTCTGAATGCAGAACAGGTTATTGTGGAAGATAACTA<br>TCGTCCGGAATTTATGCCGAGCACCGAACCGGTTCTGATGAGCCTGGTGTTTGATCCGGATACCCATCGCATTCTGGGTGGCG<br>CACTGATGAGCAAATATGATGTTAGCCAGAGTGCAAATACCCTGAGCGTGTGCATTCAGAATGAAAATACCATTGACGACCTG<br>GCAATGGTTGATATGCTGTTTCAGCCGAATTTTGATCGCCCGTTTAATTATCTGAACATCCTGGCCCAGGCCGCACAGGCAAA<br>AGTTGCTCAGAGCGTGAATGCCTAA |

Table S3: Primer sequence used in this study.

| Primer | Sequences |
|---|---|
| pQlinkHx-F | GCGGCCGCCTAGGACCCAGCTTTCTTGTACAAAG |
| pQlinkHx-R | GGATCCCTGAAAATACAGGTTTTC |
| pQlinkHx-PHBDD-F | GAAAACCTGTATTTTCAGGGATCCATGTCACAGCGTCTGGCCGCCTATG |
| pQlinkHx-PHBDD-R | CAAGAAAGCTGGGTCCTAGGCGGCCGCTTAAAACGGATAGCGACGCGGGGCGTGCTG |
| pQlinkHx-ALDH3-F | GAAAACCTGTATTTTCAGGGATCCATGACCCAGCTGGCGGGCAAATACG |
| pQlinkHx-ALDH3-R | CTTTGTACAAGAAAGCTGGGTCCTAGGCGGCCGCCTAGAACGGGTAATGACGCGGGGT ATGC |
| pQlinkMx-TpNOX-F | GAAAACCTGTATTTTCAGGGATCCatgaaggtgaccgttgttggctgc |
| pQlinkMx-TpNOX-R | CTTTGTACAAGAAAGCTGGGTCCTAGGCGGCCGCttaggcattcacgctctgagcaac ttttgcc |
| pQlinkHx-Down-R | CCACTCCATTTGTAACCACTCC |
| PHBDD-R166-F | GAATTTTCCGTTACATCTGACCGCCNNNTCAC |
| PHBDD-E259-R | CTATTACCGCCCAGNNNTAAGGCAAC |
| PHBDD-P156V-F | GGCATTATTTCAGTTTGGAATTTTCCG |
| PHBDD-L163G-F | TTCCGTTACATggcACCGCC |
| PHBDD-R166M-F | ACCGCCatgTCACTGGC |
| PHBDD-R166Q-F | ACCGCCcagTCACTGGC |
| PHBDD-T234A-F | TTCTTTTgcgGGCTCAACCCAGG |
| PHBDD-T234G-F | TTCTTTTggcGGCTCAACCCAGG |
| PHBDD-T234H-F | TTCTTTTcatGGCTCAACCCAGG |
| PHBDD-E259L-F | GTTGCCTTActgCTGGGCGG |
| ALDH3-L163G-F | TCCGCTGCATggcAGCCAG |
| ALDH3-R166M-F | AGCCAGatgAGCGTTGC |
| ALDH3-R166Q-F | AGCCAGcagAGCGTTGC |
| ALDH3-T234G-F | ATTAGCTTTggcGGCAGCACC |
| ALDH3-T234H-F | ATTAGCTTTcatGGCAGCACC |
| ALDH3-E259L-F | GGCCCTGctgCTGGGCG |
| conPHB-F | ATGAGCCAGACCATGGCAGC |
| conPHB-R | CTAAAACGGGTACTGGCGCG |
| conPHB-L163G-F | TCCCGCTGCACGGTAGCATG |
| conPHB-R166M-F | AGCATGATGAGCGTGGCA |
| conPHB-R166Q-F | AGCATGCAGAGCGTGGCA |
| conPHB-T234G-F | GATTAGTTTCGGTGGCAGCACC |
| conPHB-T234H-F | GATTAGTTTCCATGGCAGCACC |
| conPHB-E259L-F | TTGCCCTGCTGCTGGGCG |

### Chemo-enzymatic synthesis of ABBV-318 (34)

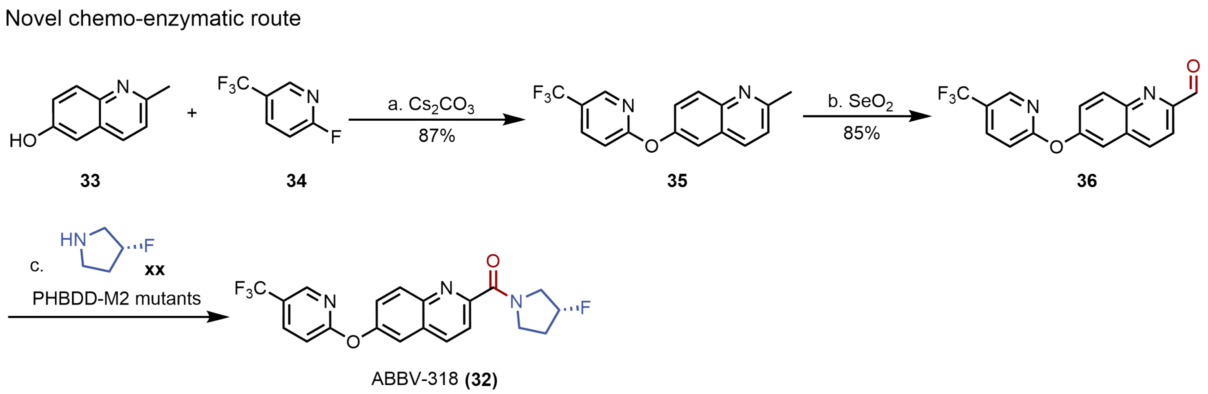


Figure S1: **Chemo-enzymatic route to ABBV-318 (34).** Nucleophilic aromatic substitution of 2-methylquinolin-6-ol (**35**) with 2-fluoro-5-(trifluoromethyl)pyridine (**36**) gives the diaryl ether **37**; benzylic oxidation with $SeO_2$ delivers the aldehyde **38**; and enzymatic oxidative amidation with (*R*)-3-fluoropyrrolidine (**33**), catalysed by whole *E. coli* cells overexpressing **M2** (**M1**-L163G-T234G), furnishes ABBV-318 (**34**).

### Compound 37:

#### 2-Methyl-6-((5-(trifluoromethyl)pyridin-2-yl)oxy)quinoline

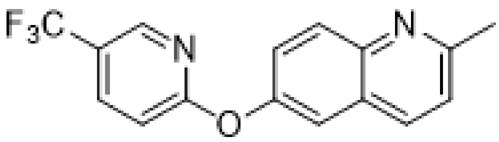

To a solution of 2-methylquinolin-6-ol (**35**; 160 mg, 1.0 mmol, 1.0 equiv.) in DMF (1 mL) were added 2-fluoro-5-(trifluoromethyl)pyridine (**36**; 215 mg, 1.3 mmol, 1.3 equiv.) and $Cs_2CO_3$ (490 mg, 1.5 mmol, 1.5 equiv.) under an argon atmosphere. The reaction was stirred at 80 °C until complete consumption of the starting material. The mixture was filtered and concentrated under reduced pressure, and the crude residue was purified by flash column chromatography (petroleum ether/EtOAc) to afford **37** as a white solid (266 mg, 87%).

**$^1$H NMR (400 MHz, acetone-$d_6$)** $\delta$ 8.48 (dt, $J$ = 2.7, 1.0 Hz, 1H), 8.22–8.15 (m, 2H), 8.01 (d, $J$ = 9.1 Hz, 1H), 7.70 (d, $J$ = 2.7 Hz, 1H), 7.57 (dd, $J$ = 9.1, 2.7 Hz, 1H), 7.42 (d, $J$ = 8.4 Hz, 1H), 7.29 (dt, $J$ = 8.7, 0.8 Hz, 1H), 2.69 (s, 3H).

**$^{13}$C NMR (101 MHz, acetone-$d_6$)** $\delta$ 166.87, 159.48, 151.49, 146.65, 146.11 (q, $J$ = 4.4 Hz), 138.15 (q, $J$ = 3.4 Hz), 136.47, 131.12, 127.97, 125.63, 124.96 (q, $J$ = 271.7 Hz), 123.33,

122.19 (q, $J$ = 32.9 Hz), 118.75, 112.78, 25.18.

**$^{19}$F NMR (471 MHz, acetone-$d_6$)** $\delta$ −62.13.

**HRMS (ESI)** $m/z$: $[M + H]^+$ calcd for $C_{16}H_{12}F_3N_2O^+$ 305.0896; found 305.0895.

**TLC** $R_f$ = 0.4 (petroleum ether/EtOAc 2:1).

#### Compound 38:

#### 6-((5-(Trifluoromethyl)pyridin-2-yl)oxy)quinoline-2-carbaldehyde

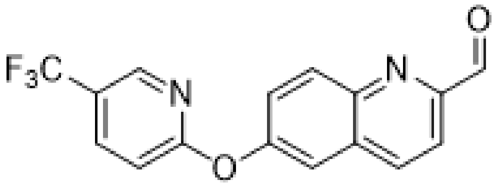

To a solution of **37** (305 mg, 1.0 mmol, 1.0 equiv.) in 1,4-dioxane (1 mL) was added $SeO_2$ (222 mg, 2.0 mmol, 2.0 equiv.) under an argon atmosphere. The reaction mixture was stirred at 80 °C until complete consumption of the starting material. The mixture was filtered and concentrated under reduced pressure, and the crude residue was purified by flash column chromatography (petroleum ether/EtOAc) to afford **38** as a white solid (270 mg, 85%).

**$^1$H NMR (400 MHz, acetone-$d_6$)** $\delta$ 10.16 (d, $J$ = 0.9 Hz, 1H), 8.59–8.48 (m, 2H), 8.32 (d, $J$ = 9.1 Hz, 1H), 8.25 (dd, $J$ = 8.7, 2.6 Hz, 1H), 8.03 (d, $J$ = 8.5 Hz, 1H), 7.91 (d, $J$ = 2.6 Hz, 1H), 7.80 (dd, $J$ = 9.2, 2.7 Hz, 1H), 7.38 (dt, $J$ = 8.7, 0.8 Hz, 1H).

**$^{13}$C NMR (101 MHz, acetone-$d_6$)** $\delta$ 194.07, 166.41, 154.42, 153.47, 146.56, 146.16 (q, $J$ = 4.5 Hz), 138.43 (q, $J$ = 3.0 Hz), 138.09, 132.85, 131.89, 127.26, 124.92 (q, $J$ = 271.7 Hz), 122.72 (q, $J$ = 32.9 Hz), 118.87, 118.27, 113.23.

**$^{19}$F NMR (471 MHz, acetone-$d_6$)** $\delta$ −62.20.

**HRMS (ESI)** $m/z$: $[M + H]^+$ calcd for $C_{16}H_{10}F_3N_2O_2^+$ 319.0689; found 319.0681.

**TLC** $R_f$ = 0.75 (petroleum ether/EtOAc 2:1).

#### Compound 34: (*R*)-(3-Fluoropyrrolidin-1-yl)(6-((5-(trifluoromethyl)pyridin-2-yl)oxy)quinolin-2-yl)methanone (ABBV-318)

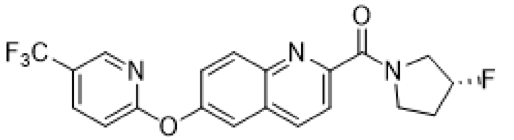


*E. coli* cells overexpressing **M2**-G163C (2.0 g) were suspended in CAPS buffer (10 mL, 0.1 M, pH 11). $NADP^+$ sodium salt (230 mg, 0.30 mmol, 1.5 equiv.), (*R*)-3-fluoropyrrolidine hydrochloride (**33**; 252 mg, 1.0 mmol, 5 equiv.), NaOH (160 mg, 1.0 mmol, 5 equiv., dissolved in 1.0 mL $H_2O$) and **38** (63.6 mg, 0.2 mmol, 1.0 equiv., dissolved in 1.0 mL DMSO) were added sequentially to the suspension. The reaction mixture was incubated at 37 °C and 220 rpm for 6 h. Saturated aqueous NaCl (5 mL) and EtOAc (20 mL) were added and the mixture was shaken vigorously to lyse the cells. The mixture was centrifuged (3,000 × *g*, 15 min) and the aqueous layer was extracted with EtOAc (2 × 20 mL). The combined organic layers were washed with saturated aqueous NaCl (10 mL), dried over anhydrous $Na_2SO_4$ and concentrated under reduced pressure. The crude residue was purified by flash column chromatography (petroleum ether/EtOAc) to afford **34** as a white solid (66.9 mg, 82%).

*Data are reported for a pair of amide rotamers.*

**$^1$H NMR (500 MHz, DMSO-$d_6$)** $\delta$ 8.58 (d, *J* = 2.5 Hz, 1H), 8.47 (d, *J* = 8.6 Hz, 1H), 8.28 (dd, *J* = 8.7, 2.6 Hz, 1H), 8.15 (dd, *J* = 9.1, 6.0 Hz, 1H), 7.95–7.83 (m, 2H), 7.71 (dd, *J* = 9.0, 2.7 Hz, 1H), 7.38 (d, *J* = 8.7 Hz, 1H), 5.40 (ddt, *J* = 54.0, 14.7, 3.5 Hz, 1H), 4.17–4.04 (m, 1H), 4.04–3.92 (m, 1H), 3.92–3.84 (m, 1H), 3.84–3.59 (m, 1H), 2.28–2.04 (m, 2H).

**$^{13}$C NMR (126 MHz, DMSO-$d_6$)** $\delta$ 165.53 (d, *J* = 25.2 Hz), 165.29, 153.12, 152.96, 151.81, 151.78, 145.29 (q, *J* = 4.4 Hz), 143.61, 143.54, 137.81 (q, *J* = 3.2 Hz), 136.82, 136.77, 131.08, 131.06, 128.70, 128.67, 125.90, 123.81 (q, *J* = 272.2 Hz), 121.05, 120.98, 120.85 (q, *J* = 32.8 Hz), 93.70 (d, *J* = 173.2 Hz), 91.65 (d, *J* = 172.1 Hz), 55.08 (d, *J* = 22.1 Hz), 53.23 (d, *J* = 22.3 Hz), 46.30, 44.49, 32.54 (d, *J* = 20.8 Hz), 29.93 (d, *J* = 21.0 Hz).

**$^{19}$F NMR (471 MHz, DMSO-$d_6$)** $\delta$ −60.18, −176.51, −176.70.

**HRMS (ESI)** *m/z*: $[M + H]^+$ calcd for $C_{20}H_{16}F_4N_3O_2^+$ 406.1173; found 406.1167.

**TLC** $R_f$ = 0.1 (petroleum ether/EtOAc 2:1).

### NMR spectra

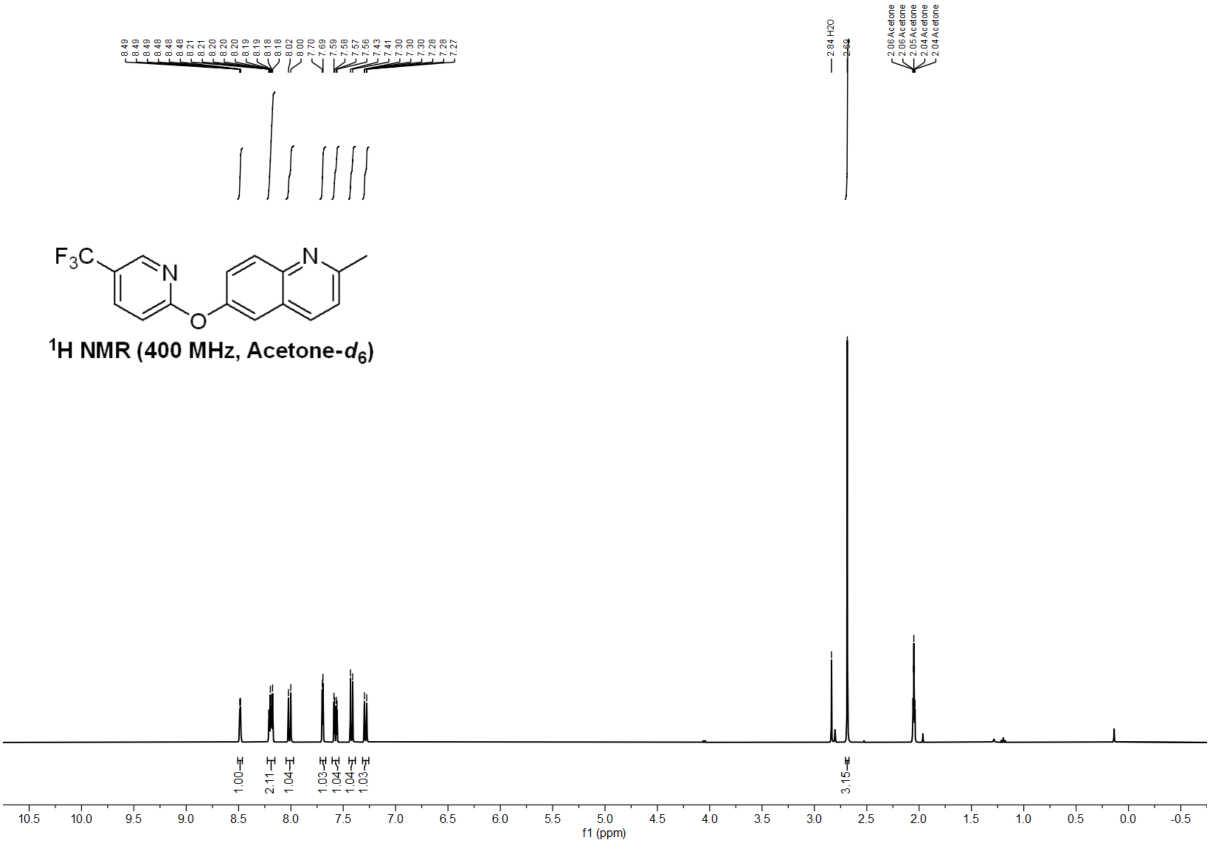


Figure S2: $^1$H NMR spectrum (400 MHz, acetone-$d_6$) of compound **37**.

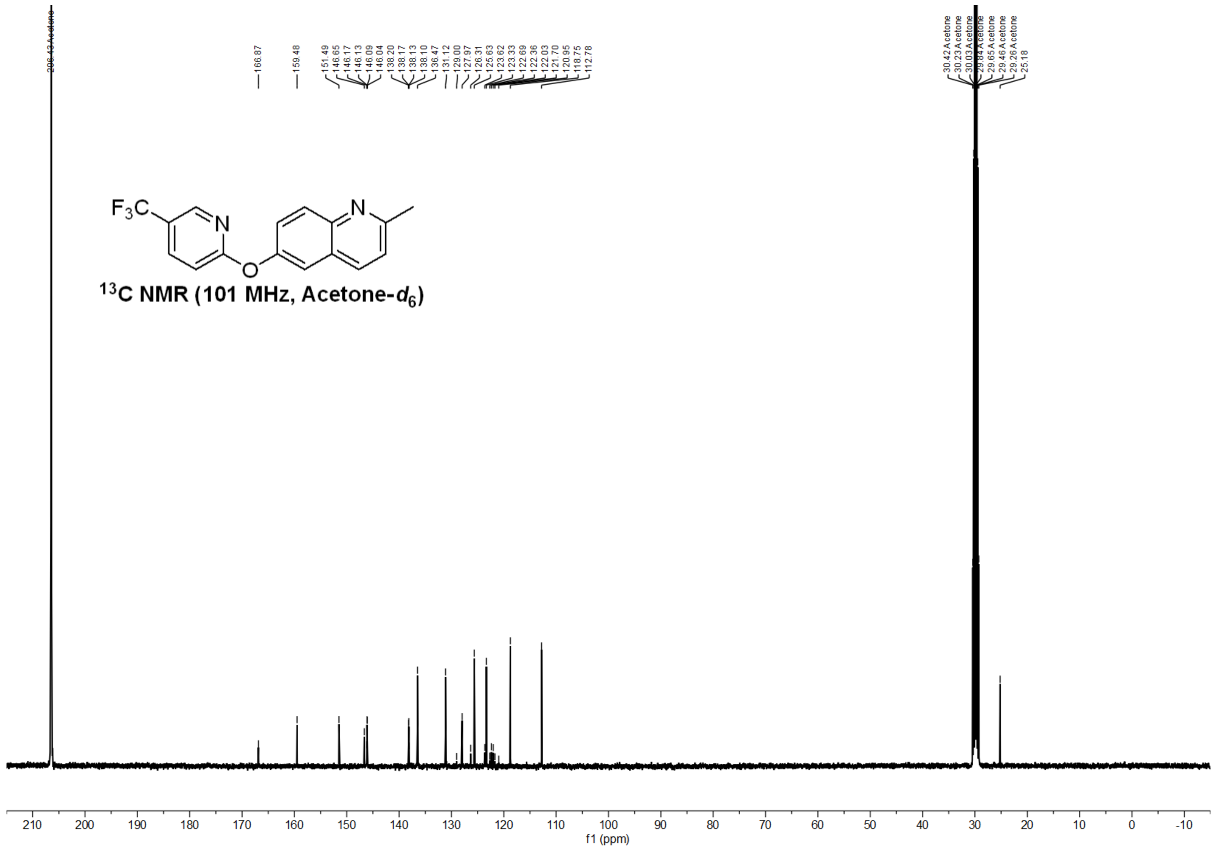

Figure S3: $^{13}C$ NMR spectrum (101 MHz, acetone-$d_6$) of compound **37**.

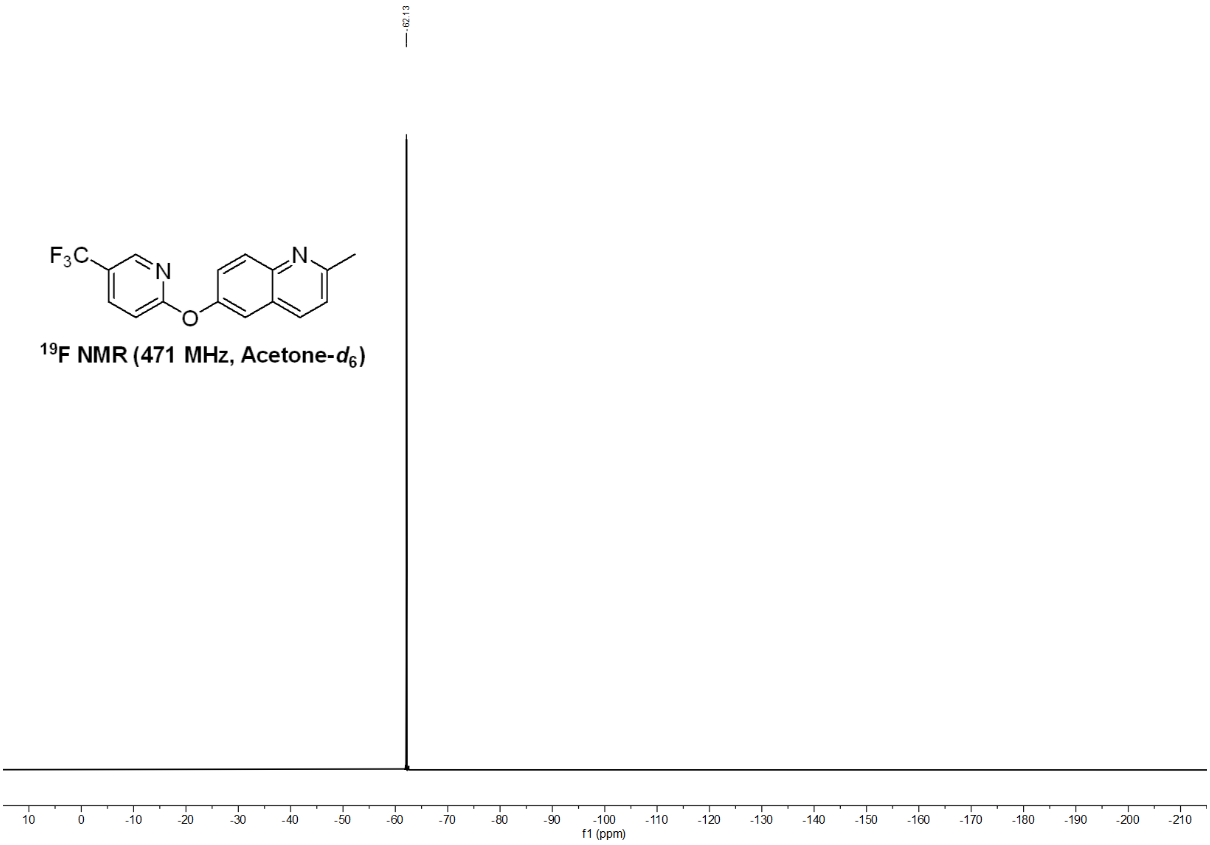


Figure S4: $^{19}F$ NMR spectrum (471 MHz, acetone-$d_6$) of compound **37**.

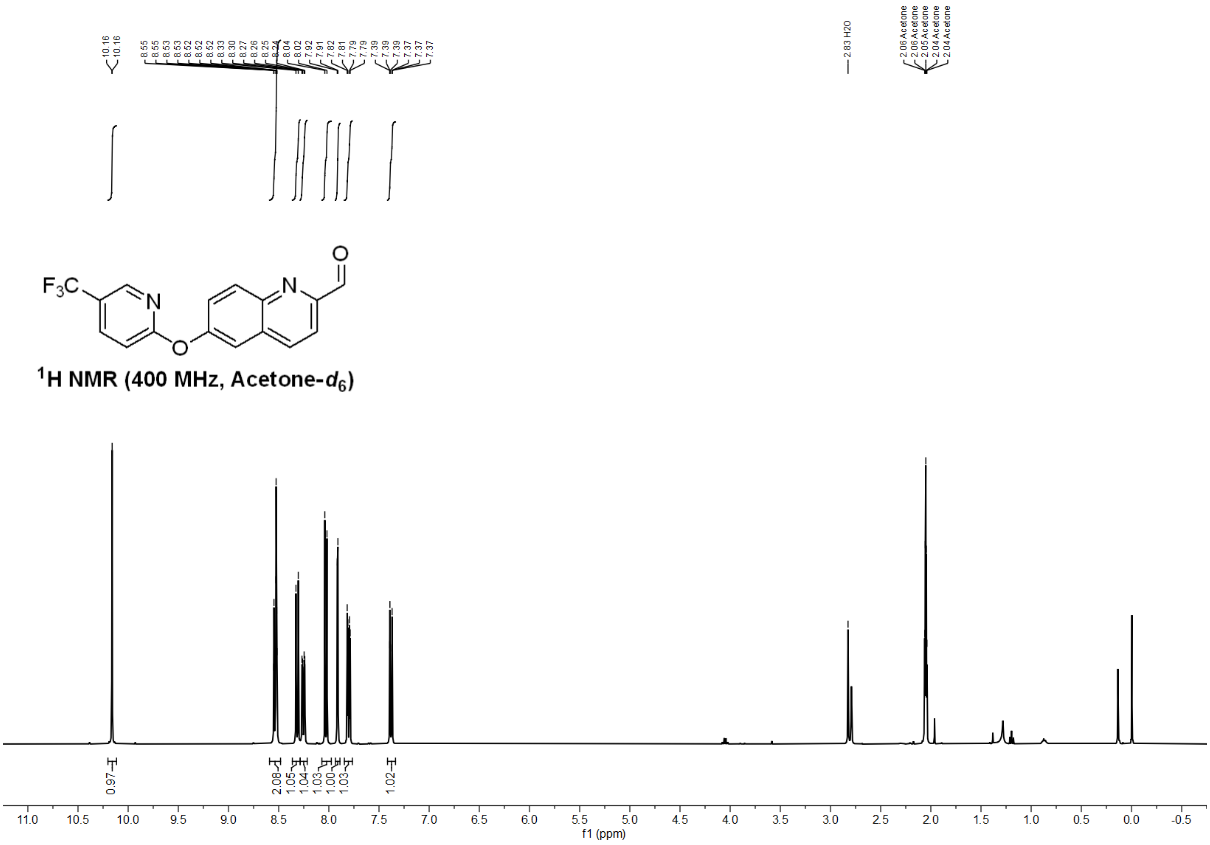


Figure S5: $^1$H NMR spectrum (400 MHz, acetone-$d_6$) of compound **38**.

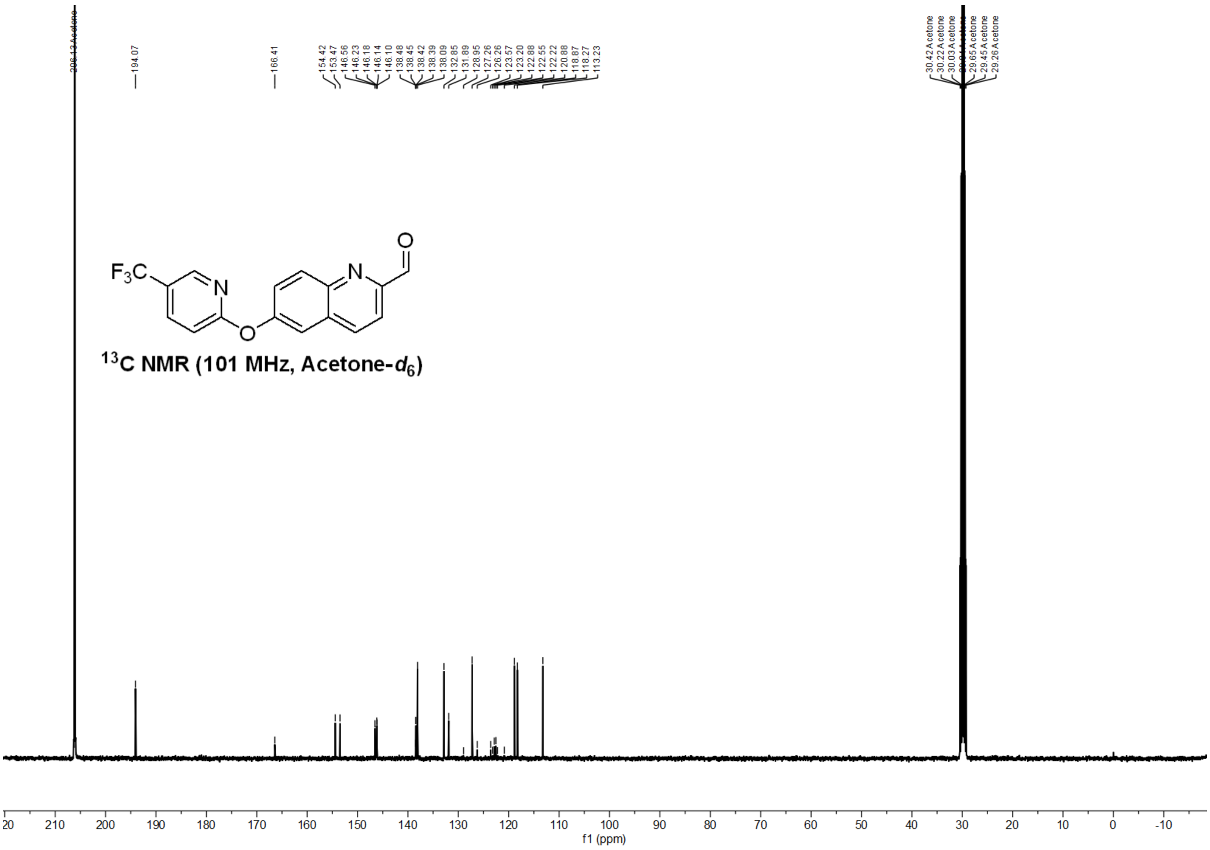


Figure S6: $^{13}$C NMR spectrum (101 MHz, acetone-$d_6$) of compound **38**.

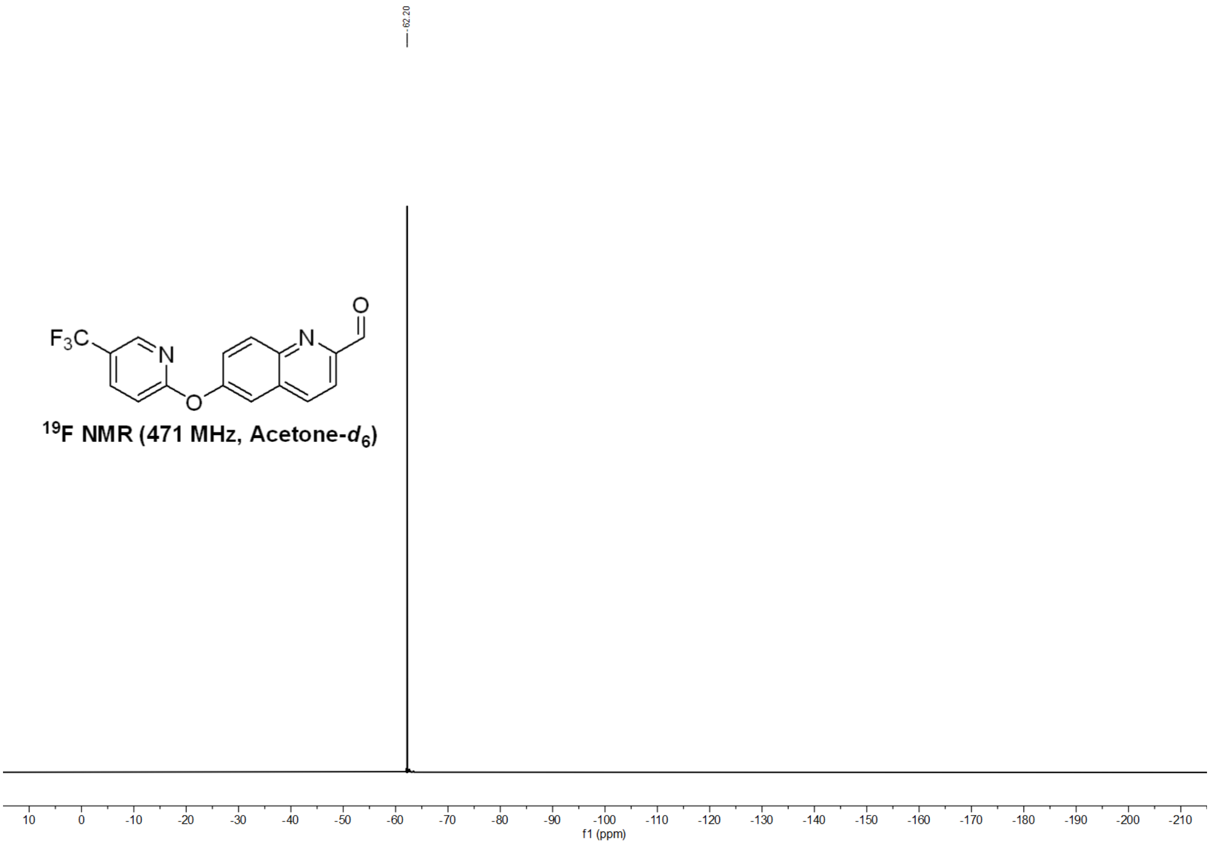


Figure S7: $^{19}F$ NMR spectrum (471 MHz, acetone-$d_6$) of compound **38**.

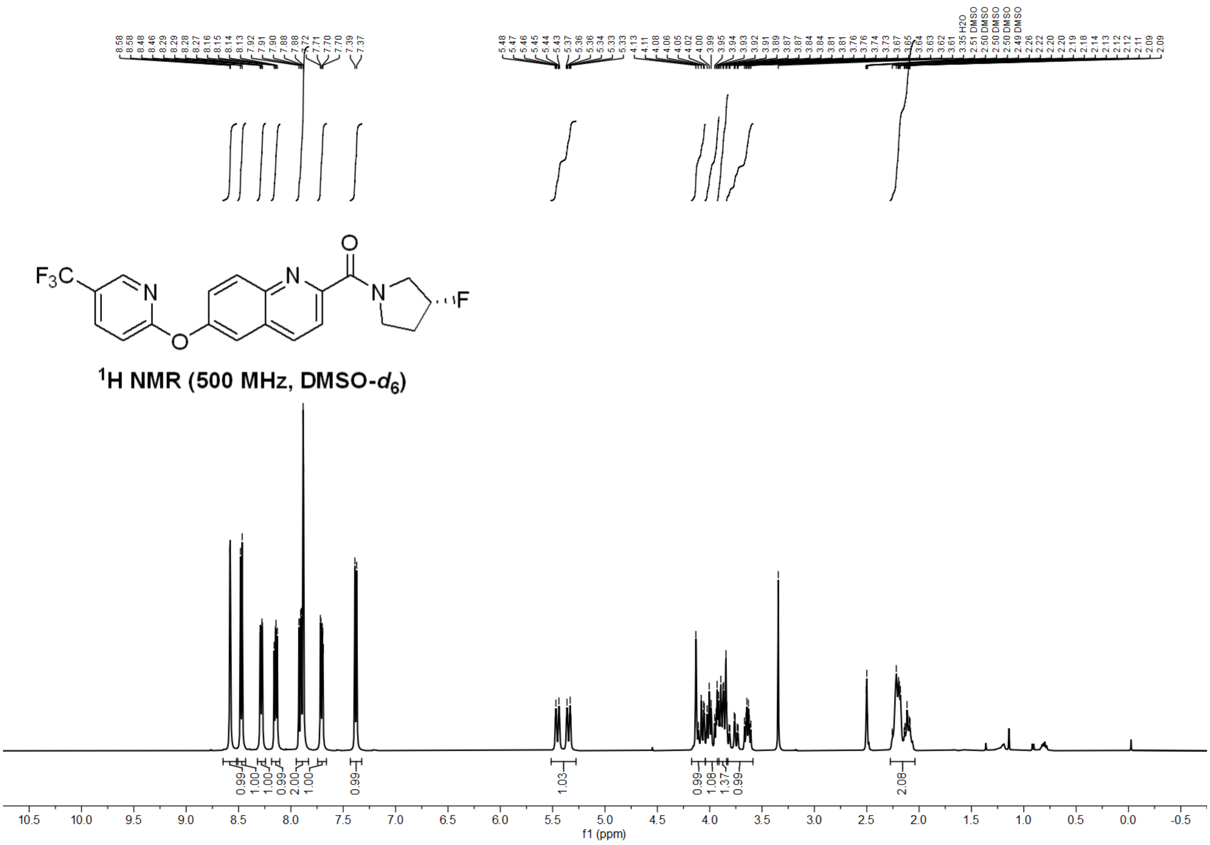

Figure S8: $^{1}$H NMR spectrum (500 MHz, DMSO-$d_6$) of ABBV-318 (**34**).

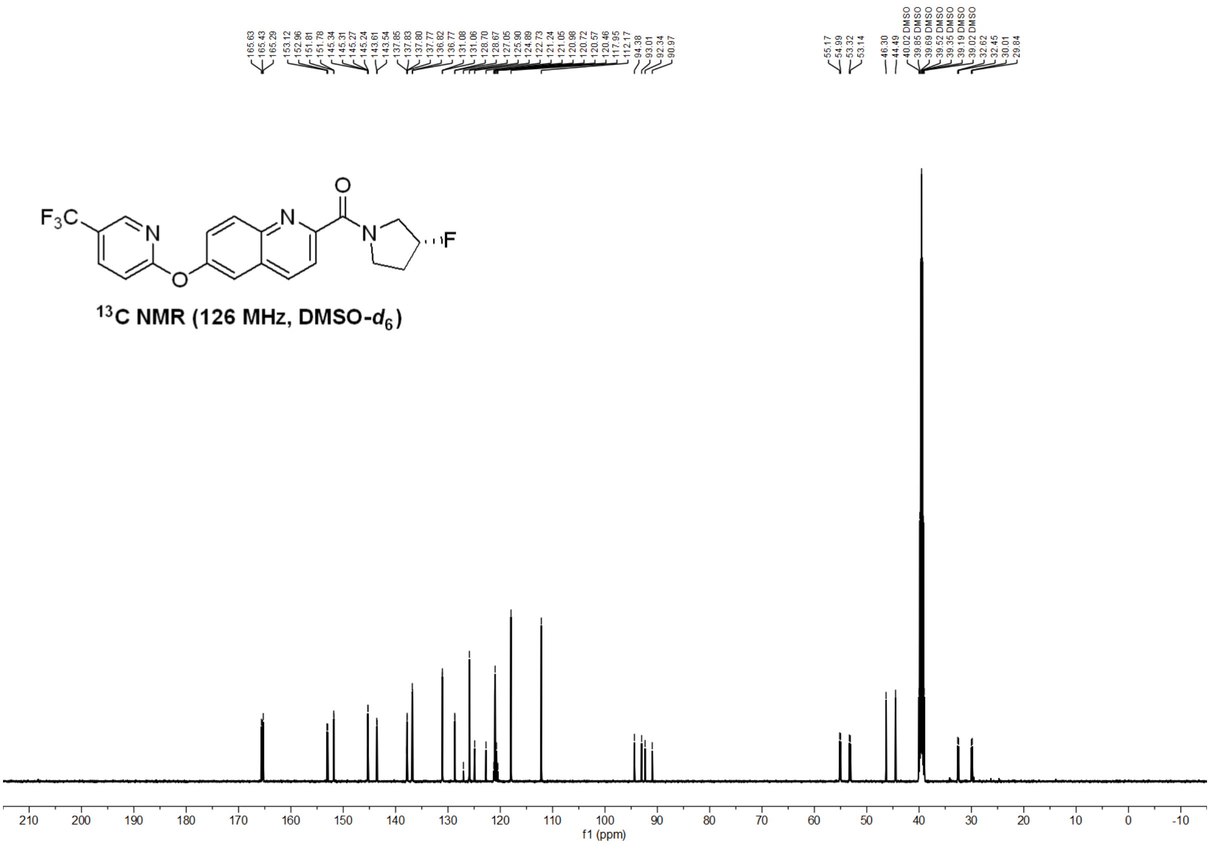

Figure S9: $^{13}$C NMR spectrum (126 MHz, DMSO-$d_6$) of ABBV-318 (**34**).

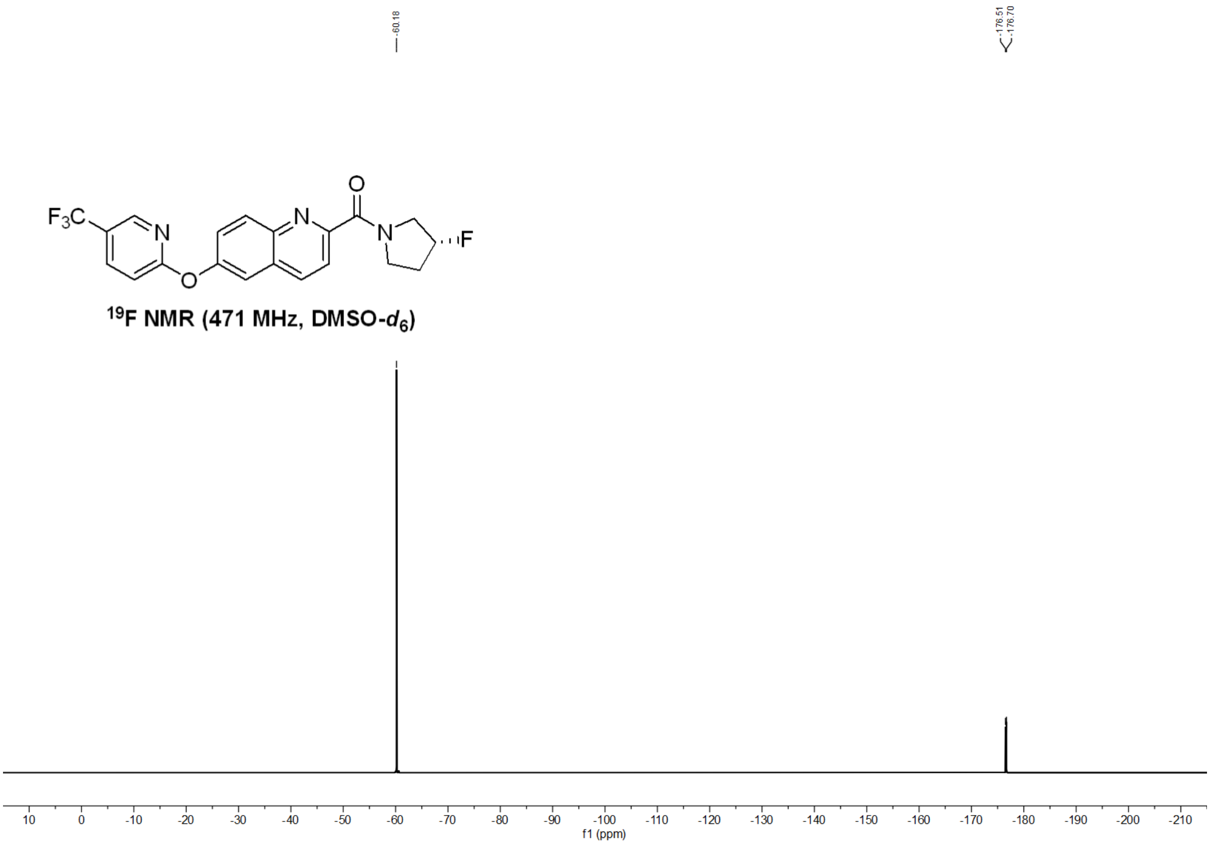


Figure S10: $^{19}$F NMR spectrum (471 MHz, DMSO-$d_6$) of ABBV-318 (**34**).
